# Experimental approach to assessing the impact of the intensity of near-natural heatwaves on three marine phanerogam species in Mediterranean lagoon environment

**DOI:** 10.1101/2025.03.26.645255

**Authors:** Constance Bourdier, Rutger De Wit, Laura Soissons, Elisa De Ronne, Vincent Ouisse

## Abstract

The increased intensity, frequency and duration of extreme climatic events (ECEs) in the context of climate change have been recognised as additional pressures for seagrass habitat in lagoon ecosystems. Among these ECE, this study focuses on the impact of marine heatwaves (MHWs) on seagrass species in Mediterranean coastal lagoons, *i.e.,* the seagrasses *Cymodocea nodosa, Nanozostera noltei,* and *Ruppia cirrhosa*. We experimentally assessed with a near-natural mesocosm, the effects of a 20-day heatwave at three intensities (+2, +4, +6°C above control near-natural conditions) in spring and summer on these seagrass species and quantified changes in growth performance, respiration, and gross primary production. Responses to these MHW varied by species and season. *C. nodosa* responded differently to MHWs in spring and summer, its net foliar growth rate being stimulated and maintained, respectively. *N. noltei* responded positively to moderate MHWs (+2, +4°C) in spring but showed vulnerability to the high-intensity MHWs (+6°C) in summer. *R. cirrhosa* remained resilient in spring and benefited from MHWs in summer, particularly at +4°C intensity, to stimulate growth. Hence, all three species were able to tolerate a single 20-day MHW with the species being increasingly better adapted in the following order *N. noltei, R. cirrhosa*, and *C. nodosa*. Increase of the initiation and growth of new leaves was a common phenological response observed in all three species, albeit at a lesser extent for *R. cirrhosa*. Our near-natural experimental approach reveals species- and season-dependent responses to MHWs, highlighting the complexity of seagrass resilience in lagoon ecosystems. These findings provide valuable insights to inform conservation strategies and improve the management of coastal habitats under future climate scenarios.

## Introduction

The semi-enclosed nature of the Mediterranean Sea causes it to warm two to three times faster than the global ocean (Diffenbaugh et al., 2007), making it one of the most affected bodies of saltwater in the world (Jordà et al., 2012; Templado, 2014), and a well-known hot-spot region for climate change (Giorgi, 2006). A rapid increase in the mean sea surface temperature (SST) of the Mediterranean sea is expected in the near future (Giorgi & Lionello, 2008). By the end of the 21ᵉ century, surface temperatures in the Mediterranean Sea are predicted to rise by 0.8°C to 3.8°C (IPCC, 2023). Besides these gradual changes, climate change is expected to increase the intensity, the frequency and the duration of sudden and relatively short-term events known as Extreme Climatic Events (ECEs) (Hobday et al., 2018; Jentsch & Beierkuhnlein, 2008). The sudden nature of ECEs leaves ecosystems with little time to adapt, making it crucial to consider them when understanding how ecosystems respond to climate change (Jentsch et al., 2007). Among ECEs, this study focusses on MHWs, characterized by sea temperatures exceeding seasonal thresholds for at least five consecutive days (Hobday et al., 2016), which can occur in all seasons. While not representing a new phenomenon, MHWs have increased in frequency, intensity, and duration over the past century, primarily due to anthropogenic climate change (IPCC, 2021). The Mediterranean Sea is expected to be highly sensitive to MHWs (Cramer et al., 2018; Darmaraki et al., 2019; Diffenbaugh et al., 2007; Lejeusne et al., 2010). Notably, the growing frequency and intensity of MHWs in Mediterranean sea have already been linked to mass mortality events (Darmaraki et al., 2019; Garrabou et al., 2009). Mediterranean coastal zone ecosystems are likely to be particularly sensitive to MHWs, as these events may potentially interact negatively with anthropogenic pressures such as chemical contamination, eutrophication and shoreline artificialization (de Wit, 2011; Lloret et al., 2008).

The Mediterranean coast is also characterized by numerous lagoons, transitional systems that link terrestrial and marine environments. These highly productive systems play a crucial role in regulating biogeochemical cycles and supporting biodiversity. This is related to their complex, mosaic-like landscapes, which provide refuge for numerous species by offering a diversity of habitats (de Wit, 2011; Iotti et al., 2023; Pérez-Ruzafa, Campillo, et al., 2019). Mediterranean coastal lagoons are subject to frequent environmental disturbances and fluctuations due to their shallowness and low thermal inertia (Nixon et al., 2004; Trombetta et al., 2019). Seasonal temperature variations are also more pronounced than in the adjacent sea (de Wit, 2011). These dynamics define them as naturally stressed environments, compelling their resident populations to develop greater tolerance and adaptability compared to their counterparts in open sea environments (Pérez-Ruzafa, Pérez-Ruzafa, et al., 2019).

The coastal lagoons comprise various linked habitats (Ouisse et al., 2023) that help species and communities thrive (Iotti et al., 2023). Within these lagoon environments, seagrass meadows play a key ecological role through their important biogeochemical and ecological functions. They contribute to climate regulation by storing carbon in sediments (Fourqurean et al., 2012) and support biodiversity (Colvin and Sne*LGR*ove, 2025), serving as nursery areas for many organisms and commercial fish species (Tournois et al., 2017). Four species of marine phanerogams are found in Mediterranean coastal lagoons. *Zostera marina* Linnaeus, 1753 is widely distributed in the northern hemisphere, i.e., in the temperate North Atlantic and North Pacific bioregions up to boreal altitudes. Its Southern limit is in the Mediterranean Sea (Green & Short, 2003; Short et al., 2007). *Nanozostera noltei* (Hornemann) (Tomlinson & Posluzny, 2001) (synonym *Zostera noltei*) ranges from the coasts of Norway in the temperate North Atlantic to Mauritania and occurs widely in the Mediterranean Sea (Green & Short, 2003). *Cymodocea nodosa* (Ucria) Ascherson, 1870 is mainly distributed along the Mediterranean coast but also occurs along the North Atlantic coast of Africa reaching Senegal, and is, therefore, considered as a southern subtropical-Mediterranean species (Green & Short, 2003; Mascaró et al., 2009). *Ruppia cirrhosa* (Petagna) Grande, 1918 mainly found in coastal waters and salt ponds in the Mediterranean, but also in estuaries and brackish lagoons in temperate zones of the northern hemisphere (Mannino et al., 2015). Each species has a different geographic distribution, and it can thus be expected that they will show different responses both towards steadily increasing temperatures and MHW. For instance, seawater temperature rise has been reported to enhance mechanisms such as respiration, growth and flowering in seagrasses (Duarte, 2002).

Global warming and MHWs can have widespread impacts on seagrass communities and the ecosystem functions and services they provide (Orth et al., 2006; Strydom et al., 2020). Globally, significant seagrass meadow mortalities and subsequent biodiversity losses have been associated with MHWs occurrences (Lefcheck et al., 2017; Shields et al., 2019; Smale et al., 2019; Strydom et al., 2020). The loss of seagrass habitat will have consequences on the associated functions and services they provide to coastal zones (Duarte, 2002).

*Zostera marina* is vulnerable with respect to increasing temperatures, both for the populations in its southern range of distribution but also in cooler climates. Shields et al., (2019) showed high temperature driven die-offs of *Z. marina* in Chesapeake Bay (USA) and its replacement by more tolerant *Ruppia* species. Numerous studies have highlighted the negative impact of MHW on *Z. marina*, including reduced growth (Kim et al., 2020) and a decreased photosynthetic performance (Winters et al., 2011). *Cymodocea nodosa,* which is considered as a thermo-tolerant species resilient to high temperatures is expected by some authors to thrive and potentially benefit from global warming in the coming decades (Savva et al., 2018). Modelling studies show that in the Mediterranean Sea under a scenario of rising seawater temperatures, *C. nodosa* could replace more than half of the *Posidonia oceanica* meadows (Papaki et al., 2020). Nevertheless, not all studies converge on this point of view. Hence, it was reported that MHW could have a negative impact on *C. nodosa,* several days after perturbation, i.e., the study of Deguette et al., (2022) shows a decline in the area/DW ratio due to a reduction of the photosynthetic capacity that could reduce growth. On the other hand, it was reported that *C. nodosa* is able to increase photosynthesis and adjust its photo-physiological processes to cope with a MHW (Costa et al., 2021; Marin-Guirao et al., 2016). Cardoso et al., (2008) reported that a MHW during the summer of 2003, had a negative impact by reducing growth production and biomass of *N. noltei* and also on the resilience of a restored population of this species, which ceased to recover after the disturbance. *N. noltei* is negatively impacted by warming conditions, leading to important reduction in shoot density (Repolho et al., 2017). In contrast, another study shows that shoot density was not affected by the simulated MHW (Simão, 2021). In temperate regions, *R. cirrhosa* exhibits resilience to warming (Rasmusson et al., 2021), and the authors suggest that it could outcompete heat-sensitive species. Temperature rise significantly affects *Ruppia* species’ germination, growth, and photosynthesis. Studies indicate that *R. cirrhosa* thrives between 20-30°C and maintains viability even at temperatures as low as 10°C, demonstrating greater temperature flexibility compared to other abovementioned seagrass species (Tsioli et al., 2019). The species reaches its thermal limit at temperatures between 32-34°C, beyond which it cannot survive (Tsioli et al., 2019). However, there are no studies on the effects of MHW on *R. cirrhosa* growth.

Experimental laboratory studies have proven essential for understanding the effects of isolated parameters. However, outdoor near-natural mesocosm studies that adopt a more realistic approach by mimicking seasonality, diel cycles, and stochastically fluctuating temperature regimes in Mediterranean lagoons remain absent. The scale of the study is also important to consider, as different responses can be detected depending on the population studied (Marin-Guirao et al., 2018; Nguyen et al., 2021). It is therefore essential to experimentally assess the impact of MHWs in Mediterranean coastal lagoons on seagrass populations, which are naturally adapted to these variable environments. These shallow ecosystems experience significant temperature variations, ranging from 4°C to over 30°C annually, with pronounced diel fluctuations that can reach 16–29°C in spring and 19–40°C in summer. Such strong fluctuations in shallow coastal lagoons are related to heat exchanges driven by the sun and the wind that vary strongly during a diel cycle. So far, however, most experimental studies (Deguette et al., 2022; Marin-Guirao et al., 2018; Rasmusson et al., 2021; Repolho et al., 2017; Tsioli et al., 2019), often indoor experiments, have used fixed temperature settings, which do not reflect the natural conditions. In these cases, the temperatures of both the controls and the warming treatments remained constant during the day. Consequently, direct comparisons with populations thriving in their natural environments are tricky. Similarly, comparisons with findings from the scarce literature remain challenging, as the reported responses often varied depending on the study location or focused on different variables than those examined in this study.

This study aims at examining for the first time how three seagrass species (*C. nodosa, N. noltei*, and *R. cirrhosa*) from a Mediterranean coastal lagoon respond to marine heatwaves of varying intensities during spring and summer. Because the negative effects of MHWs on *Z. marina* have been well-documented in the literature as abovementioned, this species is not included in the present study. We specifically focus on key traits including growth, productivity, and respiration. Based on two outdoor *ex-situ* near-natural mesocosm experiments that simulated typical Mediterranean lagoon temperature fluctuations, we evaluated the species’ responses to 20-day marine heatwaves at three intensity levels (+2°C, +4°C, and +6°C above control conditions), intensity and duration was determined based on preliminary studies conducted by our team (data from Petton et al., 2024) and aligned with IPCC projections. Our outdoor setup closely mimics a natural system, reproducing the temperature fluctuations of the shallow coastal lagoon while preserving natural incident light conditions. Hence, our study allowed us to replicate these fluctuations for baseline conditions and study the impact of +2 - +6 °C increases above these fluctuating baselines for the MHWs of different intensities. Applying fixed temperature upon a naturally fluctuating background as a single factor, while maintaining the other conditions as closely similar to *in situ* conditions, provides a clearer understanding of the impact of MHWs. We predict that heatwave intensity will affect growth rates and metabolism differently across species, depending on their individual thermal tolerances and the seasonal timing of heat exposure. Furthermore, we expect that the responses observed in this study will differ from those reported in other ecosystems, as seagrasses in Mediterranean coastal lagoons experience a particularly high level of environmental variability, which may influence their resilience to thermal stress.

## Materials and methods

### Study area and biological material collection

The Thau lagoon is a restricted coastal lagoon with two inlets on average 5 metres deep, (Fiandrino et al., 2017). The averaged water temperature varies during the year between 8 ± 1.8°C (winter) and 26 ± 1.8°C (summer), while salinity varies between 35 ± 2.0 in early spring and 41 ± 1.5 in summer (REPHY network, 2021: January 2016 - May 2021).

The taxonomy of the Mediterranean lagoon seagrass species is in agreement with the World Register of Marine Species (WoRMS - https://www.marinespecies.org), which integrated the recent taxonomic revision in the Zosteraceae family (Tomlinson & Posluzny, 2001; Sullivan & Short, 2023). *Nanozostera noltei* was sampled in the southern part of the Thau lagoon (3.57330 / 43.35990) at approximately 2.5-meter depth. *Cymodocea nodosa* was sampled in the northern part of the lagoon (3.65503 / 43.40541) at 1.3-meter depth. *Ruppia cirrhosa* was sampled in the eastern part of the lagoon (3.63793 / 43.39355) at 1.3 meters depth. Since *Zostera marina* reaches its southern distribution limit in Mediterranean lagoon (Green & Short, 2003; Rueda et al., 2008) and numerous study have demonstrate its vulnerability to MHWs (see Introduction), it was not included in the present study.

For each species, a total of 144 shoots with their roots and rhizomes were meticulously collected by scuba diving on 18 April 2023 and on 18 July 2023 for the spring and summer experiments, respectively. The collection process involved hand-picking the shoots underwater from different locations within the seagrass meadow to ensure a representative sampling of the area. The shoots were then transported to the laboratory using buckets filled with the lagoon water. The collected shoots were replanted in experimental units (see below) in reconstituted sediments of the same granulometry. Therefore, sediments were collected from each sampling site and upon return to the laboratory dried (60°C for 24 hours) and subsequently sieved (2 mm mesh) to remove fauna and plant residues.

### Experimentation

We conducted an outdoor near-natural mesocosm using four fiberglass tanks (200L) filled with Thau lagoon’s water. The experimental setup consisted of four tanks reflecting four treatments:

- A control tank regulated to follow the natural temperature variations of the Thau lagoon.
- A tank with temperature elevated by +2°C above the control.
- A tank with temperature elevated by +4°C above the control.
- A tank with temperature elevated by +6°C above the control

Before heatwave exposure, all specimens were maintained under the same regulated experimental conditions, reflecting the natural conditions of the Thau lagoon during 15 days. Heatwaves of varying intensities were applied over a period of 20 days. After heatwave exposure period, all specimens were maintained under the same conditions as the Control treatment for an additional 15 days to simulate the natural conditions of the lagoon (Figure 1). We conducted this experimental design across two seasons: spring and summer 2023.

**Figure 1.**
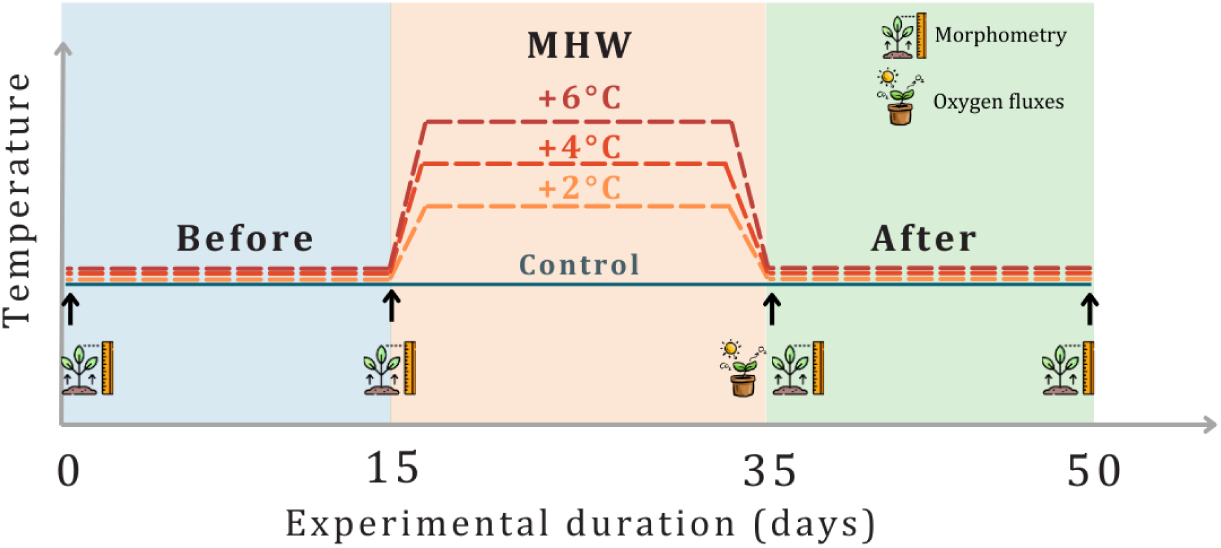
: Timelines for the experimental conditions and timing of sampling in the mesocosms showing the main events with respect to natural conditions, i.e., acclimation period before heatwave exposure (blue), during heatwave exposure (orange), and return period after heatwave exposure (green). Arrows with their associated pictograms indicate the timing of samplings for morphometry measurements on seagrass shoots and oxygen fluxes measurements.

To regulate the temperature, we used a heater and a cooler in each tank. A sensor was deployed in the Thau lagoon to measure the real-time water temperature (readings taken every 10 seconds). This data was automatically transmitted to a program, enabling real-time adjustments of the water temperature in the control tank to the same temperature as measured in the lagoon. In parallel, in the tanks used for the other treatments, the temperatures were adjusted following the timelines shown in Figure 1. Three pumps per tank recirculated the water to ensure a homogeneous water bath.

In each experimental unit, six shoots, along with their rhizomes and roots, were selected. The rhizome segments were standardized according to species-specific protocols: for *N. noltei*, rhizomes were adjusted to three segments; for *R. cirrhosa*, rhizomes were adjusted to two segments in spring experiments and three segments in summer experiments; and for *C. nodosa*, only horizontal rhizomes were retained. The standardized shoots were then placed in individual 3L cylindrical glass experimental units (10 cm in diameter, 30 cm in height) that had been pre-filled with 550 cm³ of sediment (approximately 7 cm in depth). This resulted in a total of 36 shoots per treatment (control, +2°C, +4°C, and +6°C) for each species. Subsequently, each set of six experimental units per species was arranged in experimental tanks, leading to a total of 18 experimental units per tank.

Water temperature was monitored continuously (a reading every 5 minutes) with autonomous sensors (HOBO, UA-002-64, ± 0,2°C) placed in each experimental tank. The experimental system was set up outdoors to take advantage of the natural variation in light, with the aim of reproducing *in-situ* conditions as closely as possible. The available light intensity was continuously measured in close proximity to the experimental system using an LI-190 planar sensor probe connected to a data acquisition unit (Campbell, CR1000).

The water level and salinity in the experimental tanks were monitored three times a day, and adjustments were made by refilling with fresh water to offset evaporation. Additionally, the water in each treatment was refreshed weekly to maintain adequate nutrient levels. The samples for nutrients concentrations of nutrients (NH_4_^+^, NO_3_^-^+NO_2_^-^ and PO_4_^3-^) was estimated from spot samples taken twice a week.

### Analysis

Morphological, physiological and metabolic responses of three seagrass species were measured throughout the experiment (Figure 1). Nutrients sample was analysed in the laboratory using the method described by David et al. (2019).

At the onset of the experiment, we conducted initial morphological measurements for each species. Leaf area was measured (length × width) and the number of leaves per shoot was counted for all species. Leaf measurements were performed on individual shoots, involving the temporary removal of each specimen from the sediment. Subsequently, measurements were taken for each specimen, including the number of shoots, leaves, length and width of leaves, length and width of rhizomes, ensuring a comprehensive assessment of morphological characteristics. After the measurements, each specimen was carefully placed back into the experimental units during each measurement session.

These measurements, which reflect the creation of new plant matter, were first used to calculate individual leaf growth rates. For each leaf, the growth rate was determined using the following formula:

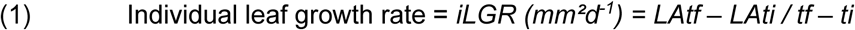

where LA is the leaf area (length * width) at final time (tf) and initial time (ti). For newly formed leaves, corrections were applied. Negative values, which represented cut leaves, were adjusted to zero. To obtain the growth rate (mm^-2^.d^-1^) at the shoot level, these individual leaf growth rates were then summed for all leaves within each shoot:

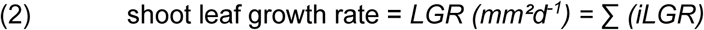

New leaf gain rate (mm-2.d-1) was quantified by tracking individual leaf appearance between consecutive measurements (ti and tf). For each shoot, new leaf gain rate was calculated as:

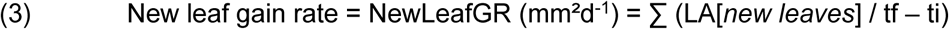

where LA is the recorded area of new leaf. Shoots without leaves creation were assigned a zero rate.

Old leaf gain rate (mm^-2^.d^-1^) was quantified by tracking the retention of two leaves between consecutive measurement times (ti and tf). For each shoot, old leaf gain rate was calculated as:

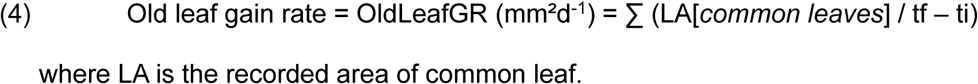

The Leaf Growth Rate (*LGR*) was defined as the sum of *NewLGR* and *OldLeafGR*, hence

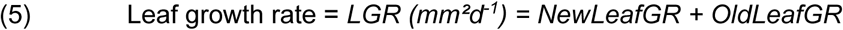

In addition, we calculated a 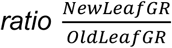 *(hereafter referred to as: New-Old leaf* ratio) to characterise the proportions of growth allocated to the formation of new and old leaves, respectively.

Because the leaf growth rate does not take into account the loss of leaf area due to the falling leaves, the leaf loss rate (mm^-2^.d^-1^) were quantified by tracking individual leaf disappearance between consecutive measurements (ti and tf). For each shoot, leaf loss rate was calculated as:

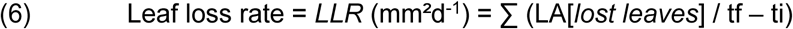

where LA is the last recorded leaf area before loss. Shoots without leaf loss were assigned a zero rate.

In each experimental unit, at the end of heatwave exposure, oxygen fluxes were measured using optical sensor (WTW FDO 925) through short-term incubations of 1 to 1.5 hours per session (Figure 1). This process involved calculating the difference between initial and final oxygen concentrations in both dark and light conditions to determine community respiration (CR: negative or zero) and Net Community Production (NCP: negative, positive or zero), respectively. Gross Community Productivity (GCP, positive or zero) was calculated by adjusting NCP for CR and normalizing it to the total leaf area (*tot LA*) of seagrass shoots in each experimental unit at the time of measurement.

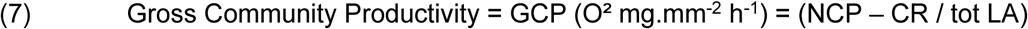

### Statistical analysis

All statistical analyses were conducted using Rstudio (R version 4.4.1) with significance set at p=0.05. To assess the impact of different experimental conditions on temperature, daily averages of the measurements were first calculated. Due to non-normality, daily temperature averages were assessed across experimental periods using the Kruskal-Wallis test, with subsequent pairwise comparisons conducted using the Dunn test and Benjamini-Hochberg correction. Similar Kruskal-Wallis tests were applied to salinity, light, and nutrient concentration data. Leaf Growth rate (*LGR*) and Leaf Loss rate (*LLR*) were analysed using linear mixed-effect models, with treatments and experimental periods as fixed effects and experimental units (replicate) in random effects. Post-hoc pairwise comparisons were using emmeans package, with p-value adjusted for multiple comparisons using the Tuckey method. Graphic diagnostics were used to verify normality and homogeneity of variances. Oxygen fluxes were analysed using Kruskal-Wallis test. Occasionally, when during the acclimation period, treatments presented a *LGR* significantly different from the control, we did not further report the pairwise comparisons between these treatments and the control during heatwave exposure (noted as “*nc*” in Table 2-a), as their response might have depended on prior conditions rather than solely on the experimental treatment. Nevertheless, within these treatments it was still meaningful to compare *LGR* during the heatwave and return periods with the acclimation period (Table 2-b).

## Results

### Experimental conditions

Light, salinity and inorganic nutrient concentrations in the outdoor experimental set-up during experiments are show in Table 1 for the spring and summer experiments, respectively. The length of the photoperiod increased by 11.3 % (1.52 hours) from the start to the end of the spring experiment, accompanied by a rise in the average daily light integral (DLI) by 84.7 % (Table 1; DLI = 23.5 ± 16.4 to 43.4 ± 6.1 mol photons.m^-^².d^-1^). In contrast, during the summer experiment, the length of the photoperiod decreased by 13.1 % (1.93 hours), leading to a corresponding decline in average DLI of 29.9 % (39.8 ± 12.25 to 27.9 ± 11.16 mol photons.m^-^².d^-1^). Salinity was homogeneous among treatments and experimental periods with an overall mean of 38.9 ± 0.9 psu in spring and 40.2 ± 1.6 psu in summer. Nutrient concentrations remained stable during both experiments (Table 1).

**Table 1:** Environmental conditions in the incubation system, including photoperiod, daily light integrate (DLI), salinity, and concentrations of nutrients (NH4, NOx, and PO4) before, during, and after heatwave exposure, for both spring and summer seasons. Photoperiod and DLI data were calculated in proximity to the experimental system, while the other parameters were measured on the experimental tanks water. The values represent the means ± standard deviation of all conditions (control, +2, +4, and +6) since no statistical differences were observed.

|  |  | Before | Heatwave | After |
| --- | --- | --- | --- | --- |
|  |  | 15 days | 20 days | 15 days |
| Spring | Photoperiod (h) | 13.40 - 13.95 | 14.02 - 14.57 | 14.53 - 14.92 |
| | DLI (mol photons.m<sup>-2</sup>.d<sup>-1</sup>) | 23.5 ± 16.4 | 32.7 ± 11.8 | 43.4 ± 6.1 |
| | Salinity | 38.5 ± 0.7 | 39.3 ± 0.9 | 38.8 ± 0.7 |
| | NH<sub>4</sub> (μmol.L<sup>-1</sup>) | 0.63 ± 0.95 | 0.50 ± 0.85 | 0.63 ± 1.03 |
| | NO<sub>x</sub> (μmol.L<sup>-1</sup>) | 0.40 ± 0.34 | 0.38 ± 0.29 | 0.52 ± 0.57 |
| | PO<sub>4</sub> (μmol.L<sup>-1</sup>) | 0.075 ± 0.05 | 0.109 ± 0.08 | 0.163 ± 0.12 |
| Summer | Photoperiod (h) | 14.77- 14.30 | 14.48 - 13.47 | 13.43 - 12.83 |
| | DLI (mol photons.m<sup>-2</sup>.d<sup>-1</sup>) | 39.8 ± 12.25 | 32.6 ± 9.73 | 27.9 ± 11.16 |
| | Salinity (psu) | 40.67 ± 1.76 | 39.70 ± 1.45 | 40.53 ± 1.63 |
| | NH<sub>4</sub> (μmol.L<sup>-1</sup>) | 0.43 ± 0.64 | 0.59 ± 1.09 | 0.55 ± 0.84 |
| | NO<sub>x</sub> (μmol.L<sup>-1</sup>) | 0.49 ± 0.36 | 0.57 ± 0.66 | 0.45 ± 0.44 |
| | PO<sub>4</sub> (μmol.L<sup>-1</sup>) | 0.06 ± 0.05 | 0.07 ± 0.05 | 0.07 ± 0.06 |

Before heatwave exposure (acclimation periods), water temperatures for both spring and summer experiments were uniform across treatments (Figure 2). Following seasonal patterns, water temperatures were consequently higher in summer than in spring, i.e., 27.80 ± 3.80 °C, and 20.63 ± 3.03 °C, respectively. During the heatwave exposure periods, both in spring and summer, the temperature in the control treatments (+ 0 °C) did not vary significantly with respect to the acclimation period. In contrast, there were significant temperature differences between the control and the three heatwave treatments. Post hoc comparisons indicated that water temperature increased significantly for the +2°C, +4°C, and +6°C treatments compared to the control, with mean differences of 1.69 ± 0.55°C (p = 0.02), 4.07 ± 1.16°C (p < 0.001), 5.78 ± 1.02°C (p < 0.001), respectively in spring and 1.51 ± 1.40°C (p = 0.06), 3.02 ± 1.65°C (p < 0.001), 4.39 ± 1.66°C (p < 0.001) in summer. During the final experimental period (the return phase), each treatment returned to follow the natural temperature fluctuations in Thau lagoon (Figure 2). Unexpectedly, the return phase in spring coincided with an uncommon natural heatwave that was comparable to a heatwave treatment of approximately +4°C (25.12 ± 2.46 °C). In contrast for the summer experiment, the temperatures were lower than during the acclimation and heatwave exposure periods (24.93 ± 3.67 °C), as expected for the season.

**Figure 2.**
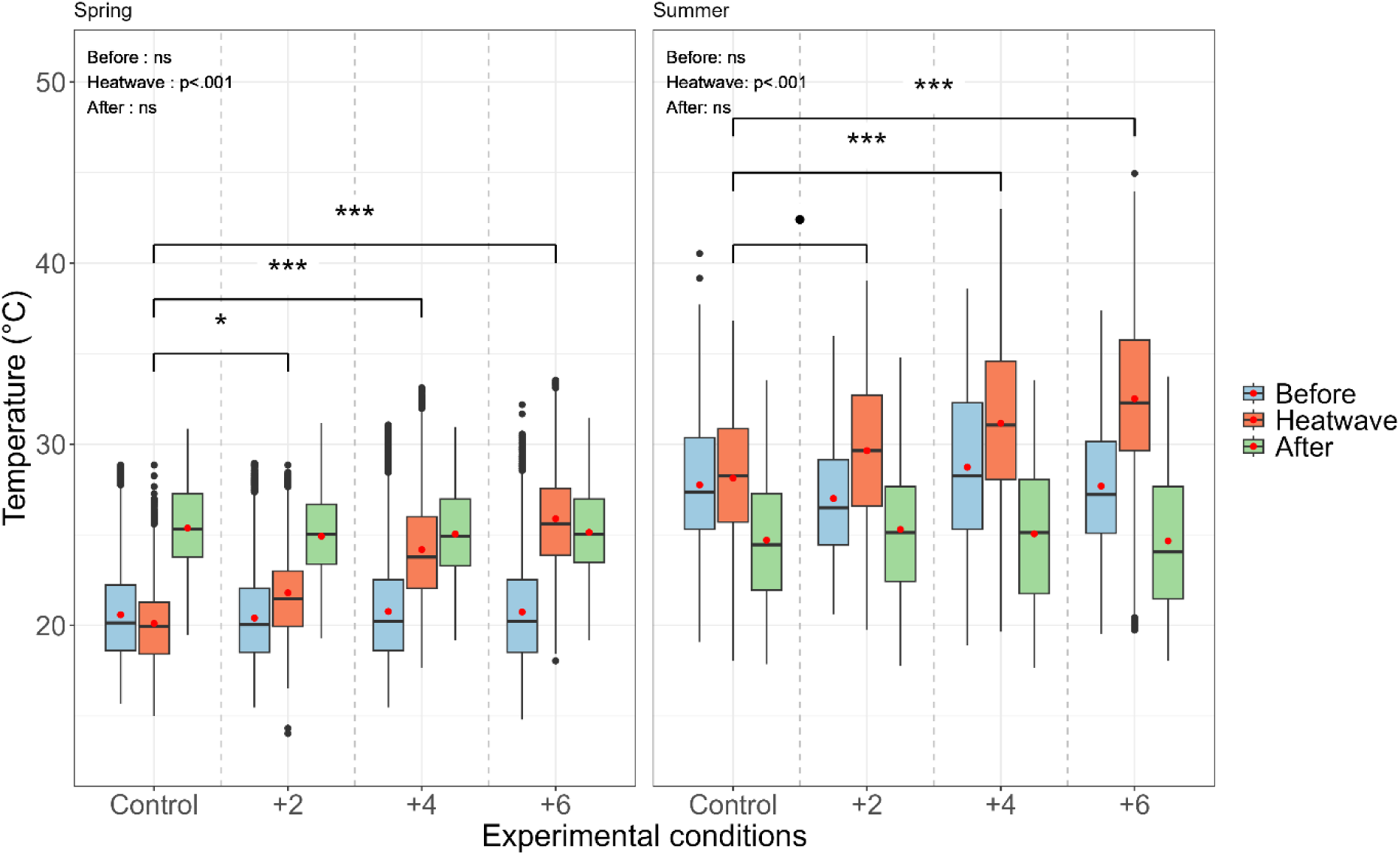
: Temperature variations in spring (left) and in summer (right) before heatwave exposure (in blue), during heatwave exposure (in orange), and after heatwave exposure (in green) for the ‘control’, ‘+2’, ‘+4’, and ‘+6’ treatments. Kruskal-Wallis tests indicate the significant differences between treatments. Data are represented as box plots, with the mean shown in red. The horizontal line within each box represents the median, and the edges of the boxes correspond to the 25^th^ and 75^th^ percentiles. The ends of the whiskers indicate the 5^th^ and 95^th^ percentiles, and the dots represent outliers.

### Seagrass responses to experimental conditions

While all the statements based on pairwise comparisons of experimental results for Leaf Growth rate (*LGR*) and Leaf Loss rate (*LLR*) reported in this section are supported by statistical tests (p < 0.05), we did neither report their results in the text nor in the corresponding figures (Figures. 2-4, panels a-d) to maintain their readability, but rather refer to Table 2 (Post-hoc pairwise comparisons with the p-values adjusted for multiple comparisons using the Tuckey method, see methods).

**Table 2:**
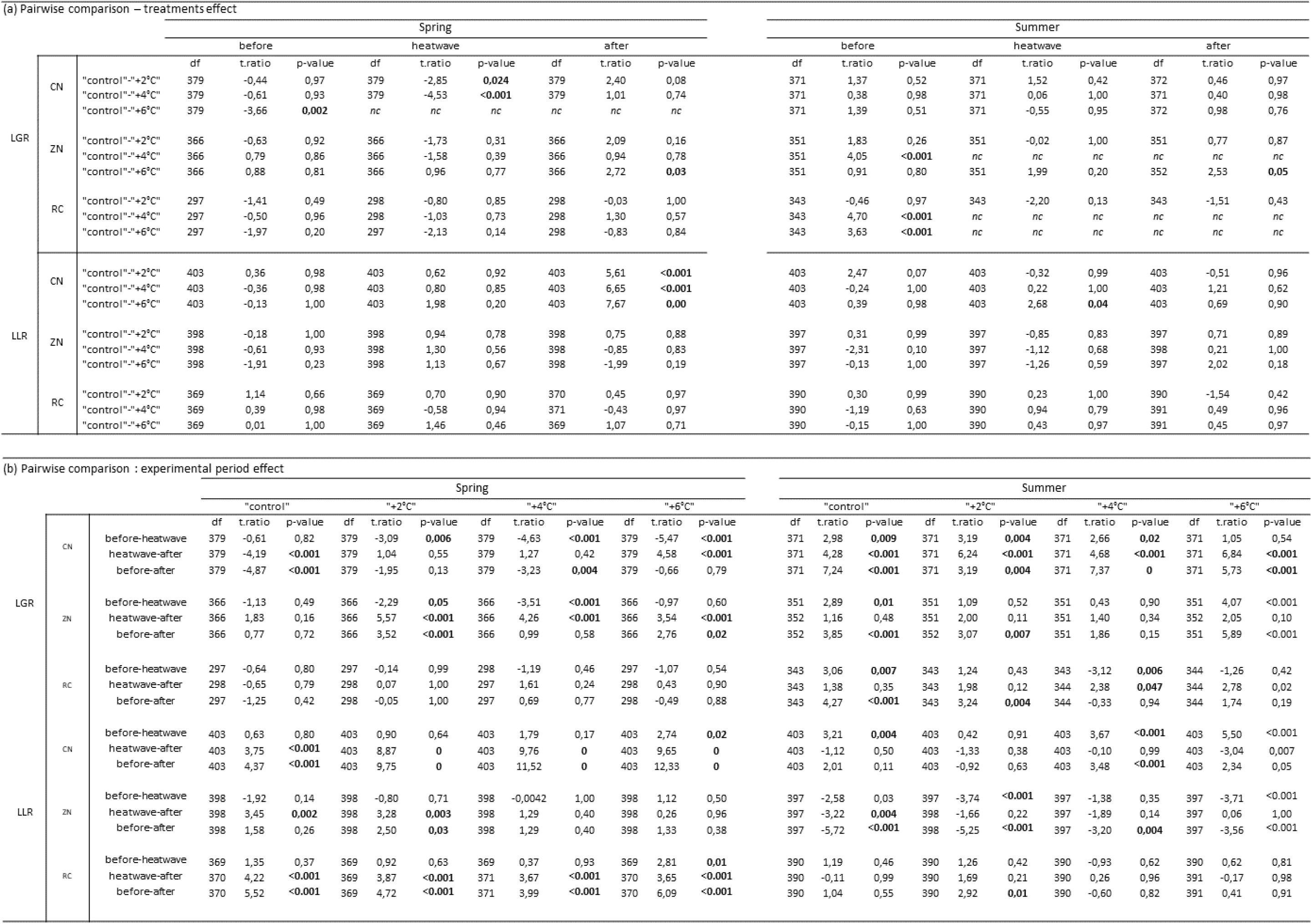
Pairwise comparisons of the effects of heatwave treatment and experimental period on the *LGR* and *LLR* of *Cymodocea nodosa* (CN), *Nanozostera noltei* (ZN), and *Ruppia cirrhosa* (RC). “nc” indicates that statistics were not calculated when there was a significant difference (p > 0.05) during the acclimation phase (“before”).*Ruppia cirrhosa* (RC).

#### Acclimation periods before heatwave exposure

Prior to heatwave exposure (acclimation period), leaf growth rate (*LGR*) with 4 exceptions (see Table 2) and all leaf loss rates (*LLR*) were homogeneous across treatments for all species in both spring and summer. The *New-Old leaf* ratio was close to one for the three species in both spring and summer, which showed an equilibrium between growth of new and old leaves. On average, the rate of gain of new leaves was higher in summer for *C. nodosa* and *R. cirrhosa* but lower for *N. noltei*. Meanwhile, the old leaf gain rate was higher in summer for *C. nodosa* and *N. noltei* and remained similar for *R. cirrhosa* (Figures 3b–e, 4b–e, and 5b–e).

**Figure 3:**
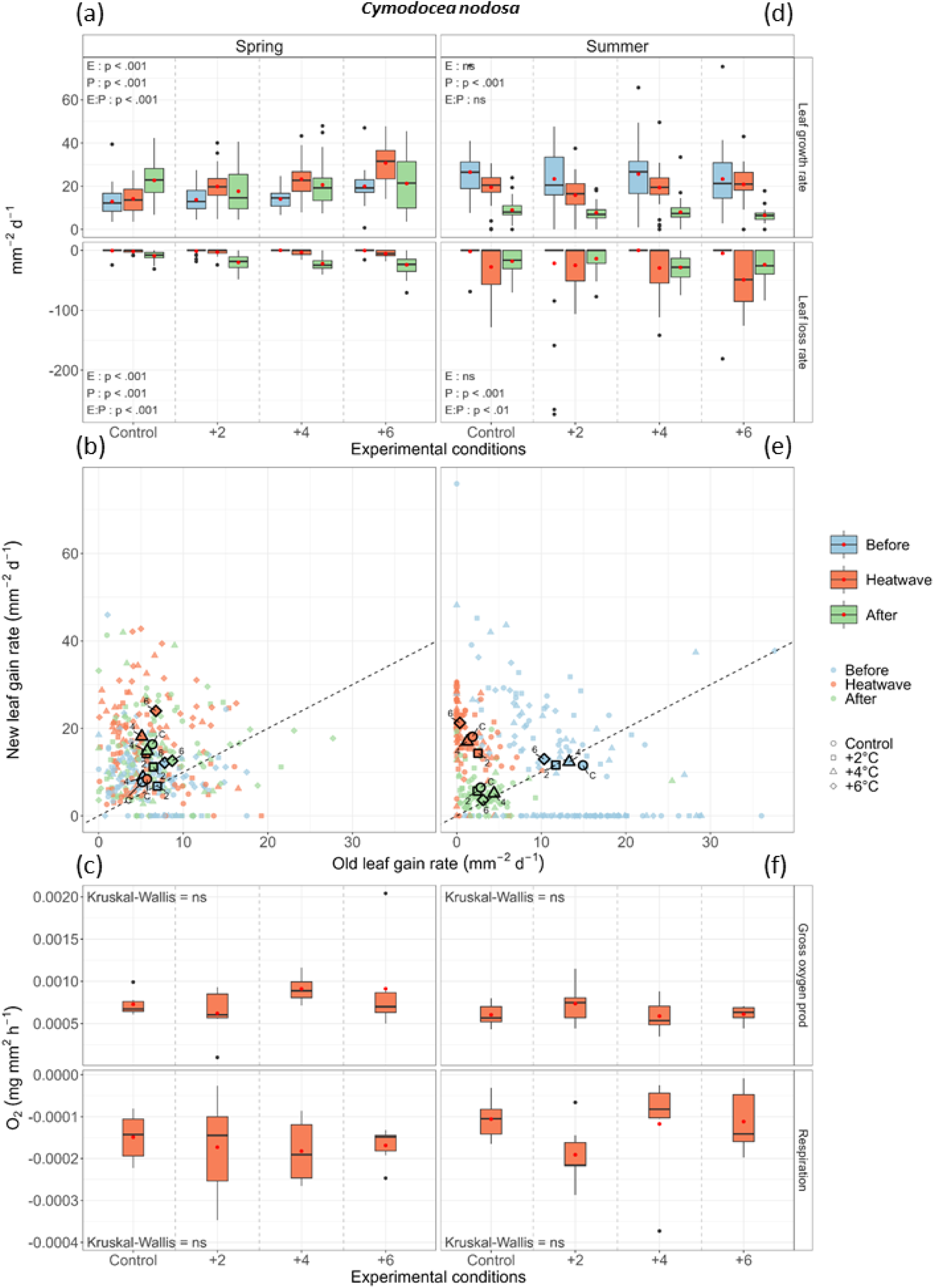
Effects of experimental marine heatwaves on the growth and physiological responses of *Cymodocea nodosa*. (a, d) Leaf Growth Rate (*LGR*) and Leaf Loss Rates (*LLR* - mm² d⁻¹) measured in spring and summer across four experimental treatments (Control, +2°C, +4°C, +6°C). Statistical significance is indicated for experimental effects (E), period effects (P), and their interaction (E:P) with their p-values derived from the ANOVA of a linear mixed-effects model (lmerTest), post-hoc pairwise comparisons are listed in Table 2. (b, e) *New-Old leaf* ratio across experimental periods (Before, Heatwave, After). Data points are colored by period, and different symbols indicate temperature treatments. The dashed line represents a 1:1 relationship. Larger points represent the means of the respective data clouds. (c, f) Gross oxygen production and respiration rates (mg O₂ mm⁻² h⁻¹) under different temperature treatments. Kruskal-Wallis tests indicate no significant differences between treatments. Boxplots display the distribution of the data, with the mean shown in red. The horizontal line within each box represents the median, while the box edges correspond to the 25th and 75th percentiles. Whiskers extend to the 5th and 95th percentiles, and individual points represent extreme values

**Figure 4:**
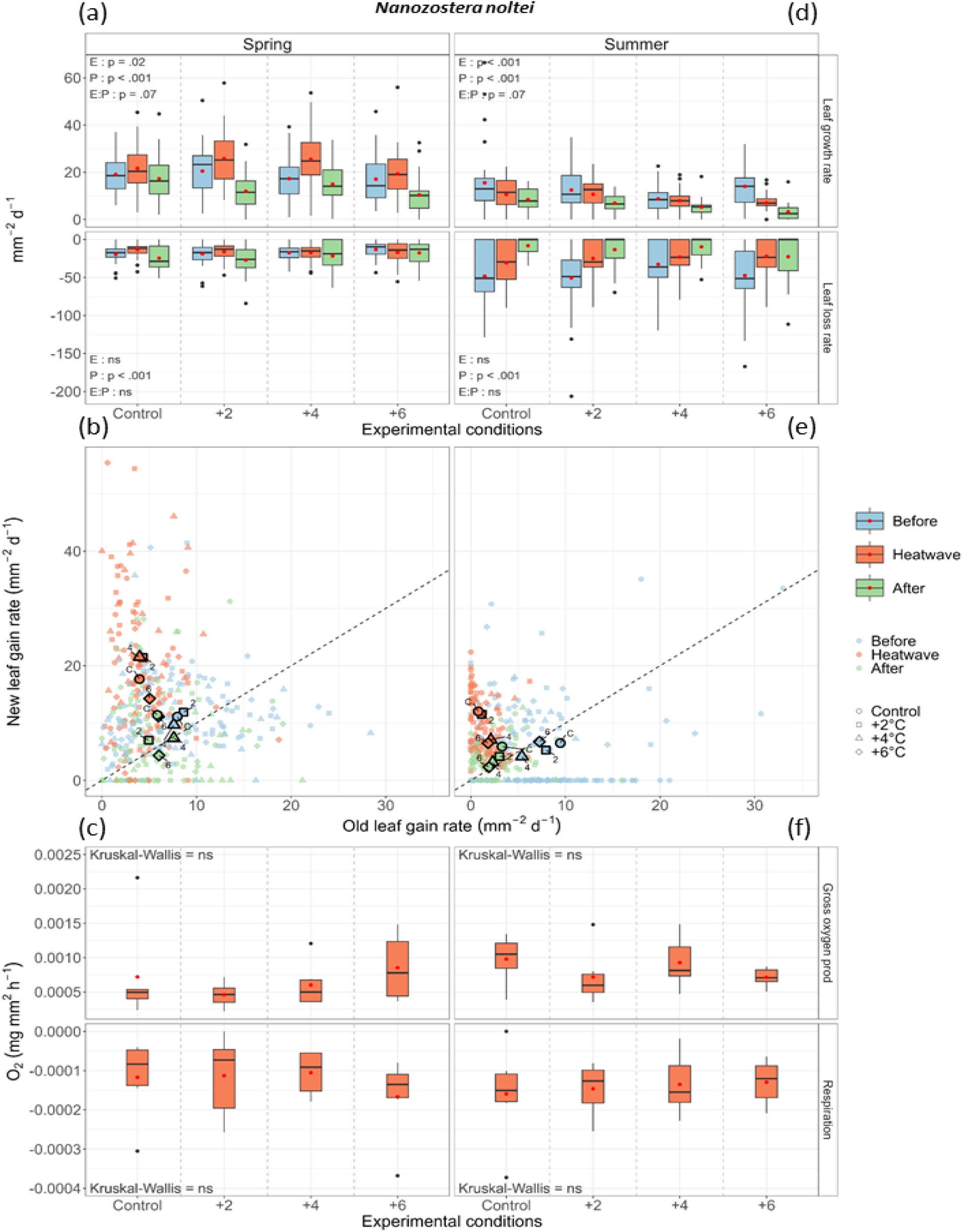
Effects of experimental marine heatwaves on the growth and physiological responses of *Nanozostera noltei*. (a, d) Leaf Growth Rate (*LGR*) and Leaf Loss Rates (*LLR* - mm² d⁻¹) measured in spring and summer across four experimental treatments (Control, +2°C, +4°C, +6°C). Statistical significance is indicated for experimental effects (E), period effects (P), and their interaction (E:P) with their p-values derived from the ANOVA of a linear mixed-effects model (lmerTest), post-hoc pairwise comparisons are listed in Table 2. (b, e*). New-Old leaf* ratio (growth ratio of new to old leaves) across experimental periods (Before, Heatwave, After). Data points are colored by period, and different symbols indicate temperature treatments. The dashed line represents a 1:1 relationship. Larger points represent the means of the respective data clouds. (c, f) Gross oxygen production and respiration rates (mg O₂ mm⁻² h⁻¹) under different temperature treatments. Kruskal-Wallis tests indicate no significant differences between treatments. Boxplots display the distribution of the data, with the mean shown in red. The horizontal line within each box represents the median, while the box edges correspond to the 25th and 75th percentiles. Whiskers extend to the 5th and 95th percentiles, and individual points represent extreme values.

**Figure 5:**
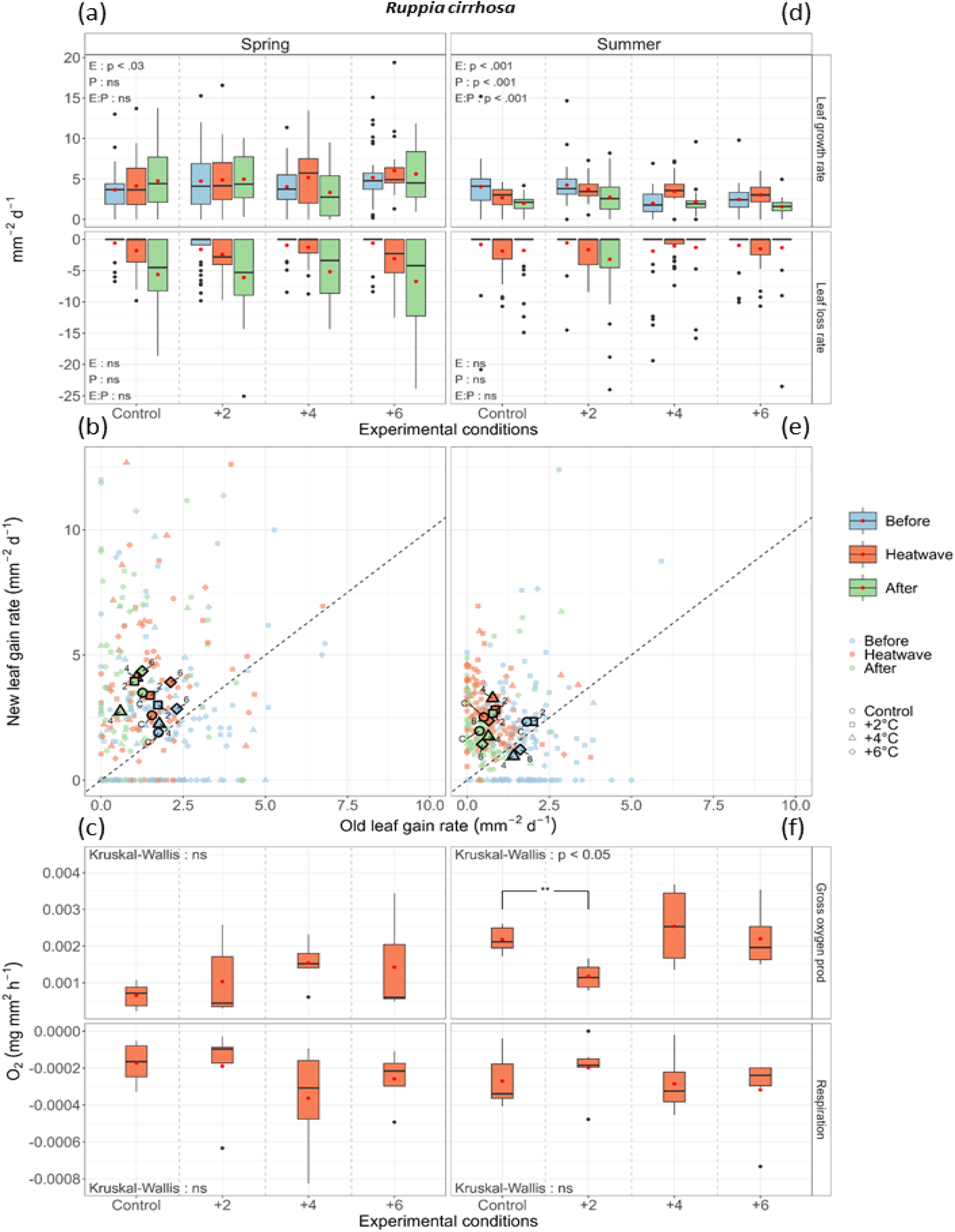
Effects of experimental marine heatwaves on the growth and physiological responses of *Ruppia cirrhosa*. (a, b) Leaf Growth Rate (*LGR*) and Leaf Loss Rates (*LLR* -mm² d⁻¹) measured in spring and summer across four experimental treatments (Control, +2°C, +4°C, +6°C). Statistical significance is indicated for experimental effects (E), period effects (P), and their interaction (E:P) with their p-values derived from the ANOVA of a linear mixed-effects model (lmerTest). Post-hoc pairwise comparisons are listed in Table 2. (b, e) *New-Old leaf* ratio (growth ratio of new to old leaves) across experimental periods (Before, Heatwave, After). Data points are colored by period, and different symbols indicate temperature treatments. The dashed line represents a 1:1 relationship. Larger points represent the means of the respective data clouds. (c, f) Gross oxygen production and respiration rates (mg O₂ mm⁻² h⁻¹) under different temperature treatments. Kruskal-Wallis tests indicate the significant differences between treatments. Boxplots display the distribution of the data, with the mean shown in red. The horizontal line within each box represents the median, while the box edges correspond to the 25th and 75th percentiles. Whiskers extend to the 5th and 95th percentiles, and individual points represent extreme values.

### Cymodocea nodosa

#### Cymodocea nodosa response during the spring experiment

No mortality was observed for *C. nodosa* throughout the entire spring experiment. During heatwave exposure, *LGR* in Controls remained constant compared to the acclimation period (14.11 ± 6.19 mm².d-1), whereas it significantly increased in the heatwave treatments (+2°C = 19.85 ± 7.78 mm².d^-1^; +4°C = 23.23 ± 8.31 mm².d^-1^; +6°C = 30.76 ± 8.80 mm².d^-1^). After heatwave exposure, *LGR* was homogeneous across treatments. Hence, upon the transition from heatwave exposure to the return period, *LGR* increased in controls (22.71 ± 8.90 mm².d^-1^), remained steady for the +2°C and +4°C treatments (17.65 ± 10.48 mm².d^-1^ and 20.58 ± 10.23 mm².d^-1^, respectively) and decreased for the +6°C treatment (21.26 ± 12.56 mm².d^-1^). The variability of these *LGR* was highest in the +6°C treatment (Figure 3a, Table 2). *LLR* of *C. nodosa*, in spring (Figure 3a), increased during the experiment, the *LLR* was homogeneous between treatments during heatwave exposure (Figure 3a). After heatwave exposure, *LLR* was significantly higher for those individuals who had experienced a heatwave (Figure 3a). The relationship between the *NewLeafGR* and the *OldLeafGR* remained stable for the control, but showed an increasing *New-Old leaf* ratio for treatments during heatwave exposure (Figure 3b). This increase was more pronounced in the +4°C and +6°C treatments, where the ratio exceeded 3. After heatwave exposure, the *New-Old leaf* ratio increased for controls (ratio = 2.6) but decreased for treatment (still higher than 1, Figure 3b). During heatwave exposure both gross oxygen production and respiration rates were similar across all treatments (Figure 3c).

#### Cymodocea nodosa response during the summer experiment

No mortality was observed before heatwave exposure, whatever the treatment. During heatwave exposure, one shoot died in the controls, three shoots at +2°C, none at +4°C and one at +6°C. After heatwave exposure, mortality remained low with one shoot in the controls, two at +2°C, five at +4°C and three at +6°C. During heatwave exposure, no significant differences were observed in *LGR* across treatments. The *LGR* of the Control treatment and the +2°C and the +4°C treatments significantly decreased compared to the period before heatwave exposure (19.52 ± 7.54 mm².d^-1^, 15.89 ± 8.72 mm².d^-1^ and 19.38 ± 9.43 mm².d^-1^, respectively), while the *LGR* remained similar for the +6°C treatment (20.84 ± 8.91 mm².d^-1^). After heatwave exposure, *LGR* was homogeneous across treatments. Compared to heatwave exposure, *LGR* drastically decreased for all treatments (Figure 3d, Table 2b). During heatwave exposure, *LLR* significantly increased for the Control, +4°C, and +6°C treatments, while the *LLR* remained steady for the +2°C treatment. *LLR* during heatwave exposure was highest for the +6°C treatment (Figure 3d). After heatwave exposure, *LLR* was similar across treatments and comparable to the period during heatwave exposure, except for the +6°C treatment where it significantly decreases (−24.02 ± 22.38 mm².d^-1^; Figure 3d). During heatwave exposure, the relationship between *NewLeafGR* and *OldLeafLGR* of all treatments deviated from equilibrium by increasing the *New-Old leaf* ratio. The mean *OldLeafLGR* decreased sharply and approaches zero for the +4°C and +6°C treatments, which directly impacts the ratio, making it particularly high for these two treatments (ratio = 13.7 ± 18.00; 56.56 ± 87.46, respectively, Figure 3e). The mean *NewLeafGR* remained similar to the Control and +2°C treatments. After heatwave exposure, in all treatments the *New-Old leaf* ratio returned to close to 1, however with lower *NewLeafGR* and *OldLeafLGR* than observed before heatwave exposure (Figure 3e). During heatwave exposure, both gross oxygen production and respiration rates were similar across all treatments (Figure 3f).

### Nanozostera noltei

#### Nanozostera noltei response during the spring experiment

Overall shoot mortality was very low for the 36 shoots used for each treatment. In the control treatments, no shoot was lost. At +2°C one shoot died during heatwave exposure and another after. At +4°C, one shoot died during this period, and two afterwards. At +6°C, three shoots died during exposure, and two more afterwards. During heatwave exposure, *LGR* of the Control treatment remained stable compared to the acclimation period, while the *LGR* of the other treatments was not statistically different from the Control (Figure 4a - Table 2a). However, pairwise comparisons within each treatment comparing the acclimation and subsequent heatwave exposure periods showed an increase in the +4°C treatment between the before and heatwave exposure periods (Figure 4a – Table 2b). After heatwave exposure, the *LGR* values remained similar between treatments, with only the +6°C treatment showing significantly lower rates. Specifically, while the Control treatment maintained stable *LGR* values, all experimental treatments showed decreased rates (Figure 4a). The *LLR* remained homogeneous with the Control across all treatments throughout the entire experiment (Figure 4a – Table 2a). However, pairwise comparisons within each experiment indicate a tendency for decrease in the Control and +2°C treatment between heatwave exposure and after (Figure 4a – Table 2b). During heatwave exposure, *NewLeafGR* increased across all treatments, with the most substantial rise observed at +2°C and +4°C, where it more than doubled compared to before heatwave exposure (Figure 4b). *OldLeafGR* generally decreased, though this reduction was less pronounced in the +6°C treatment (Figure 4b). The *New-Old leaf* ratio rise significantly in all treatments, especially at +4°C, indicating a shift toward new leaf growth over old leaf maintenance. After heatwave exposure, *NewLeafGR* returned to levels similar to or below exposure period, while *OldLeafGR* recovered slightly across all treatments. Consequently, the ratio dropped back close to pre-heatwave values, reflecting a return toward equilibrium. During heatwave exposure, both gross oxygen production and respiration rates were similar across all treatments (Figure 4c).

#### Nanozostera noltei response during the summer experiment

One shoot died before heatwave exposure in the controls, two shoots at +2°C, three at +4°C, and one shoot at +6°C. During heatwave exposure, six shoots died in the controls group, five at +2°C, four at +4°C, and four plants at +6°C. After heatwave exposure, two plants died in the controls, one at +2°C, one at +4°C, and six at +6°C. During heatwave exposure, no significant differences were observed in *LGR* across treatments (Figure 4d – Table 2a). The *LGR* of the Control treatment and the +6°C treatment significantly decreased compared to the period before heatwave exposure (10.55 ± 6.75 mm².d^-1^ and 7.05 ± 4.07 mm².d^-1^, respectively), while the *LGR* remained similar for the +2°C and the +4°C treatment (10.64 ± 6.38 mm².d^-1^ and 7.97 ± 4.69 mm².d^-1^ - Figure 4d – Table 2b). After heatwave exposure, the *LGR* was homogeneous across treatments, with a tendency to be lower in the +6°C treatment (Figure 4d). During heatwave exposure, *LLR* significantly decreased for the Control, +2°C, and +6°C treatments, while the *LLR* remained steady for the +4°C treatment. The *LLR* during heatwave exposure were significantly highest for the +6°C treatment (Figure 4d). After heatwave exposure, *LLR* was still decreasing for the Control treatment (−8.17 ± 11.28 mm².d^-1^), and remained homogeneous for the treatments (+2°C = −13.31 ± 19.47 mm².d^-1^; +4°C = −9.73 ± 15.27 mm².d^-1^; +6°C = −22.65 ± 28.39 mm².d^-1^). During heatwave exposure, *NewLeafGR* increased across all treatments, with the highest rise in the Control and +2°C treatments (Figure 4e). *OldLeafGR* decreased sharply in the control and +2°C treatments. The *New-Old leaf* ratio increased in the Control and +2°C treatments, indicating a shift toward new leaf growth, whereas it remained much lower for +4°C and +6°C (Figure 4e). After heatwave exposure, the *New-Old leaf* ratio returned close to 1 with lower *NewLeafGR* and *OldLeafLGR* values than before heatwave. During heatwave exposure, both gross oxygen production and respiration rates were similar across all treatments (Figure 4f).

### Ruppia cirrhosa

#### Ruppia cirrhosa response during the spring experiment

*Ruppia cirrhosa* showed substantially more mortality than the other two species. Ten shoots died in the controls in total (5 during heatwave, 5 after). At +2°C, 13 shoots died (1 before, 6 during heatwave, 6 after). At +4°C, 16 shoots died (4 before, 10 during heatwave, 2 after). At +6°C, 14 shoots died (11 during heatwave, 3 after). The *LGR* stayed homogeneous during and after heatwave exposure for all treatments (Figure 5a – Table 2a). During heatwave exposure, *LLR* remained homogeneous with the Control across all treatments. However, pairwise comparisons within each treatment indicated an increase in *LLR* in the +6°C treatment compared to before heatwave exposure (+6°C = −3.07 ± 3.42 mm².d^-1^; Table 2b). After heatwave exposure, *LLR* was homogeneous across all treatment, but showed an overall increase compared to the heatwave period (Figure 5a – Table 2b). During heatwave exposure, *NewLeafGR* increased across all treatments, with the most pronounced rise at +4°C, where it nearly doubles. *OldLeafGR* decreases in all conditions, with the sharpest decline in +4°C treatments (Figure 5b). *The New-Old leaf* ratio increased across treatments, especially in +4°C, indicating a shift toward growth of new leaves over old leaves. After heatwave exposure, *NewLeafGR* remains high or continued increasing, *OldLeafGR* showed a slight increase in some treatments, and the *New-Old leaf* ratio remained elevated, particularly in +4°C (Figure 5b). During heatwave exposure. both gross oxygen production and respiration rates were similar across all treatments (Figure 5c).

#### Ruppia cirrhosa response during the summer experiment

In the controls, 9 shoots died in total (5 during heatwave, 4 after). At +2°C, 7 shoots died (1 during heatwave, 6 after). At +4°C, 12 shoots died (4 before, 6 during heatwave, 2 after). At +6°C, 11 shoots died (3 before, 4 during heatwave, 4 after). During heatwave exposure, Control *LGR* decreased compared to before exposure (2.64± 1.40 mm².d^-1^). *LGR* was homogeneous across treatments, but increased compared to before exposure for the +4°C treatment (3.45 ± 1.90 mm².d^-1^). After heatwave exposure, *LGR* was homogeneous across all treatments, but decreased for +4°C and +6°C treatment compared to during heatwave exposure (Figure 5d – Table 2a). During heatwave exposure, *NewLeafGR* increased, particularly in the +4°C treatment, where it more than tripled compared to before heatwave exposure. In contrast, *OldLeafGR* generally decreased across all treatments. After heatwave exposure, the *New-Old leaf* ratio remained above 1, with *NewLeafGR* staying stable in the +2°C treatment and decreasing in the others. *OldLeafGR* remained stable compared to its levels during heatwave exposure (Figure 5e). During heatwave exposure, both gross oxygen production and respiration rates were similar across all treatments except for the +2°C treatment, where gross oxygen production was significantly lower than the Control (Figure 5f).

## Discussion

### Summary of Species Responses

The results of these two experiments highlight different responses depending on the intensity of the simulated heatwave, the season, and the studied species. *C. nodosa* behaved differently in spring and summer. In spring, this species showed a clear stimulation of *LGR* during all MHW, which coincided with a shift to new leaves growth (increase of the *New-Old leaf* ratio); albeit these positive effects were off-set during the return phase by increased *LLR*. Nevertheless, the species appeared very well adapted to both moderate (i.e., +2°C) and severe (+ 6°C) MHW in spring, by increasing the leaves’ turnover while maintaining high net growth rates. During summer, this species was also well adapted to the MHW, also showing a shift towards new leaves formation, but without an increase of *LGR*. The return phase was again characterized by an increase of leave losses *LLR*, with a concomitant reduction of *LGR*. This was particularly the result of lower *NewLeafGR* than during the previous MHW, which may be related to the decreasing light conditions towards the end of the summer season (see Table 1). *N. noltei* showed more subtle differences between these two seasons. In spring, the +4 °C treatments resulted in a slight stimulation of *LGR* during the MHW, with no effects at +6 °C, while in contrast in summer the +2 °C and +4 °C treatments showed no effect and there was a reduction of *LGR* at + 6 °C. The increase of the *New-Old leaf* ratio during the MHW could compensate for the increased *LLR* during MHW in summer. Hence, *N. noltei* appears very well adapted to the moderate heatwaves both in spring and summer; in spring it can even tolerate an extreme (+ 6°C) MHW while in summer it is more vulnerable to such an extreme (+ 6°C) MHW. *R. cirrhosa* was the only species that showed significant mortality during these experiments and occasionally 34 % of the shoots died in one of the experimental units. On average 48 % of the mortality occurred during a MHW treatment period. Hence MHW could have negative impacts on this species inducing increased mortality, although it is more likely that the experimental conditions were not favourable for certain shoots of this species. In contrast, the remaining shoots of *R. cirrhosa* appeared very well adapted to MHW, as both during spring and summer there was hardly any effect on *LGR*. It’s worth noting that *R. cirrhosa* generally exhibited lower growth rates than the other species, which may be partially attributed to its narrower leaves resulting in less surface area per leaf. Similar to the other species, *R. cirrhosa* showed an increased the *New-Old leaf* ratio during MHW, indicating a shift towards initiation of growth of new leaves.

### Influence of thermal tolerance, seasonal cycles and link with biogeographical distributions

Our study indicates that the three species differed with respect to their sensitivity towards MHW in the following order of increasing vulnerability, i.e. *C. nodosa, R. cirrhosa, N. noltei*. Such differences could be related to lethal thermal thresholds, seasonal patterns and is linked to their biogeographic distribution along latitudinal gradients. The lethal thermal thresholds vary considerably between these three species: up to 40°C for *C. nodosa* according to Bennett et al., (2022), up to 38°C for *R. cirrhosa* according to Tsioli et al., (2019), as well as for *N. noltei* (Massa et al., 2009). This thermal hierarchy largely explains their contrasting responses to the MHW simulated in our study, where *C. nodosa* maintained its growth despite occasional peaks of up to 44°C in summer. MHW do not affect species uniformly across the seasons, but interact with their intrinsic life cycles. Hence, seasonal variations in the response of seagrass to different intensities of simulated marine heatwaves are linked to the natural cycle of this species within the Mediterranean lagoon. In spring, which represents the active growth period for *N. noltei*, growth was stimulated by moderate marine heatwaves (+2°C and +4°C), while extreme marine heatwaves (+6°C) had no effect. This response is in line with the observations of Lee et al., (2007), who demonstrated a positive correlation between temperature and growth in spring, but a negative one in summer. Conversely, *C. nodosa* maintains its growth during spring and summer. *R. cirrhosa* follows a different seasonal pattern, with peak development in late spring and summer, followed by a marked reduction in autumn and virtual disappearance in winter (Mannino & Graziano, 2016).

This difference could explain why *R. cirrhosa*’s growth was hardly impacted during MHWs in summer, simply corresponding to its natural development peak. While occurring ubiquitously in coastal environments, the precise biogeographic distribution is not known (Green & Short, 2003). Both *C. nodosa* and *N. noltei* are Mediterranean species that also occur in the North Atlantic along the coasts of Western Europe and West Africa. However, the distribution of *N. noltei* extends further north than that of *C. nodosa*, while the southern distribution limit of *C. nodosa* is located further south than that of *N. noltei*. Also at the regional scale, *C. nodosa* is widely distributed in Corsican lagoons along the Tyrrhenian Sea (Garrido et al., 2013) and uncommon in the lagoons along the colder Gulf of Lion (Le Fur et al., 2018). Hence, the biogeographic distribution of both species is coherent with their different susceptibilities towards MHW.

### Ecological plasticity of the three species

The three species studied show varying degrees of ecological plasticity, reflecting their respective evolutionary histories and ecological niches. *C. nodosa* stands out for its remarkable plasticity, which enables it not only to tolerate, but sometimes to benefit from high temperatures. This characteristic, combined with its pronounced annual cycle (Agostini et al., 2003; Cancemi et al., 2002), gives it an adaptive advantage in highly variable Mediterranean lagoon environments. As pointed out by Egea et al., (2018), *C. nodosa* may even benefit from future climate scenarios, although this increase in production may be accompanied by a depletion of carbon reserves, making it potentially vulnerable to additional stresses. *R. cirrhosa* reveals a different environmental specialization, favoring hyperhaline environments (Verhoeven, 1975) rather than the Mediterranean Sea itself. This adaptation to extreme conditions may explain its remarkable tolerance to MHW, as our study detected no significant change in its growth or oxygen flux under the experimental conditions. This resilience corroborates the observations of Rasmusson et al., (2021) and suggests that *R. cirrhosa* could maintain its viability in the face of global warming.

It should, however, be highlighted that the abovementioned study by Rasmusson et al., (2021) focused on *R. cirrhosa* in coastal environments along the Baltic Sea in Sweden. This raises the question, whether ecotypes of these species do exist that are genetically adapted to their thermal environments. For instance, one study compared individuals from the Northern Hemisphere (considered cold-adapted) to those from the Southern Hemisphere (considered warm-adapted) and found that the heatwave stimulated the growth of individuals from cold environments while inhibiting the growth of those from warm environments (Nguyen et al., 2021). However, another study compared two Spanish populations of *C. nodosa* from the Mediterranean Sea, separated by 700 km. Due to contrasting summer temperature regimes, one population is considered warm-adapted and the other cold-adapted. The authors found that the cold-environment population was more sensitive to warming, whereas the warm-environment population, for example, increased its growth during the heatwave (Marin-Guirao et al., 2018). Significant interpopulation variability in thermal responses has also been observed for southern populations of *N. noltei* (Massa et al., 2011), which showed a moderate response to heat, while northern populations underwent more marked changes, mainly linked to down-regulation of genes during heat shock.

### Phenological responses to MHW

The systematic increase in the *New-Old leaf* ratio observed in all three species, albeit less pronounced in *R. cirrhosa*, suggests their common phenological response to MHWs. This shift likely reflects an acceleration of natural leaf turnover in response to thermal stress, particularly in *C. nodosa*, a fast-growing opportunistic species where MHW exposure intensifies tissue renewal. The production of new leaves may also involve structural adaptations to heat stress, such as changes in leaf anatomy. For instance, studies on other seagrass species have shown that high temperatures can increase mesophyll tissue thickness (Purnama et al., 2018), a response that, in terrestrial plants, is associated with enhanced plasma membrane permeability—an adaptation that improves gas exchange and heat dissipation while limiting exposure to thermal stress. Additionally, modifications in leaf architecture, particularly the production of smaller leaves, may further optimize gas exchange and heat dissipation, reducing the impact of heat stress. The weaker response observed in *R. cirrhosa* could be attributed to its intrinsic morphology, as its naturally thin leaves require fewer structural adjustments. In *Posidonia oceanica*, young leaves under heat stress halt their growth to redirect energy toward protective mechanisms, notably through the activation of genes such as HSP90, AOX, and BI, which help prevent cellular damage and programmed cell death, thus preserving essential tissues for future survival (Ruocco et al., 2019). This response, combined with an increased production of new leaves, is likely linked to the expression of heat shock proteins (HSP), which enhance thermal tolerance and protect critical cellular functions. Moreover, the activation of photoprotective mechanisms involving heat-responsive genes helps maintain photosynthetic and respiratory activity (Beca-Carretero et al., 2018, 2018; Marin-Guirao et al., 2016). These morpho-physiological adjustments, combined with the species’ ecological strategies—*Ruppia* as a colonizing species, *Nanozostera* as a colonizing/opportunistic species, and *Cymodocea* as an opportunistic species—suggest that species with short life cycles and high regeneration potential exhibit greater resilience to heat stress episodes.

### Post-heatwave responses

Our study also included the post-heatwave period (15 days), as MHW effects often emerge after initial stress. While MHWs can reduce biomass by limiting leaf growth (Deguette et al., 2022), temperature fluctuations in our near-natural system led to a +4°C heatwave in late spring, preventing us from assessing full recovery. Cumulative heat events can negatively affect growth even if individual MHWs do not (Saha et al., 2020), raising concerns as MHWs become more frequent. Initial observations suggest *C. nodosa* can exploit moderate MHWs for growth, maintaining resilience unless extreme heat persists. In contrast, *N. noltei* showed limited capacity to sustain growth under repeated stress, especially in summer, while *R. cirrhosa* remained stable, demonstrating strong thermal resilience.

These findings highlight species-specific adaptation strategies. *C. nodosa* benefits from moderate warming but struggles under prolonged heat, while *N. noltei*’s increased sensitivity suggests a lower resilience threshold, potentially impacting its competitiveness in future climates. *R. cirrhosa*’s ability to maintain stable growth despite repeated stress reinforces its potential for persistence in fluctuating environments. As MHW frequency increases, a key question arises: how will seagrasses respond to multiple consecutive events in short periods, and what are the long-term consequences for ecosystem stability?

### Conclusions and perspectives

Our study investigates for the first time the effect of MHW intensity in spring and summer on three key structuring species in a Mediterranean lagoon, it is very interesting that in our incubation systems, all three species were able to cope with the experimental MHW of 20 days in spring and summer, and only for *N. noltei* a clearly negative impact on growth was observed for the highest intensity (+ 6 °C) in summer. This study supports the importance of considering the provenance of the shoots used in experiments and confirms that organisms inhabiting these lagoons have developed robust survival strategies to withstand more extreme and variable conditions than marine species in more stable ecosystems. With intensification of ECEs, MHWs will become more frequent, our study investigates the effect of one MHWs with different intensities. However, the after period show us the importance of designing studies on the effect of cumulative MHWs on the resilience of seagrass. This is even more relevant for Mediterranean lagoon that are naturally stressed environment.

## Supporting information

Script

data_oxy_summer

data_oxy_spring

metadata

data_temperature_summer

data_temperature_spring

data_growth_summer

data_growth_spring

## Acknowledgements

The authors sincerely thank Nicolas Cimiterra for his support in plant collection, as well as Jean-Yves Coail and Patrick Berriet for their invaluable technical assistance with the creation of the mesocosm system. They also extend their gratitude to Elodie Foucault and Gregory Messiaen for nutrient analysis with special thanks to Gregory Messiaen for his significant contribution to instrumentation. Finally, they thank Elise Lacoste for her help with statistical analyses.

## Author contributions

C.B, RdW, L.S and VO. conceived the study and designed the experimental approach; C.B and E.D/R performed the laboratory analyses; C.B, RdW, L.S, V.O analysed the data and performed the statistical analyses; C.B, RdW writing-original draft preparation; RdW, L.S, V.O writing-review and editing. VO funding acquisition. All authors reviewed the manuscript.

## Data, scripts, code, and supplementary information availability

Data and script are available online on bioRxiv : https://doi.org/10.1101/2025.03.26.645255 ; (Bourdier et al., 2025).

