## Supplementary material for "Experimental approach to assessing the impact of the intensity of near-natural heatwaves on three marine phanerogam species in Mediterranean lagoon environment": Script

````
---
title: "morphometry ; oxygene ; water temperature"
author: "Constance Bourdier"
date: "2025-03-26"
output: pdf
---

#Setup

```{r}

### Définition des packages nécessaires
packages <- c(
  "tidyverse", "lubridate", "emmeans", "lme4", "ggpubr", "FSA",
  "knitr", "kableExtra", "mgcv", "here", "reshape", "car", "gridExtra",
  "png", "gtable", "webshot", "ggsignif", "ggforce", "patchwork",
"ggrepel"
)

### Installation des packages manquants
new_packages <- packages[!packages %in% rownames(installed.packages())]
if (length(new_packages) > 0) install.packages(new_packages)

### Chargement des packages
invisible(lapply(packages, library, character.only = TRUE))

### Paramètres globaux de `knitr`
knitr::opts_chunk$set(
  echo = TRUE,          # Afficher le code
  warning = FALSE,      # Masquer les warnings
  message = FALSE,      # Masquer les messages
  fig.width = 10,       # Largeur des figures plus raisonnable
  fig.height = 6        # Hauteur ajustée
)

```

## Growth responses

## prepare data

```{r}
#| label: Prepare data

charger_donnees_biometrie <- function(saison) {
  if (saison == "ete") {
    fichier <- "biometrie_ete.csv"
    separateur <- ","
  } else if (saison == "printemps") {
    fichier <- "biometrie_printemps2.csv"
    separateur <- ","
  } else {
    stop("Saison non reconnue. Utilisez 'ete' ou 'printemps'.")
  }

  # Chemin vers le répertoire des données
  # Donnees_biometrique <- "~" # add your working directory here
  setwd(Donnees_biometrique)
````
````

````
  # Charger les données
  biometrie <- read.table(fichier, sep = separateur, header = TRUE, dec =
",")

  return(biometrie)
}

biometrie_ete <- charger_donnees_biometrie(saison = "ete")

biometrie_printemps <- charger_donnees_biometrie(saison = "printemps")

```

#### Analysis of SPRING data

```{r}
#| label: SPRING

## Preprocessing and data preparation for seagrass leaf biometry in
spring

biometrie_printemps$DATE_MESURE <- dmy(biometrie_printemps$DATE_MESURE)

biometrie_printemps <- biometrie_printemps %>% mutate(ind =
paste0(ESPECE, REPLICAT, EXPERIENCE, INDIVIDUS, NUMERO_FEUILLE))  ##
combiner les colonnes en une seule
biometrie_printemps <- biometrie_printemps %>% mutate(ind_pieds =
paste0(ESPECE, REPLICAT, EXPERIENCE, INDIVIDUS))  ## combiner les
colonnes en une seule

#### Leaf Surface Area Calculation ####
biometrie_printemps$LARGEUR_FEUILLE_mm <-
as.numeric(biometrie_printemps$LARGEUR_FEUILLE_mm)
biometrie_printemps$LONGUEUR_FEUILLE_mm <-
as.numeric(biometrie_printemps$LONGUEUR_FEUILLE_mm)

biometrie_printemps$ESPECE <- as.factor(biometrie_printemps$ESPECE)
biometrie_printemps$REPLICAT <- as.factor(biometrie_printemps$REPLICAT)
biometrie_printemps$EXPERIENCE <-
as.factor(biometrie_printemps$EXPERIENCE)


## Detailed growth tracking: calculate surface area and time differences
between measurements

biometrie_printemps2 <- biometrie_printemps

biometrie_printemps2$LARGEUR_FEUILLE_mm <-
as.numeric(biometrie_printemps2$LARGEUR_FEUILLE_mm)
biometrie_printemps2[,"SURFACE_FOLIAIRE_MM2"] <-
(biometrie_printemps2[,"LONGUEUR_FEUILLE_mm"])*(biometrie_printemps2[,"LA
RGEUR_FEUILLE_mm"])


  for (n_ind in unique(biometrie_printemps2$ind)) {
    subset_biometrie <- biometrie_printemps2[biometrie_printemps2$ind
%in% n_ind, ]
````

```
    subset_biometrie <-
subset_biometrie[order(subset_biometrie$DATE_MESURE), ]
    if (nrow(subset_biometrie) > 1) {
      for (i in 2:nrow(subset_biometrie)) {
        t0 <- subset_biometrie$DATE_MESURE[i - 1]
        t1 <- subset_biometrie$DATE_MESURE[i]
        biometrie_printemps2[biometrie_printemps2$ind %in% n_ind &
biometrie_printemps2$DATE_MESURE %in% t1, "diff_SF"] <-
          subset_biometrie$SURFACE_FOLIAIRE_MM2[i] -
subset_biometrie$SURFACE_FOLIAIRE_MM2[i - 1]
        biometrie_printemps2[biometrie_printemps2$ind %in% n_ind &
biometrie_printemps2$DATE_MESURE %in% t1, "diff_Temps"] <-
          as.numeric(t1 - t0)
      }
    }
  }

## Correction for new leaves measurements
biometrie_printemps2 <- biometrie_printemps2 %>%
    mutate(TEMPS_MESURE = TEMPS_MESURE %>% str_remove("T") %>%
as.numeric())

## Handle missing values for new leaves
indice_NA <- which(is.na(biometrie_printemps2$diff_SF))
indice_date <- which(biometrie_printemps2$DATE_MESURE > as.Date("2023-04-
21", format = "%Y-%m-%d"))
indice_NA_NF <- intersect(indice_NA, indice_date)

biometrie_printemps2bis <- biometrie_printemps2
biometrie_printemps2bis[indice_NA_NF,"diff_SF"] <-
biometrie_printemps2bis[indice_NA_NF,"SURFACE_FOLIAIRE_MM2"]

## Time difference correction for new leaves
  for (i in indice_NA_NF) {
    TF <- biometrie_printemps2bis[i,"TEMPS_MESURE"]
    DATE_F <- biometrie_printemps2bis[i,"DATE_MESURE"]
    PIEDS <- biometrie_printemps2bis[i,"ind_pieds"]
    indice <- which(biometrie_printemps2bis[,"TEMPS_MESURE"] == (TF-1) &
biometrie_printemps2bis[,"ind_pieds"] == PIEDS)
    if (length(indice) > 0) {
      DATE_I <- unique(biometrie_printemps2bis[indice,"DATE_MESURE"])
      biometrie_printemps2bis[i, "diff_Temps"] <- as.numeric(DATE_F -
DATE_I)
    }
  }

## Calculate net growth rate
biometrie_printemps2bis[,"Taux_croissance_NET"] <-
biometrie_printemps2bis[,"diff_SF"]/biometrie_printemps2bis[,"diff_Temps"
]

## Adjust negative growth rates (cut leaves) to zero
biometrie_printemps2bis <- biometrie_printemps2bis %>%
  mutate(Taux_final = ifelse(Taux_croissance_NET < 0, 0,
Taux_croissance_NET))

## Aggregate net growth rate by individual
```

```
croissance_net_printemps <-
aggregate(biometrie_printemps2bis[,"Taux_final"],
biometrie_printemps2bis[,c("ESPECE","REPLICAT","INDIVIDUS","DATE_MESURE",
"TEMPS_MESURE","EXPERIENCE","ind_pieds")], sum, na.rm = T)
colnames(croissance_net_printemps)[colnames(croissance_net_printemps)==
"x"] <- "Taux_de_croissance_par_pieds"

indice_t0 <- which(croissance_net_printemps[,"TEMPS_MESURE"] == "0")
croissance_net_printemps <- croissance_net_printemps[-indice_t0,]
croissance_net_printemps$SAISON <- "spring"

croissance_net_printemps <- croissance_net_printemps %>%
    mutate(TEMPS_MESURE = str_c("T", TEMPS_MESURE) %>% as.factor())

croissance_net_printemps$ESPECE <-
factor(croissance_net_printemps$ESPECE,
                                levels = c("CN", "ZN", "RC"))

levels(croissance_net_printemps$ESPECE) <- c("Cymodocea nodosa","Zostera
noltei","Ruppia cirrhosa")

## Analysis of new leaf formation
## Aggregate surface area of new leaves by individual and measurement
period
nouvelles_feuille_evol <- biometrie_printemps2bis[indice_NA_NF,]
## Calculate total surface area of new leaves per individual
nouvelles_feuille_evol_pieds <-
aggregate(nouvelles_feuille_evol[,"SURFACE_FOLIAIRE_MM2"],nouvelles_feuil
le_evol[,c("ESPECE","EXPERIENCE","REPLICAT","INDIVIDUS","TEMPS_MESURE","i
nd_pieds")], sum, na.rm = T)
colnames(nouvelles_feuille_evol_pieds)[colnames(nouvelles_feuille_evol_pi
eds)== "x"] <- "somme_new_SF"

nouvelles_feuille_evol_pieds[,"PERIODE"]<-
nouvelles_feuille_evol_pieds[,"TEMPS_MESURE"]

## Create list of individuals by measurement period
nb_pieds <-
unique(biometrie_printemps2bis[,c("ESPECE","EXPERIENCE","REPLICAT","INDIV
IDUS","TEMPS_MESURE", "ind_pieds","diff_Temps")])

## Rename time measurement to period and remove time = 0
nb_pieds[,"PERIODE"]<-  nb_pieds[,"TEMPS_MESURE"]
indice_0 <- which(nb_pieds[,"TEMPS_MESURE"] == 0)
nb_pieds <- nb_pieds[-indice_0,]

## Merge tables of new leaves and all individuals
merge_NF_evol_spring <-
merge(nb_pieds[,c("ESPECE","EXPERIENCE","REPLICAT","INDIVIDUS","ind_pieds
","diff_Temps","PERIODE")],
nouvelles_feuille_evol_pieds[,c("ESPECE","EXPERIENCE","REPLICAT","INDIVID
US","PERIODE","ind_pieds","somme_new_SF")], all.x = T)

## Replace NA with 0 for individuals without new leaves
indice_na <- is.na(merge_NF_evol_spring[,"somme_new_SF"]) == T
merge_NF_evol_spring[indice_na,"somme_new_SF"] = 0

#rendre diff_temps numérique
```

```
merge_NF_evol_spring[,"diff_Temps"] <-
as.numeric(merge_NF_evol_spring[,"diff_Temps"])

## Calculate new leaf formation index (surface area per day)
merge_NF_evol_spring$indice_NF <- (merge_NF_evol_spring$somme_new_SF /
merge_NF_evol_spring$diff_Temps)
merge_NF_evol_spring$PERIODE <- as.factor(merge_NF_evol_spring$PERIODE)

## Calculate mean and standard deviation of new leaf surface area by
species and experiment
mean_NF_spring <-
aggregate(merge_NF_evol_spring[,"somme_new_SF"],merge_NF_evol_spring[,c("
ESPECE","PERIODE","EXPERIENCE")], mean, na.rm = T)
colnames(mean_NF_spring)[colnames(mean_NF_spring)== "x"] <- "mean_NF"

sd_NF_spring <-
aggregate(merge_NF_evol_spring[,"somme_new_SF"],merge_NF_evol_spring[,c("
ESPECE","PERIODE","EXPERIENCE")], mean, na.rm = T)
colnames(sd_NF_spring)[colnames(sd_NF_spring)== "x"] <- "sd_NF"

stat_NF_spring <- merge(mean_NF_spring, sd_NF_spring)

## Function to identify lost leaves between measurement periods
disparues_periode_complet_correction <- function(biometrie_printemps2bis,
temps1, temps2) {
  biometrie_printemps2bis %>%
    group_by(ESPECE, EXPERIENCE, REPLICAT, INDIVIDUS) %>%
    filter(any(TEMPS_MESURE == temps1) & any(TEMPS_MESURE == temps2)) %>%
# Inclure seulement les individus avec données pour les deux temps
    ungroup() %>%
    filter(TEMPS_MESURE %in% c(temps1, temps2)) %>%
    group_by(ESPECE, EXPERIENCE, REPLICAT, INDIVIDUS) %>%
    mutate(feuilles_suivantes = list(NUMERO_FEUILLE[TEMPS_MESURE ==
temps2]),
           diff_Temps = ifelse(TEMPS_MESURE == temps1,
as.numeric(difftime(DATE_MESURE[TEMPS_MESURE == temps2][1],
DATE_MESURE[TEMPS_MESURE == temps1][1], units = "days")), diff_Temps))
%>%
    filter(TEMPS_MESURE == temps1) %>%
    filter(!NUMERO_FEUILLE %in% unlist(feuilles_suivantes)) %>%
    ungroup()
}

## Apply function to identify lost leaves between different time periods
disparues_T0_T1_corrected <-
disparues_periode_complet_correction(biometrie_printemps2bis, 0, 1) %>%
mutate(periode = "T0 à T1")
disparues_T1_T2_corrected <-
disparues_periode_complet_correction(biometrie_printemps2bis, 1, 2) %>%
mutate(periode = "T1 à T2")
disparues_T2_T3_corrected <-
disparues_periode_complet_correction(biometrie_printemps2bis, 2, 3) %>%
mutate(periode = "T2 à T3")

## Combine lost leaves data
tableau_disparues_corrected <- bind_rows(disparues_T0_T1_corrected,
disparues_T1_T2_corrected, disparues_T2_T3_corrected) %>%
```

```
  select(ESPECE, EXPERIENCE, REPLICAT, INDIVIDUS, TEMPS_MESURE, periode,
NUMERO_FEUILLE, ind_pieds, SURFACE_FOLIAIRE_MM2, diff_SF, diff_Temps) %>%
  arrange(ESPECE, EXPERIENCE, REPLICAT, INDIVIDUS, periode)

feuille_perdues_evol_pieds <-
aggregate(tableau_disparues_corrected[,"SURFACE_FOLIAIRE_MM2"],tableau_di
sparues_corrected[,c("ESPECE","EXPERIENCE","REPLICAT","INDIVIDUS","TEMPS_
MESURE","ind_pieds","diff_Temps")], sum, na.rm = T)
colnames(feuille_perdues_evol_pieds)[colnames(feuille_perdues_evol_pieds)
== "SURFACE_FOLIAIRE_MM2"] <- "somme_perte_SF"

feuille_perdues_evol_pieds[,"PERIODE"]<-
feuille_perdues_evol_pieds[,"TEMPS_MESURE"]

# List of plants per period
nb_pieds_perte <-
unique(biometrie_printemps2bis[,c("ESPECE","EXPERIENCE","REPLICAT","INDIV
IDUS","TEMPS_MESURE", "ind_pieds","DATE_MESURE")])

# Rename "TEMPS_MESURE" to "PERIODE" and remove time period 3
nb_pieds_perte[,"PERIODE"]<-  nb_pieds_perte[,"TEMPS_MESURE"]
# Remove time period 3 if needed
# indice_3 <- which(nb_pieds_perte[,"TEMPS_MESURE"] == 3)
# nb_pieds_perte <- nb_pieds_perte[-indice_3,]

# Merge the lost leaves table with the table containing all plants
merge_PF_evol_spring <- merge(nb_pieds_perte,
feuille_perdues_evol_pieds[,c("ESPECE","EXPERIENCE","REPLICAT","INDIVIDUS
","PERIODE","diff_Temps","ind_pieds","somme_perte_SF")], all.x = T)
merge_PF_evol_spring <-
unique(merge_PF_evol_spring[c("ESPECE","REPLICAT","INDIVIDUS","PERIODE","
EXPERIENCE","ind_pieds","somme_perte_SF","diff_Temps")])

indice_T3_perte <- which(merge_PF_evol_spring[,"PERIODE"] == 3)
merge_PF_evol_spring_sansT3 <- merge_PF_evol_spring[-indice_T3_perte,]

# If 'somme_perte_SF' is NA, it means no leaves were lost, so replace
with 0
indice_na <- is.na(merge_PF_evol_spring_sansT3[,"somme_perte_SF"]) == T
merge_PF_evol_spring_sansT3[indice_na,"somme_perte_SF"] = 0

# Make the leaf loss values negative
merge_PF_evol_spring_sansT3$somme_perte_SF <- -
abs(merge_PF_evol_spring_sansT3$somme_perte_SF)

# Calculate the leaf production index (new leaves per day)
merge_PF_evol_spring_sansT3$indice_PF <-
(merge_PF_evol_spring_sansT3$somme_perte_SF /
merge_PF_evol_spring_sansT3$diff_Temps)

merge_PF_evol_spring_sansT3$PERIODE <-
as.factor(merge_PF_evol_spring_sansT3$PERIODE)

# If 'indice_PF' is NA, it means no leaf loss, so replace with 0
indice_na_perte <- is.na(merge_PF_evol_spring_sansT3[,"indice_PF"]) == T
merge_PF_evol_spring_sansT3[indice_na_perte,"indice_PF"] = 0
```

```
#Evolution of the leaf area (SF) for existing leaves between two time
points (mm²/day)

# Create the different data tables (P1, P2, and P3)
### P1 ----
# Select rows where TEMPS_MESURE is less than or equal to 1 (T0 and T1)
indice_P1 <- which(biometrie_printemps2bis[,"TEMPS_MESURE"] <= "1")
P1 <- biometrie_printemps2bis[indice_P1,]

# Modify the leaf area for cut leaves:

# Loop through the rows of the table to detect negative differences
for (i in 2:nrow(P1)) {
  # Check if the leaf area difference is not NA and is negative
  if (!is.na(P1$diff_SF[i]) && P1$diff_SF[i] < 0) {  # If the difference
is negative and not NA
    # Find the T0 value for the same individual and leaf
    T0_value <- P1$SURFACE_FOLIAIRE_MM2[P1$ind == P1$ind[i] &
P1$TEMPS_MESURE == "0"]
    if (length(T0_value) == 1) {  # Check if T0 exists
      # Replace the leaf area value at T1 with the T0 value
      P1$SURFACE_FOLIAIRE_MM2[i] <- T0_value
    }
  }
}


# Identify common leaves between T0 and T1 in P1
communes_num_feuille <- intersect(
  P1$ind[P1$TEMPS_MESURE == "0"],
  P1$ind[P1$TEMPS_MESURE == "1"]
)


# Filter the data to keep only the common leaves (present at T0 and
T1)P1_communes <- P1[P1$ind %in% communes_num_feuille &
P1_communes <- P1[P1$ind %in% communes_num_feuille &
                    P1$TEMPS_MESURE %in% c("0", "1"), ]


# Sum the leaf area (SF) by individual per time point
Somme_SF_ind_P1 <- aggregate(P1_communes[,"SURFACE_FOLIAIRE_MM2"],
P1_communes[,c("ESPECE","REPLICAT","INDIVIDUS","TEMPS_MESURE","EXPERIENCE
","ind_pieds")], sum, na.rm = T)
colnames(Somme_SF_ind_P1)[which(colnames(Somme_SF_ind_P1)== "x")] <-
"Somme_SF_par_individus"

# Merge with P1_communes to get the 'diff_Temps' column
P1_final <-
merge(P1_communes[,c("ESPECE","REPLICAT","INDIVIDUS","TEMPS_MESURE","EXPE
RIENCE","ind_pieds","diff_Temps")],
                   Somme_SF_ind_P1)

P1_final <-
unique(P1_final[c("ESPECE","REPLICAT","INDIVIDUS","TEMPS_MESURE","EXPERIE
NCE","ind_pieds","Somme_SF_par_individus","diff_Temps")])
P1_final[,"TEMPS_MESURE"] <- as.factor(P1_final[,"TEMPS_MESURE"])
```

```
# Add a new empty column for evolution
P1_final$evolution <- NA

# Apply the evolution calculation for each individual
for (ind in unique(P1_final$ind_pieds)) {
  # Filter data for the current individual
  P1_ind <- P1_final[P1_final$ind_pieds == ind, ]

  # Check if both T0 and T1 are present for this individual
  if (all(c("0", "1") %in% P1_ind$TEMPS_MESURE)) {
    # Filter data to get the T0 and T1 values
    T0 <- P1_ind[P1_ind$TEMPS_MESURE == "0", "Somme_SF_par_individus"]
    T1 <- P1_ind[P1_ind$TEMPS_MESURE == "1", "Somme_SF_par_individus"]

    # Calculate the evolution (difference between T1 and T0)
    evolution <- (T1 - T0)

    # Add the evolution to the dataframe for the T1 rows of the
individual
    P1_final$evolution[P1_final$ind_pieds == ind & P1_final$TEMPS_MESURE
== "1"] <- evolution
  }
}

# Divide by the number of days to get evolution per day:
P1_final$evolution_jour <- NA
P1_final[,"evolution_jour"] <-
P1_final[,"evolution"]/P1_final[,"diff_Temps"]

# Remove T0 data (we only want the data for T1 and onwards)
indice_T0 <- which(P1_final[,"TEMPS_MESURE"] == "0")
P1_SPRING <- P1_final[-indice_T0,]  # Remove T0 rows

### P2 ----
# Select rows where TEMPS_MESURE is either 1 or 2 (T1 and T2)
indice_P2 <- which(biometrie_printemps2bis[,"TEMPS_MESURE"] == "1" |
biometrie_printemps2bis[,"TEMPS_MESURE"] == "2")
P2 <- biometrie_printemps2bis[indice_P2,]

# Modify the leaf area for cut leaves:

# Loop through the rows of the table to detect negative differences
for (i in 2:nrow(P2)) {
  # Check if the leaf area difference is not NA and is negative
  if (!is.na(P2$diff_SF[i]) && P2$diff_SF[i] < 0) {  # If the difference
is negative and not NA
    # Find the T1 value for the same individual and leaf
    T1_value <- P2$SURFACE_FOLIAIRE_MM2[P2$ind == P2$ind[i] &
P2$TEMPS_MESURE == "1"]
    if (length(T1_value) == 1) {  # Check if T1 exists
      # Replace the leaf area value at T1 with the T1 value
      P2$SURFACE_FOLIAIRE_MM2[i] <- T1_value
    }
  }
}

# Identify common leaves between T1 and T2 in P2
```

```
# Find the leaf numbers present at both T1 and T2
communes_num_feuille <- intersect(
  P2$ind[P2$TEMPS_MESURE == "1"],
  P2$ind[P2$TEMPS_MESURE == "2"]
)

# Filter the data to keep only the common leaves (present at T1 and T2)
P2_communes <- P2[P2$ind %in% communes_num_feuille &
                    P2$TEMPS_MESURE %in% c("1", "2"), ]

# Sum the leaf area (SF) by individual per time point
Somme_SF_ind_P2 <- aggregate(P2_communes[,"SURFACE_FOLIAIRE_MM2"],
P2_communes[,c("ESPECE","REPLICAT","INDIVIDUS","TEMPS_MESURE","EXPERIENCE
","ind_pieds")], sum, na.rm = T)
colnames(Somme_SF_ind_P2)[which(colnames(Somme_SF_ind_P2)== "x")] <-
"Somme_SF_par_individus"  # Rename the column to 'Somme_SF_par_individus'

# Merge with P2_communes to get the 'diff_Temps' column
P2_final <-
merge(P2_communes[c("ESPECE","REPLICAT","TEMPS_MESURE","EXPERIENCE","ind_
pieds","diff_Temps")],

Somme_SF_ind_P2[c("ESPECE","REPLICAT","INDIVIDUS","TEMPS_MESURE","EXPERIE
NCE","ind_pieds","Somme_SF_par_individus")])

# Remove duplicates caused by multiple leaves (for the same individual)
per time point
P2_final <-
unique(P2_final[c("ESPECE","REPLICAT","INDIVIDUS","TEMPS_MESURE","EXPERIE
NCE","ind_pieds","Somme_SF_par_individus","diff_Temps")])

P2_final[,"TEMPS_MESURE"] <- as.factor(P2_final[,"TEMPS_MESURE"])

# Add a new empty column for evolution
P2_final$evolution <- NA

# Apply the evolution calculation for each individual
for (ind in unique(P2_final$ind_pieds)) {
  # Filter data for the current individual
  P2_ind <- P2_final[P2_final$ind_pieds == ind, ]

  # Check if both T1 and T2 are present for this individual
  if (all(c("1", "2") %in% P2_ind$TEMPS_MESURE)) {
    # Filter data to get the T1 and T2 values
    T1 <- P2_ind[P2_ind$TEMPS_MESURE == "1", "Somme_SF_par_individus"]
    T2 <- P2_ind[P2_ind$TEMPS_MESURE == "2", "Somme_SF_par_individus"]

    # Calculate the evolution (difference between T2 and T1)
    evolution <- (T2 - T1)

    # Add the evolution to the dataframe for the T2 rows of the
individual
    P2_final$evolution[P2_final$ind_pieds == ind & P2_final$TEMPS_MESURE
== "2"] <- evolution
  }
}

# Divide by the number of days to get evolution per day:
```

```
P2_final$evolution_jour <- NA
P2_final[,"evolution_jour"] <-
P2_final[,"evolution"]/P2_final[,"diff_Temps"]

# Remove T1 data (we only want the data for T2 and onwards)
indice_T1 <- which(P2_final[,"TEMPS_MESURE"] == "1")
P2_SPRING <- P2_final[-indice_T1,]  # Remove T1 rows


###### P3 #####

# Select rows where TEMPS_MESURE is either 2 or 3 (T2 and T3)
indice_P3 <- which(biometrie_printemps2bis[,"TEMPS_MESURE"] == "2" |
biometrie_printemps2bis[,"TEMPS_MESURE"] == "3")
P3 <- biometrie_printemps2bis[indice_P3,]

# Modify the leaf area for cut leaves:

# Loop through the rows of the table to detect negative differences
for (i in 2:nrow(P3)) {
  # Check if the leaf area difference is not NA and is negative
  if (!is.na(P3$diff_SF[i]) && P3$diff_SF[i] < 0) {  # If the difference
is negative and not NA
    # Find the T2 value for the same individual and leaf
    T2_value <- P3$SURFACE_FOLIAIRE_MM2[P3$ind == P3$ind[i] &
P3$TEMPS_MESURE == "2"]
    if (length(T2_value) == 1) {  # Check if T2 exists
      # Replace the leaf area value at T3 with the T2 value
      P3$SURFACE_FOLIAIRE_MM2[i] <- T2_value
    }
  }
}

# Identify common leaves between T2 and T3 in P3

# Find the leaf numbers present at both T2 and T3
communes_num_feuille <- intersect(
  P3$ind[P3$TEMPS_MESURE == "2"],
  P3$ind[P3$TEMPS_MESURE == "3"]
)

# Filter the data to keep only the common leaves (present at T2 and T3)
P3_communes <- P3[P3$ind %in% communes_num_feuille &
                    P3$TEMPS_MESURE %in% c("2", "3"), ]

# Sum the leaf area (SF) by individual per time point
Somme_SF_ind_P3 <- aggregate(P3_communes[,"SURFACE_FOLIAIRE_MM2"],
P3_communes[,c("ESPECE","REPLICAT","INDIVIDUS","TEMPS_MESURE","EXPERIENCE
","ind_pieds")], sum, na.rm = T)
colnames(Somme_SF_ind_P3)[which(colnames(Somme_SF_ind_P3)== "x")] <-
"Somme_SF_par_individus"

# Merge with P3_communes to get the 'diff_Temps' column
P3_final <-
merge(P3_communes[c("ESPECE","REPLICAT","TEMPS_MESURE","EXPERIENCE","ind_
pieds","diff_Temps")],
```

```
Somme_SF_ind_P3[c("ESPECE","REPLICAT","INDIVIDUS","TEMPS_MESURE","EXPERIE
NCE","ind_pieds","Somme_SF_par_individus")])

# Remove duplicates caused by multiple leaves (for the same individual)
per time point
P3_final <-
unique(P3_final[c("ESPECE","REPLICAT","INDIVIDUS","TEMPS_MESURE","EXPERIE
NCE","ind_pieds","Somme_SF_par_individus","diff_Temps")])

P3_final[,"TEMPS_MESURE"] <- as.factor(P3_final[,"TEMPS_MESURE"])

# Add a new empty column for evolution
P3_final$evolution <- NA

# Apply the evolution calculation for each individual
for (ind in unique(P3_final$ind_pieds)) {
  # Filter data for the current individual
  P3_ind <- P3_final[P3_final$ind_pieds == ind, ]

  # Check if both T2 and T3 are present for this individual
  if (all(c("2", "3") %in% P3_ind$TEMPS_MESURE)) {
    # Filter data to get the T2 and T3 values
    T2 <- P3_ind[P3_ind$TEMPS_MESURE == "2", "Somme_SF_par_individus"]
    T3 <- P3_ind[P3_ind$TEMPS_MESURE == "3", "Somme_SF_par_individus"]

    # Calculate the evolution (difference between T3 and T2)
    evolution <- (T3 - T2)

    # Add the evolution to the dataframe for the T3 rows of the
individual
    P3_final$evolution[P3_final$ind_pieds == ind & P3_final$TEMPS_MESURE
== "3"] <- evolution
  }
}

# Divide by the number of days to get evolution per day:
P3_final$evolution_jour <- NA
P3_final[,"evolution_jour"] <-
P3_final[,"evolution"]/P3_final[,"diff_Temps"]

# Remove T2 data (we only want the data for T3 and onwards)
indice_T2 <- which(P3_final[,"TEMPS_MESURE"] == "2")
P3_SPRING <- P3_final[-indice_T2,]

### Combine the three datasets (P1, P2, P3)
EVOL_SPRING <- rbind(P1_SPRING, P2_SPRING, P3_SPRING)

```

#### Analysis of SUMMER data

```{r}
#| label: SUMMER
#|
## Preprocessing and data preparation for seagrass leaf biometry in
summer
biometrie_ete$DATE_MESURE <- dmy(biometrie_ete$DATE_MESURE)
```

```
biometrie_ete <- biometrie_ete %>% mutate(ind = paste0(ESPECE, REPLICAT,
EXPERIENCE, INDIVIDUS, NUMERO_FEUILLE))  ## combiner les colonnes en une
seule
biometrie_ete <- biometrie_ete %>% mutate(ind_pieds = paste0(ESPECE,
REPLICAT, EXPERIENCE, INDIVIDUS))  ## combiner les colonnes en une seule

#### Leaf Surface Area Calculation ####
biometrie_ete$LARGEUR_FEUILLE_mm <-
as.numeric(biometrie_ete$LARGEUR_FEUILLE_mm)
biometrie_ete$LONGUEUR_FEUILLE_mm <-
as.numeric(biometrie_ete$LONGUEUR_FEUILLE_mm)

biometrie_ete$ESPECE <- as.factor(biometrie_ete$ESPECE)
biometrie_ete$REPLICAT <- as.factor(biometrie_ete$REPLICAT)
biometrie_ete$EXPERIENCE <- as.factor(biometrie_ete$EXPERIENCE)

# Différence de surface foliaire par feuille entre deux temps de mesures
(diff SF et diff temps)

biometrie_ete2 <- biometrie_ete

biometrie_ete2$LARGEUR_FEUILLE_mm <-
as.numeric(biometrie_ete2$LARGEUR_FEUILLE_mm)
biometrie_ete2[,"SURFACE_FOLIAIRE_MM2"] <-
(biometrie_ete2[,"LONGUEUR_FEUILLE_mm"])*(biometrie_ete2[,"LARGEUR_FEUILL
E_mm"])

## Detailed growth tracking: calculate surface area and time differences
between measurements
 for (n_ind in unique(biometrie_ete2$ind)) {
    subset_biometrie <- biometrie_ete2[biometrie_ete2$ind %in% n_ind, ]
    subset_biometrie <-
subset_biometrie[order(subset_biometrie$DATE_MESURE), ]
    if (nrow(subset_biometrie) > 1) {
      for (i in 2:nrow(subset_biometrie)) {
        t0 <- subset_biometrie$DATE_MESURE[i - 1]
        t1 <- subset_biometrie$DATE_MESURE[i]
        biometrie_ete2[biometrie_ete2$ind %in% n_ind &
biometrie_ete2$DATE_MESURE %in% t1, "diff_SF"] <-
          subset_biometrie$SURFACE_FOLIAIRE_MM2[i] -
subset_biometrie$SURFACE_FOLIAIRE_MM2[i - 1]
        biometrie_ete2[biometrie_ete2$ind %in% n_ind &
biometrie_ete2$DATE_MESURE %in% t1, "diff_Temps"] <-
          as.numeric(t1 - t0)
      }
    }
  }

## Correction for new leaves measurements

biometrie_ete2 <- biometrie_ete2 %>%
    mutate(TEMPS_MESURE = TEMPS_MESURE %>% str_remove("T") %>%
as.numeric())

## Handle missing values for new leaves
indice_NA_ete <- which(is.na(biometrie_ete2$diff_SF))
indice_date_ete <- which(biometrie_ete2$DATE_MESURE > as.Date("2023-07-
21", format = "%Y-%m-%d"))
```

```
indice_NA_NF_ete <- intersect(indice_NA_ete, indice_date_ete)

biometrie_ete2bis <- biometrie_ete2
biometrie_ete2bis[indice_NA_NF_ete,"diff_SF"] <-
biometrie_ete2bis[indice_NA_NF_ete,"SURFACE_FOLIAIRE_MM2"]

## Time difference correction for new leaves
  for (i in indice_NA_NF_ete) {
    TF <- biometrie_ete2bis[i,"TEMPS_MESURE"]
    DATE_F <- biometrie_ete2bis[i,"DATE_MESURE"]
    PIEDS <- biometrie_ete2bis[i,"ind_pieds"]
    indice <- which(biometrie_ete2bis[,"TEMPS_MESURE"] == (TF-1) &
biometrie_ete2bis[,"ind_pieds"] == PIEDS)
    if (length(indice) > 0) {
      DATE_I <- unique(biometrie_ete2bis[indice,"DATE_MESURE"])
      biometrie_ete2bis[i, "diff_Temps"] <- as.numeric(DATE_F - DATE_I)
    }
  }

## Calculate net growth rate
biometrie_ete2bis[,"Taux_croissance_NET"] <-
biometrie_ete2bis[,"diff_SF"]/biometrie_ete2bis[,"diff_Temps"]

## Adjust negative growth rates (cut leaves) to zero
biometrie_ete2bis <- biometrie_ete2bis %>%
  mutate(Taux_final = ifelse(Taux_croissance_NET < 0, 0,
Taux_croissance_NET))


## Aggregate net growth rate by individual
croissance_net <- aggregate(biometrie_ete2bis[,"Taux_final"],
biometrie_ete2bis[,c("ESPECE","REPLICAT","INDIVIDUS","DATE_MESURE","TEMPS
_MESURE","EXPERIENCE","ind_pieds")], sum, na.rm = T)
colnames(croissance_net)[colnames(croissance_net)== "x"] <-
"Taux_de_croissance_par_pieds"


indice_t0 <- which(croissance_net[,"TEMPS_MESURE"] == "0")
croissance_net <- croissance_net[-indice_t0,]
croissance_net$SAISON <- "summer"
croissance_net <- croissance_net %>%
    mutate(TEMPS_MESURE = str_c("T", TEMPS_MESURE) %>% as.factor())
croissance_net$ESPECE <- factor(croissance_net$ESPECE,
                                levels = c("CN", "ZN", "RC"))

# Changer les noms des niveaux de ESPECE
levels(croissance_net$ESPECE) <- c("Cymodocea nodosa","Zostera
noltei","Ruppia cirrhosa")


## Analysis of new leaf formation
## Aggregate surface area of new leaves by individual and measurement
period
nouvelles_feuille_evol_ete <- biometrie_ete2bis[indice_NA_NF_ete,]
## Calculate total surface area of new leaves per individual
nouvelles_feuille_evol_pieds_ete <-
aggregate(nouvelles_feuille_evol_ete[,"SURFACE_FOLIAIRE_MM2"],nouvelles_f
```

```
euille_evol_ete[,c("ESPECE","EXPERIENCE","REPLICAT","INDIVIDUS","TEMPS_ME
SURE","ind_pieds")], sum, na.rm = T)
colnames(nouvelles_feuille_evol_pieds_ete)[colnames(nouvelles_feuille_evo
l_pieds_ete)== "x"] <- "somme_new_SF"

nouvelles_feuille_evol_pieds_ete[,"PERIODE"]<-
nouvelles_feuille_evol_pieds_ete[,"TEMPS_MESURE"]

## Create list of individuals by measurement period
test_pieds_ete <-
unique(biometrie_ete2bis[,c("ESPECE","EXPERIENCE","REPLICAT","INDIVIDUS",
"TEMPS_MESURE", "ind_pieds","diff_Temps")])

## Rename time measurement to period and remove time = 0
test_pieds_ete[,"PERIODE"]<-  test_pieds_ete[,"TEMPS_MESURE"]
indice_0 <- which(test_pieds_ete[,"TEMPS_MESURE"] == 0)
test_pieds_ete <- test_pieds_ete[-indice_0,]


## Merge tables of new leaves and all individuals
merge_NF_evol_summer <-
merge(test_pieds_ete[,c("ESPECE","EXPERIENCE","REPLICAT","INDIVIDUS","ind
_pieds","diff_Temps","PERIODE")],
nouvelles_feuille_evol_pieds_ete[,c("ESPECE","EXPERIENCE","REPLICAT","IND
IVIDUS","PERIODE","ind_pieds","somme_new_SF")], all.x = T)

## Replace NA with 0 for individuals without new leaves
indice_na_ete <- is.na(merge_NF_evol_summer[,"somme_new_SF"]) == T
merge_NF_evol_summer[indice_na_ete,"somme_new_SF"] = 0
merge_NF_evol_summer[,"diff_Temps"] <-
as.numeric(merge_NF_evol_summer[,"diff_Temps"])

## Calculate new leaf formation index (surface area per day)
merge_NF_evol_summer$indice_NF <- (merge_NF_evol_summer$somme_new_SF /
merge_NF_evol_summer$diff_Temps)
merge_NF_evol_summer$PERIODE <- as.factor(merge_NF_evol_summer$PERIODE)

mean_NF_summer <-
aggregate(merge_NF_evol_summer[,"somme_new_SF"],merge_NF_evol_summer[,c("
ESPECE","PERIODE","EXPERIENCE")], mean, na.rm = T)
colnames(mean_NF_summer)[colnames(mean_NF_summer)== "x"] <- "mean_NF"

sd_NF_summer <-
aggregate(merge_NF_evol_summer[,"somme_new_SF"],merge_NF_evol_summer[,c("
ESPECE","PERIODE","EXPERIENCE")], mean, na.rm = T)
colnames(sd_NF_summer)[colnames(sd_NF_summer)== "x"] <- "sd_NF"

stat_NF_summer <- merge(mean_NF_summer, sd_NF_summer)

## Function to identify lost leaves between measurement periods
disparues_periode_complet_ete <- function(biometrie_ete2bis, temps1,
temps2) {
  biometrie_ete2bis %>%
    group_by(ESPECE, EXPERIENCE, REPLICAT, INDIVIDUS) %>%
    filter(any(TEMPS_MESURE == temps1) & any(TEMPS_MESURE == temps2)) %>%
# Inclure seulement les individus avec données pour les deux temps
    ungroup() %>%
    filter(TEMPS_MESURE %in% c(temps1, temps2)) %>%
```

```
    group_by(ESPECE, EXPERIENCE, REPLICAT, INDIVIDUS) %>%
    mutate(feuilles_suivantes = list(NUMERO_FEUILLE[TEMPS_MESURE ==
temps2]),
           diff_Temps = ifelse(TEMPS_MESURE == temps1,
as.numeric(difftime(DATE_MESURE[TEMPS_MESURE == temps2][1],
DATE_MESURE[TEMPS_MESURE == temps1][1], units = "days")), diff_Temps))
%>%
    filter(TEMPS_MESURE == temps1) %>%
    filter(!NUMERO_FEUILLE %in% unlist(feuilles_suivantes)) %>%
    ungroup()
}

## Apply function to identify lost leaves between different time periods
disparues_T0_T1_corrected_ete <-
disparues_periode_complet_ete(biometrie_ete2bis, 0, 1) %>% mutate(periode
= "T0 à T1")
disparues_T1_T2_corrected_ete <-
disparues_periode_complet_ete(biometrie_ete2bis, 1, 2) %>% mutate(periode
= "T1 à T2")
disparues_T2_T3_corrected_ete <-
disparues_periode_complet_ete(biometrie_ete2bis, 2, 3) %>% mutate(periode
= "T2 à T3")

## Combine lost leaves data
tableau_disparues_corrected_ete <-
bind_rows(disparues_T0_T1_corrected_ete, disparues_T1_T2_corrected_ete,
disparues_T2_T3_corrected_ete) %>%
  select(ESPECE, EXPERIENCE, REPLICAT, INDIVIDUS, TEMPS_MESURE, periode,
NUMERO_FEUILLE, ind_pieds, SURFACE_FOLIAIRE_MM2, diff_SF, diff_Temps) %>%
  arrange(ESPECE, EXPERIENCE, REPLICAT, INDIVIDUS, periode)

feuille_perdues_evol_pieds_ete <-
aggregate(tableau_disparues_corrected_ete[,"SURFACE_FOLIAIRE_MM2"],tablea
u_disparues_corrected_ete[,c("ESPECE","EXPERIENCE","REPLICAT","INDIVIDUS"
,"TEMPS_MESURE","ind_pieds","diff_Temps")], sum, na.rm = T)
colnames(feuille_perdues_evol_pieds_ete)[colnames(feuille_perdues_evol_pi
eds_ete)== "SURFACE_FOLIAIRE_MM2"] <- "somme_perte_SF"

# Change "TEMPS_MESURE" to "PERIODE" and remove time period 3
feuille_perdues_evol_pieds_ete[,"PERIODE"]<-
feuille_perdues_evol_pieds_ete[,"TEMPS_MESURE"]

# List of plants per period
test_pieds_perte_ete <-
unique(biometrie_ete2bis[,c("ESPECE","EXPERIENCE","REPLICAT","INDIVIDUS",
"TEMPS_MESURE", "ind_pieds","DATE_MESURE")])

test_pieds_perte_ete[,"PERIODE"]<-  test_pieds_perte_ete[,"TEMPS_MESURE"]

# Merge the lost leaves table with the table containing all plants
merge_PF_evol_summer <- merge(test_pieds_perte_ete,
feuille_perdues_evol_pieds_ete[,c("ESPECE","EXPERIENCE","REPLICAT","INDIV
IDUS","PERIODE","diff_Temps","ind_pieds","somme_perte_SF")], all.x = T)


indice_T3_perte <- which(merge_PF_evol_summer[,"PERIODE"] == 3)
merge_PF_evol_summer_sansT3 <- merge_PF_evol_summer[-indice_T3_perte,]
```

```
# If 'somme_perte_SF' is NA, it means no leaves were lost, so replace
with 0
indice_na_ete <- is.na(merge_PF_evol_summer_sansT3[,"somme_perte_SF"]) ==
T
merge_PF_evol_summer_sansT3[indice_na,"somme_perte_SF"] = 0

# Make the leaf loss values negative
merge_PF_evol_summer_sansT3$somme_perte_SF <- -
abs(merge_PF_evol_summer_sansT3$somme_perte_SF)

# Calculate the leaf production index (new leaves per day)
merge_PF_evol_summer_sansT3$indice_PF <-
(merge_PF_evol_summer_sansT3$somme_perte_SF /
merge_PF_evol_summer_sansT3$diff_Temps)

merge_PF_evol_summer_sansT3$PERIODE <-
as.factor(merge_PF_evol_summer_sansT3$PERIODE)

# If 'indice_PF' is NA, it means no leaf loss, so replace with 0
indice_na_perte <- is.na(merge_PF_evol_summer_sansT3[,"indice_PF"]) == T
merge_PF_evol_summer_sansT3[indice_na_perte,"indice_PF"] = 0


#Evolution of the leaf area (SF) for existing leaves between two time
points (mm²/day)

# Create the different data tables (P1, P2, and P3)
### P1 ----
# Select rows where TEMPS_MESURE is less than or equal to 1 (T0 and T1)
indice_P1_ete <- which(biometrie_ete2bis[,"TEMPS_MESURE"] <= "1")
P1_ete <- biometrie_ete2bis[indice_P1_ete,]

for (i in 2:nrow(P1_ete)) {
  if (!is.na(P1_ete$diff_SF[i]) && P1_ete$diff_SF[i] < 0) {
    T0_value <- P1_ete$SURFACE_FOLIAIRE_MM2[P1_ete$ind == P1_ete$ind[i] &
P1_ete$TEMPS_MESURE == "0"]
    if (length(T0_value) == 1) {
      P1_ete$SURFACE_FOLIAIRE_MM2[i] <- T0_value
    }
  }
}

communes_num_feuille <- intersect(
  P1_ete$ind[P1_ete$TEMPS_MESURE == "0"],
  P1_ete$ind[P1_ete$TEMPS_MESURE == "1"]
)

P1_ete_communes <- P1_ete[P1_ete$ind %in% communes_num_feuille &
                    P1_ete$TEMPS_MESURE %in% c("0", "1"), ]


Somme_SF_ind_P1_ete <-
aggregate(P1_ete_communes[,"SURFACE_FOLIAIRE_MM2"],
P1_ete_communes[,c("ESPECE","REPLICAT","INDIVIDUS","TEMPS_MESURE","EXPERI
ENCE","ind_pieds")], sum, na.rm = T)
colnames(Somme_SF_ind_P1_ete)[which(colnames(Somme_SF_ind_P1_ete)== "x")]
<- "Somme_SF_par_individus"
```

```
P1_ete_final <-
merge(P1_ete_communes[,c("ESPECE","REPLICAT","TEMPS_MESURE","EXPERIENCE",
"ind_pieds","diff_Temps")],
                   Somme_SF_ind_P1_ete)
P1_ete_final <-
unique(P1_ete_final[c("ESPECE","REPLICAT","INDIVIDUS","TEMPS_MESURE","EXP
ERIENCE","ind_pieds","Somme_SF_par_individus","diff_Temps")])

P1_ete_final[,"TEMPS_MESURE"] <- as.factor(P1_ete_final[,"TEMPS_MESURE"])


P1_ete_final$evolution <- NA

for (ind in unique(P1_ete_final$ind_pieds)) {
  P1_ete_ind <- P1_ete_final[P1_ete_final$ind_pieds == ind, ]
  if (all(c("0", "1") %in% P1_ete_ind$TEMPS_MESURE)) {
    T0 <- P1_ete_ind[P1_ete_ind$TEMPS_MESURE == "0",
"Somme_SF_par_individus"]
    T1 <- P1_ete_ind[P1_ete_ind$TEMPS_MESURE == "1",
"Somme_SF_par_individus"]
    evolution <- (T1 - T0)
    P1_ete_final$evolution[P1_ete_final$ind_pieds == ind &
P1_ete_final$TEMPS_MESURE == "1"] <- evolution
  }
}

P1_ete_final$evolution_jour <- NA
P1_ete_final[,"evolution_jour"] <-
P1_ete_final[,"evolution"]/P1_ete_final[,"diff_Temps"]


indice_T0_ete <- which(P1_ete_final[,"TEMPS_MESURE"] == "0")
P1_ete <- P1_ete_final[-indice_T0_ete,]

### P2 ----
indice_P2_ete <- which(biometrie_ete2bis[,"TEMPS_MESURE"] == "1" |
biometrie_ete2bis[,"TEMPS_MESURE"] == "2")
P2_ete <- biometrie_ete2bis[indice_P2_ete,]

for (i in 2:nrow(P2_ete)) {

  if (!is.na(P2_ete$diff_SF[i]) && P2_ete$diff_SF[i] < 0) {
    T1_value <- P2_ete$SURFACE_FOLIAIRE_MM2[P2_ete$ind == P2_ete$ind[i] &
P2_ete$TEMPS_MESURE == "1"]
    if (length(T1_value) == 1) {
      P2_ete$SURFACE_FOLIAIRE_MM2[i] <- T1_value
    }
  }
}

communes_num_feuille <- intersect(
  P2_ete$ind[P2_ete$TEMPS_MESURE == "1"],
  P2_ete$ind[P2_ete$TEMPS_MESURE == "2"]
)

P2_ete_communes <- P2_ete[P2_ete$ind %in% communes_num_feuille &
                    P2_ete$TEMPS_MESURE %in% c("1", "2"), ]
```

```
Somme_SF_ind_P2_ete <-
aggregate(P2_ete_communes[,"SURFACE_FOLIAIRE_MM2"],
P2_ete_communes[,c("ESPECE","REPLICAT","INDIVIDUS","TEMPS_MESURE","EXPERI
ENCE","ind_pieds")], sum, na.rm = T)
colnames(Somme_SF_ind_P2_ete)[which(colnames(Somme_SF_ind_P2_ete)== "x")]
<- "Somme_SF_par_individus"

P2_ete_final <-
merge(P2_ete_communes[c("ESPECE","REPLICAT","TEMPS_MESURE","EXPERIENCE","
ind_pieds","diff_Temps")],

Somme_SF_ind_P2_ete[c("ESPECE","REPLICAT","INDIVIDUS","TEMPS_MESURE","EXP
ERIENCE","ind_pieds","Somme_SF_par_individus")])

P2_ete_final <-
unique(P2_ete_final[c("ESPECE","REPLICAT","INDIVIDUS","TEMPS_MESURE","EXP
ERIENCE","ind_pieds","Somme_SF_par_individus","diff_Temps")])

P2_ete_final[,"TEMPS_MESURE"] <- as.factor(P2_ete_final[,"TEMPS_MESURE"])

P2_ete_final$evolution <- NA

for (ind in unique(P2_ete_final$ind_pieds)) {
  P2_ete_ind <- P2_ete_final[P2_ete_final$ind_pieds == ind, ]
  if (all(c("1", "2") %in% P2_ete_ind$TEMPS_MESURE)) {
    T1 <- P2_ete_ind[P2_ete_ind$TEMPS_MESURE == "1",
"Somme_SF_par_individus"]
    T2 <- P2_ete_ind[P2_ete_ind$TEMPS_MESURE == "2",
"Somme_SF_par_individus"]
    evolution <- (T2 - T1)
    P2_ete_final$evolution[P2_ete_final$ind_pieds == ind &
P2_ete_final$TEMPS_MESURE == "2"] <- evolution
  }
}

P2_ete_final$evolution_jour <- NA
P2_ete_final[,"evolution_jour"] <-
P2_ete_final[,"evolution"]/P2_ete_final[,"diff_Temps"]

indice_T1_ete <- which(P2_ete_final[,"TEMPS_MESURE"] == "1")
P2_ete <- P2_ete_final[-indice_T1_ete,]

###### P3 ----

indice_P3_ete <- which(biometrie_ete2bis[,"TEMPS_MESURE"] == "2" |
biometrie_ete2bis[,"TEMPS_MESURE"] == "3")
P3_ete <- biometrie_ete2bis[indice_P3_ete,]
for (i in 2:nrow(P3_ete)) {
  if (!is.na(P3_ete$diff_SF[i]) && P3_ete$diff_SF[i] < 0) {
    T2_value <- P3_ete$SURFACE_FOLIAIRE_MM2[P3_ete$ind == P3_ete$ind[i] &
P3_ete$TEMPS_MESURE == "2"]
    if (length(T2_value) == 1) {
      P3_ete$SURFACE_FOLIAIRE_MM2[i] <- T2_value
    }
  }
}
```

```
communes_num_feuille <- intersect(
  P3_ete$ind[P3_ete$TEMPS_MESURE == "2"],
  P3_ete$ind[P3_ete$TEMPS_MESURE == "3"]
)

P3_ete_communes <- P3_ete[P3_ete$ind %in% communes_num_feuille &
                    P3_ete$TEMPS_MESURE %in% c("2", "3"), ]


Somme_SF_ind_P3_ete <-
aggregate(P3_ete_communes[,"SURFACE_FOLIAIRE_MM2"],
P3_ete_communes[,c("ESPECE","REPLICAT","INDIVIDUS","TEMPS_MESURE","EXPERI
ENCE","ind_pieds")], sum, na.rm = T)
colnames(Somme_SF_ind_P3_ete)[which(colnames(Somme_SF_ind_P3_ete)== "x")]
<- "Somme_SF_par_individus"

P3_ete_final <-
merge(P3_ete_communes[c("ESPECE","REPLICAT","TEMPS_MESURE","EXPERIENCE","
ind_pieds","diff_Temps")],

Somme_SF_ind_P3_ete[c("ESPECE","REPLICAT","INDIVIDUS","TEMPS_MESURE","EXP
ERIENCE","ind_pieds","Somme_SF_par_individus")])

P3_ete_final <-
unique(P3_ete_final[c("ESPECE","REPLICAT","INDIVIDUS","TEMPS_MESURE","EXP
ERIENCE","ind_pieds","Somme_SF_par_individus","diff_Temps")])

P3_ete_final[,"TEMPS_MESURE"] <- as.factor(P3_ete_final[,"TEMPS_MESURE"])

P3_ete_final$evolution <- NA

for (ind in unique(P3_ete_final$ind_pieds)) {
  P3_ete_ind <- P3_ete_final[P3_ete_final$ind_pieds == ind, ]
  if (all(c("2", "3") %in% P3_ete_ind$TEMPS_MESURE)) {
    T2 <- P3_ete_ind[P3_ete_ind$TEMPS_MESURE == "2",
"Somme_SF_par_individus"]
    T3 <- P3_ete_ind[P3_ete_ind$TEMPS_MESURE == "3",
"Somme_SF_par_individus"]
    evolution <- (T3 - T2)
    P3_ete_final$evolution[P3_ete_final$ind_pieds == ind &
P3_ete_final$TEMPS_MESURE == "3"] <- evolution
  }
}

P3_ete_final$evolution_jour <- NA
P3_ete_final[,"evolution_jour"] <-
P3_ete_final[,"evolution"]/P3_ete_final[,"diff_Temps"]


indice_T2_ete <- which(P3_ete_final[,"TEMPS_MESURE"] == "2")
P3_ete <- P3_ete_final[-indice_T2_ete,]

### Combined the 3 tables (for the graphics)
EVOL_ete <- rbind(P1_ete, P2_ete, P3_ete)

```

#### Figure 3-4-5 (panel a & d)

````
```{r}

#### spring ##
croissance_net_printemps2 <- croissance_net_printemps

### indice_0 <- which(croissance_net_printemps2[,"TEMPS_MESURE"] == 0)
### croissance_net_printemps2 <- croissance_net_printemps2[-indice_0,]

croissance_net_printemps2$PERIODE_FINAL <-
factor(croissance_net_printemps2$TEMPS_MESURE,
                                          levels = c("T1", "T2", "T3"),
                                          labels = c("P1", "P2", "P3"))

croissance_net_printemps2$ESPECE <-
factor(croissance_net_printemps2$ESPECE,
                                          levels = c("Cymodocea nodosa",
"Zostera noltei", "Ruppia cirrhosa"),
                                          labels = c("CN", "ZN", "RC"))

croissance_net_printemps_pivot <- croissance_net_printemps2 %>%
  pivot_longer(cols = c(Taux_de_croissance_par_pieds), names_to =
"Variable", values_to = "INDICE")

croissance_net_printemps_pivot$saison <- "spring"

colnames(croissance_net_printemps_pivot)[colnames(croissance_net_printemp
s_pivot) == "TEMPS_MESURE"] <- "PERIODE"

croissance_net_printemps_pivot$PERIODE <-
as.factor(croissance_net_printemps_pivot$PERIODE)

##Leaf loss rate
merge_PF_evol_spring_sansT3$PERIODE_FINAL <-
factor(merge_PF_evol_spring_sansT3$PERIODE,
                                          levels = c(0, 1, 2),
                                          labels = c("P1", "P2", "P3"))

PERTE_printemps_pivot <- merge_PF_evol_spring_sansT3 %>%
  pivot_longer(cols = c(indice_PF), names_to = "Variable", values_to =
"INDICE")

PERTE_printemps_pivot$saison <- "spring"

PERTE_printemps_pivot$PERIODE <- as.factor(PERTE_printemps_pivot$PERIODE)


Resume_tx_perte_spring <- merge_PF_evol_spring_sansT3 %>%
  group_by(ESPECE, EXPERIENCE, PERIODE_FINAL) %>%
  dplyr::summarise(
    mean_tx = mean(indice_PF),
    sd_tx = sd(indice_PF),
    n_obs = n(),
    se_tx = sd_tx / sqrt(n_obs)
  )

#combiner les deux tableaux avant de faire le graphique
````
````

```
GRAPH_Spring <- bind_rows(croissance_net_printemps_pivot,
PERTE_printemps_pivot)

## summer ##

croissance_net_ete <- croissance_net


croissance_net_ete$PERIODE_FINAL <-
factor(croissance_net_ete$TEMPS_MESURE,
                                          levels = c("T1", "T2", "T3"),
                                          labels = c("P1", "P2", "P3"))

croissance_net_ete$ESPECE <- factor(croissance_net_ete$ESPECE,
levels = c("Cymodocea nodosa", "Zostera noltei", "Ruppia cirrhosa"),
labels = c("CN", "ZN", "RC"))


croissance_net_ete_pivot <- croissance_net_ete %>%
  pivot_longer(cols = c(Taux_de_croissance_par_pieds), names_to =
"Variable", values_to = "INDICE")

croissance_net_ete_pivot$saison <- "summer"

colnames(croissance_net_ete_pivot)[colnames(croissance_net_ete_pivot) ==
"TEMPS_MESURE"] <- "PERIODE"

croissance_net_ete_pivot$PERIODE <-
as.factor(croissance_net_ete_pivot$PERIODE)

## Leaf loss rate

merge_PF_evol_summer_sansT3$PERIODE_FINAL <-
factor(merge_PF_evol_summer_sansT3$PERIODE,
                                          levels = c(0, 1, 2),
                                          labels = c("P1", "P2", "P3"))

PERTE_ete_pivot <- merge_PF_evol_summer_sansT3 %>%
  pivot_longer(cols = c(indice_PF), names_to = "Variable", values_to =
"INDICE")

PERTE_ete_pivot$saison <- "summer"

PERTE_ete_pivot$TEMPS_MESURE <- as.factor(PERTE_ete_pivot$TEMPS_MESURE)

Resume_tx_perte_summer <- merge_PF_evol_summer_sansT3 %>%
  group_by(ESPECE, EXPERIENCE, PERIODE_FINAL) %>%
  dplyr::summarise(
    mean_tx = mean(indice_PF),
    sd_tx = sd(indice_PF),
    n_obs = n(),
    se_tx = sd_tx / sqrt(n_obs)
  )

GRAPH_Summer<- bind_rows(croissance_net_ete_pivot, PERTE_ete_pivot)

#Combined for graphic
GRAPH1 <-bind_rows(GRAPH_Spring, GRAPH_Summer)
```

```
couleur_valeurs <- c("#9ecae1", "#f77f58", "#a1d99b")
etiquettes <- c("Before", "Heatwave","After")


GRAPH1_ZN <-
GRAPH1 %>%
  filter(ESPECE == "ZN") %>%
  mutate(Variable = factor(Variable, levels =
c("Taux_de_croissance_par_pieds", "indice_PF"))) %>%
  ggplot(aes(x = factor(EXPERIENCE, levels = c("C", "2", "4", "6")), y =
INDICE, fill = PERIODE_FINAL)) +
  geom_boxplot(show.legend = T)+
  geom_vline(xintercept = 1.5, linetype = "dashed", color = "gray", size
= 0.5) +
  geom_vline(xintercept = 2.5, linetype = "dashed", color = "gray", size
= 0.5) +
  geom_vline(xintercept = 3.5, linetype = "dashed", color = "gray", size
= 0.5) +
  stat_summary(fun = mean, geom = "point", color = "red", position =
position_dodge(width = 0.75), show.legend = T) +
  scale_fill_manual(values = couleur_valeurs, labels = etiquettes) +
  scale_x_discrete(breaks = c("C", "2", "4", "6"),
  labels = c("Control", "+2", "+4", "+6")) +
  theme_bw() +
   facet_grid(Variable~saison, scales = "free_y",
              labeller = labeller(saison = c("spring" = "Spring",
"summer" = "Summer"),
                                  Variable =
c("Taux_de_croissance_par_pieds" = "Leaf growth rate", "indice_PF" =
"Leaf loss rate"))) +
   labs(y = expression(mm^-2~d^-1),
        x = "Experimental conditions",
        fill = " ",
        title = " ") +
   theme(strip.text.x = element_text(size = 20),
         strip.text.y = element_text(size = 15),
         strip.background = element_rect(fill = "white"),
        axis.text.x = element_text(size = 18),
        axis.text.y = element_text(size = 18),
        axis.title.y = element_text(size = 20),
        axis.title.x = element_text(size = 20),
        legend.text = element_text(size = 18),
        legend.key.size = unit(1.5, "cm"))+
  geom_text(data = . %>% filter(Variable ==
"Taux_de_croissance_par_pieds", saison == "spring"),
            aes(x = 0.45, y = 60,
                label = "E : p = .02\nP : p < .001\nE:P : p = .07"),
            inherit.aes = FALSE, size = 5, color = "#4D4D4D", hjust = 0)
+
  geom_text(data = . %>% filter(Variable == "indice_PF", saison ==
"spring"),
            aes(x = 0.45, y = -200,
                label = "E : ns\nP : p < .001\nE:P : ns"),
            inherit.aes = FALSE, size = 5,color = "#4D4D4D", hjust = 0,
vjust = 0.1)  +
```

```
  geom_text(data = . %>% filter(Variable ==
"Taux_de_croissance_par_pieds", saison == "summer"),
            aes(x = 0.45, y = max(INDICE, na.rm = TRUE) * 0.9,
                label = "E : p < .001\nP : p < .001\nE:P : p = .07"),
            inherit.aes = FALSE, size = 5, color = "#4D4D4D", hjust = 0)
+
  geom_text(data = . %>% filter(Variable == "indice_PF", saison ==
"summer"),
            aes(x = 0.45, y = -200,
                label = "E : ns\nP : p < .001\nE:P : ns"),
            inherit.aes = FALSE, size = 5,color = "#4D4D4D", hjust = 0,
vjust = 0.1)




GRAPH1_RC <-
GRAPH1 %>%
  filter(ESPECE == "RC") %>%
  mutate(Variable = factor(Variable, levels =
c("Taux_de_croissance_par_pieds", "indice_PF"))) %>%
  ggplot(aes(x = factor(EXPERIENCE, levels = c("C", "2", "4", "6")), y =
INDICE, fill = PERIODE_FINAL)) +
  geom_boxplot(show.legend = T)+
  geom_vline(xintercept = 1.5, linetype = "dashed", color = "gray", size
= 0.5) +
  geom_vline(xintercept = 2.5, linetype = "dashed", color = "gray", size
= 0.5) +
  geom_vline(xintercept = 3.5, linetype = "dashed", color = "gray", size
= 0.5) +
  stat_summary(fun = mean, geom = "point", color = "red", position =
position_dodge(width = 0.75), show.legend = T) +
  scale_fill_manual(values = couleur_valeurs, labels = etiquettes) +
  scale_x_discrete(breaks = c("C", "2", "4", "6"),
  labels = c("Control", "+2", "+4", "+6")) +
  theme_bw() +
   facet_grid(Variable~saison, scales = "free_y",
              labeller = labeller(saison = c("spring" = "Spring",
"summer" = "Summer "),
                                  Variable =
c("Taux_de_croissance_par_pieds" = "Leaf growth rate ", "indice_PF" =
"Leaf loss rate"))) +
   labs(y = expression(mm^-2~d^-1),
        x = "Experimental conditions",
        fill = " ",
        title = " ") +
   theme(strip.text.x = element_text(size = 20),
         strip.text.y = element_text(size = 15),
         strip.background = element_rect(fill = "white"),
        axis.text.x = element_text(size = 18),
        axis.text.y = element_text(size = 18),
        axis.title.y = element_text(size = 20),
        axis.title.x = element_text(size = 20),
        legend.text = element_text(size = 18),
        legend.key.size = unit(1.5, "cm"))+

  geom_text(data = . %>% filter(Variable ==
"Taux_de_croissance_par_pieds", saison == "spring"),
```

```
            aes(x = 0.45, y = 17,
                label = "E : p < .03\nP : ns\nE:P : ns"),
            inherit.aes = FALSE, size = 5, color = "#4D4D4D", hjust = 0)
+
  geom_text(data = . %>% filter(Variable == "indice_PF", saison ==
"spring"),
            aes(x = 0.45, y = -23,
                label = "E : ns\nP : ns\nE:P : ns"),
            inherit.aes = FALSE, size = 5,color = "#4D4D4D", hjust = 0,
vjust = 0.4)+

  geom_text(data = . %>% filter(Variable ==
"Taux_de_croissance_par_pieds", saison == "summer"),
            aes(x = 0.45, y = 17,
                label = "E: p < .001\nP : p < .001\nE:P : p < .001"),
            inherit.aes = FALSE, size = 5, color = "#4D4D4D", hjust = 0)
+
  geom_text(data = . %>% filter(Variable == "indice_PF", saison ==
"summer"),
            aes(x = 0.45, y = -23,
                label = "E : ns\nP : ns\nE:P : ns"),
            inherit.aes = FALSE, size = 5,color = "#4D4D4D", hjust = 0,
vjust = 0.4)



GRAPH1_CN <-
GRAPH1 %>%
  filter(ESPECE == "CN") %>%
  mutate(Variable = factor(Variable, levels =
c("Taux_de_croissance_par_pieds", "indice_PF"))) %>%
  ggplot(aes(x = factor(EXPERIENCE, levels = c("C", "2", "4", "6")), y =
INDICE, fill = PERIODE_FINAL)) +
  geom_boxplot(show.legend = T)+
  geom_vline(xintercept = 1.5, linetype = "dashed", color = "gray", size
= 0.5) +
  geom_vline(xintercept = 2.5, linetype = "dashed", color = "gray", size
= 0.5) +
  geom_vline(xintercept = 3.5, linetype = "dashed", color = "gray", size
= 0.5) +
  stat_summary(fun = mean, geom = "point", color = "red", position =
position_dodge(width = 0.75), show.legend = T) +
  scale_fill_manual(values = couleur_valeurs, labels = etiquettes) +
  scale_x_discrete(breaks = c("C", "2", "4", "6"),
  labels = c("Control", "+2", "+4", "+6")) +
  theme_bw() +
   facet_grid(Variable~saison, scales = "free_y",
              labeller = labeller(saison = c("spring" = "Spring",
"summer" = "Summer"),
                                  Variable =
c("Taux_de_croissance_par_pieds" = "Leaf growth rate", "indice_PF" =
"Leaf loss rate"))) +
   labs(y = expression(mm^-2~d^-1),
        x = "Experimental conditions",
        fill = " ",
        title = " ") +
   theme(strip.text.x = element_text(size = 20),
         strip.text.y = element_text(size = 15),
```

````
          strip.background = element_rect(fill = "white"),
         axis.text.x = element_text(size = 18),
         axis.text.y = element_text(size = 18),
         axis.title.y = element_text(size = 20),
         axis.title.x = element_text(size = 20),
         legend.text = element_text(size = 18),
         legend.key.size = unit(1.5, "cm"))+
  geom_text(data = . %>% filter(Variable ==
"Taux_de_croissance_par_pieds", saison == "spring"),
            aes(x = 0.45, y = 68,
                label = "E : p < .001\nP : p < .001\nE:P : p < .001"),
            inherit.aes = FALSE, size = 5, color = "#4D4D4D", hjust = 0)
+
  geom_text(data = . %>% filter(Variable == "indice_PF", saison ==
"spring"),
            aes(x = 0.45, y = -245,
                label = "E : p < .001\nP : p < .001\nE:P : p < .001"),
            inherit.aes = FALSE, size = 5,color = "#4D4D4D", hjust = 0,
vjust = 0.4)+
  geom_text(data = . %>% filter(Variable ==
"Taux_de_croissance_par_pieds", saison == "summer"),
            aes(x = 0.45, y = 68,
                label = "E : ns\nP : p < .001\nE:P : ns"),
            inherit.aes = FALSE, size = 5, color = "#4D4D4D", hjust = 0)
+
  geom_text(data = . %>% filter(Variable == "indice_PF", saison ==
"summer"),
            aes(x = 0.45, y = -245,
                label = "E : ns\nP : p < .001\nE:P : p < .01"),
            inherit.aes = FALSE, size = 5,color = "#4D4D4D", hjust = 0,
vjust = 0.4)

```

#### Figure 3-4-5 (panel b & e)

```{r}


### Spring ###
NF_spring <- merge_NF_evol_spring %>%
  mutate(PERIODE = case_when(
    PERIODE == 1 ~ "P1",
    PERIODE == 2 ~ "P2",
    PERIODE == 3 ~ "P3",
    TRUE ~ as.factor(PERIODE)
  ))

OLD_spring <- EVOL_SPRING %>%
  mutate(TEMPS_MESURE = case_when(
    TEMPS_MESURE == 1 ~ "P1",
    TEMPS_MESURE == 2 ~ "P2",
    TEMPS_MESURE == 3 ~ "P3",
    TRUE ~ as.factor(TEMPS_MESURE)
  ))

OLD_spring$PERIODE <- OLD_spring$TEMPS_MESURE
````

```
donnees_combinees_spring <- NF_spring %>%
  inner_join(OLD_spring,
             by = c("ind_pieds", "PERIODE"),
             suffix = c("_NF", "_OLD"))


moyennes_spring <- donnees_combinees_spring %>%
  group_by(PERIODE, EXPERIENCE_NF, ESPECE_NF) %>%
  summarise(
    mean_old = mean(evolution_jour, na.rm = TRUE),
    mean_New = mean(indice_NF, na.rm = TRUE),
    .groups = 'drop'
  )



### Summer ###

NF_summer <- merge_NF_evol_summer %>%
  mutate(PERIODE = case_when(
    PERIODE == 1 ~ "P1",
    PERIODE == 2 ~ "P2",
    PERIODE == 3 ~ "P3",
    TRUE ~ as.factor(PERIODE)
  ))

OLD_summer <- EVOL_ete %>%
  mutate(TEMPS_MESURE = case_when(
    TEMPS_MESURE == 1 ~ "P1",
    TEMPS_MESURE == 2 ~ "P2",
    TEMPS_MESURE == 3 ~ "P3",
    TRUE ~ as.factor(TEMPS_MESURE)
  ))

OLD_summer$PERIODE <- OLD_summer$TEMPS_MESURE


donnees_combinees_summer <- NF_summer %>%
  inner_join(OLD_summer,
             by = c("ind_pieds", "PERIODE"),
             suffix = c("_NF", "_OLD"))


moyennes_summer <- donnees_combinees_summer %>%
  group_by(PERIODE, EXPERIENCE_NF, ESPECE_NF) %>%
  summarise(
    mean_old = mean(evolution_jour, na.rm = TRUE),
    mean_New = mean(indice_NF, na.rm = TRUE),
    .groups = 'drop'
  )

### Graphiques ###

# spring
donnees_combinees_spring$saison <- "spring"
moyennes_spring$saison <- "spring"

# summer
```

```
donnees_combinees_summer$saison <- "summer"
moyennes_summer$saison <- "summer"


OLD_NEW2SAISON <- rbind(donnees_combinees_spring,
donnees_combinees_summer)
OLD_NEW2SAISON_moyennes <- rbind(moyennes_spring, moyennes_summer)


# Graphique pour l'espèce CN
OLD_NEW2SAISON_CN <- OLD_NEW2SAISON %>%
  filter(ESPECE_OLD == "CN")

OLD_NEW2SAISON_moyennes_CN <- OLD_NEW2SAISON_moyennes %>%
  filter(ESPECE_NF == "CN")

# Figure : CN

OLD_NEW_saison_CN <-
ggplot() +
  geom_point(data = OLD_NEW2SAISON_CN, aes(x = evolution_jour, y =
indice_NF, fill = as.factor(PERIODE), color = as.factor(PERIODE),
                                                   shape =
factor(EXPERIENCE_NF, levels = c("C", "2", "4", "6"))), size = 2.5, alpha
= 0.7) +
  geom_point(data = OLD_NEW2SAISON_moyennes_CN, aes(x = mean_old, y =
mean_New, shape = factor(EXPERIENCE_NF, levels = c("C", "2", "4", "6")),
                                         fill = as.factor(PERIODE)),
color = "black", size = 4, stroke = 1.5,show.legend = F) +  # Contour
noirfixe
  geom_text_repel(data = OLD_NEW2SAISON_moyennes_CN, aes(x = mean_old, y
= mean_New, label = factor(EXPERIENCE_NF, levels = c("C", "2", "4",
"6"))),
                  size = 4, color = "black", min.segment.length = 0, seed
= 10, box.padding = 0.5) +
  geom_abline(slope = 1, inteCNept = 0, linetype = "dashed", color =
"black") +
  facet_grid(~saison, scales = "free_y")+
  scale_shape_manual(values = c(21, 22, 24, 23),
                     labels = c("Control", "+2°C", "+4°C", "+6°C")) +
  scale_fill_manual(values = c("#9ecae1", "#f77f58", "#a1d99b"),
                    labels = c("Before", "Heatwave", "After"),
                    guide = "none") +
  scale_color_manual(values = c("#9ecae1", "#f77f58", "#a1d99b"),
                     labels = c("Before", "Heatwave", "After")) +
labs(
    x = expression("Old leaf gain rate"~(mm^-2~d^-1)),
       y = expression("New leaf gain rate"~(mm^-2~d^-1)),
       fill = " ",
       color = " ",
       shape = " ") +
  theme_bw() +
  xlim(0,38)+
  theme(
    strip.text.x = element_blank(),
    plot.title = element_text(hjust = 0, size = 16),
    plot.subtitle = element_text(hjust = 0.5, size = 12, color =
"gray40"),
```

```
    legend.position = "right",
    legend.box = "horizontal",
    legend.text = element_text(size = 18),
    strip.text = element_text(size = 12, face = "bold"),
    axis.text.x = element_text(size = 18),
    axis.text.y = element_text(size = 18),
    axis.title.y = element_text(size = 20),
    axis.title.x = element_text(size = 20)
  )


# Figure : ZN
OLD_NEW2SAISON_ZN <- OLD_NEW2SAISON %>%
  filter(ESPECE_OLD == "ZN")

OLD_NEW2SAISON_moyennes_ZN <- OLD_NEW2SAISON_moyennes %>%
  filter(ESPECE_NF == "ZN")


OLD_NEW_saison_ZN <-
ggplot() +
  geom_point(data = OLD_NEW2SAISON_ZN, aes(x = evolution_jour, y =
indice_NF, fill = as.factor(PERIODE), color = as.factor(PERIODE),
                                                  shape =
factor(EXPERIENCE_NF, levels = c("C", "2", "4", "6"))), size = 2.5, alpha
= 0.6) +
  geom_point(data = OLD_NEW2SAISON_moyennes_ZN, aes(x = mean_old, y =
mean_New, shape = factor(EXPERIENCE_NF, levels = c("C", "2", "4", "6")),
                                         fill = as.factor(PERIODE)),
color = "black", size = 4, stroke = 1.5,show.legend = F) +  # Contour
noirfixe
  geom_text_repel(data = OLD_NEW2SAISON_moyennes_ZN, aes(x = mean_old, y
= mean_New, label = factor(EXPERIENCE_NF, levels = c("C", "2", "4",
"6"))),
                  size = 4, color = "black", min.segment.length = 0, seed
= 10, box.padding = 0.5) +
  geom_abline(slope = 1, inteZNept = 0, linetype = "dashed", color =
"black") +
  facet_grid(~saison, scales = "free")+
  scale_shape_manual(values = c(21, 22, 24, 23),
                     labels = c("Control", "+2°C", "+4°C", "+6°C")) +
  scale_fill_manual(values = c("#9ecae1", "#f77f58", "#a1d99b"),
                    labels = c("Before", "Heatwave", "After"),
                    guide = "none") +
  scale_color_manual(values = c("#9ecae1", "#f77f58", "#a1d99b"),
                     labels = c("Before", "Heatwave", "After")) +
labs(
    x = expression("Old leaf gain rate"~(mm^-2~d^-1)),
       y = expression("New leaf gain rate"~(mm^-2~d^-1)),
       fill = " ",
       color = " ",
       shape = " ") +
  theme_bw() +
  xlim(0,35)+
  theme(
    strip.text.x = element_blank(),
    plot.title = element_text(hjust = 0, size = 16),
```

```
    plot.subtitle = element_text(hjust = 0.5, size = 12, color =
"gray40"),
    legend.position = "right",
    legend.box = "horizontal",
    legend.text = element_text(size = 18),
    strip.text = element_text(size = 12, face = "bold"),
    axis.text.x = element_text(size = 18),
    axis.text.y = element_text(size = 18),
    axis.title.y = element_text(size = 20),
    axis.title.x = element_text(size = 20)
  )


# Figure : RC
OLD_NEW2SAISON_RC <- OLD_NEW2SAISON %>%
  filter(ESPECE_OLD == "RC")

OLD_NEW2SAISON_moyennes_RC <- OLD_NEW2SAISON_moyennes %>%
  filter(ESPECE_NF == "RC")


OLD_NEW_saison_RC <-
ggplot() +
  geom_point(data = OLD_NEW2SAISON_RC, aes(x = evolution_jour, y =
indice_NF, fill = as.factor(PERIODE), color = as.factor(PERIODE),
                                                 shape =
factor(EXPERIENCE_NF, levels = c("C", "2", "4", "6"))), size = 2.5, alpha
= 0.6) +
  geom_point(data = OLD_NEW2SAISON_moyennes_RC, aes(x = mean_old, y =
mean_New, shape = factor(EXPERIENCE_NF, levels = c("C", "2", "4", "6")),
                                         fill = as.factor(PERIODE)),
color = "black", size = 4, stroke = 1.5,show.legend = F) +  # Contour
noirfixe
  geom_text_repel(data = OLD_NEW2SAISON_moyennes_RC, aes(x = mean_old, y
= mean_New, label = factor(EXPERIENCE_NF, levels = c("C", "2", "4",
"6"))),
                  size = 4, color = "black", min.segment.length = 0, seed
= 10, box.padding = 0.5) +
  geom_abline(slope = 1, intercept = 0, linetype = "dashed", color =
"black") +
  facet_grid(~saison, scales = "free")+
  scale_shape_manual(values = c(21, 22, 24, 23),
                     labels = c("Control", "+2°C", "+4°C", "+6°C")) +
  scale_fill_manual(values = c("#9ecae1", "#f77f58", "#a1d99b"),
                    labels = c("Before", "Heatwave", "After"),
                    guide = "none") +
  scale_color_manual(values = c("#9ecae1", "#f77f58", "#a1d99b"),
                     labels = c("Before", "Heatwave", "After")) +
labs(
    x = expression("Old leaf gain rate"~(mm^-2~d^-1)),
       y = expression("New leaf gain rate"~(mm^-2~d^-1)),
       fill = " ",
       color = " ",
       shape = " ") +
  theme_bw() +
  xlim(0, 10)+
  theme(
    strip.text.x = element_blank(),
```

```
    plot.title = element_text(hjust = 0, size = 16),
    plot.subtitle = element_text(hjust = 0.5, size = 12, color =
"gray40"),
    legend.position = "right",
    legend.box = "horizontal",
    legend.text = element_text(size = 18),
    strip.text = element_text(size = 12, face = "bold"),
    axis.text.x = element_text(size = 18),
    axis.text.y = element_text(size = 18),
    axis.title.y = element_text(size = 20),
    axis.title.x = element_text(size = 20)
  )

# Calcul des ratios


ratios_par_traitement_spring <- donnees_combinees_spring %>%
  # Grouper par traitement et période
  group_by(EXPERIENCE_NF, PERIODE,ESPECE_NF) %>%
  # Calculer les moyennes puis le ratio
  summarise(
    moy_new = mean(indice_NF, na.rm = TRUE),
    moy_old = mean(evolution_jour, na.rm = TRUE),
    ratio_spring = moy_new/moy_old,
    sd_new = sd(indice_NF, na.rm = TRUE),
    sd_old = sd(evolution_jour, na.rm = TRUE),
    ratio_spring_sd = ratio_spring * sqrt((sd_new / moy_new)^2 + (sd_old
/ moy_old)^2)
    ) %>%
  ungroup()


ratios_par_traitement_summer <- donnees_combinees_summer %>%
  # Grouper par traitement et période
  group_by(EXPERIENCE_NF, PERIODE,ESPECE_NF) %>%
  # Calculer les moyennes puis le ratio
  summarise(
    moy_new_summer = mean(indice_NF, na.rm = TRUE),
    moy_old = mean(evolution_jour, na.rm = TRUE),
    ratio_summer = moy_new/moy_old,
    sd_new = sd(indice_NF, na.rm = TRUE),
    sd_old = sd(evolution_jour, na.rm = TRUE),
    ratio_summer_sd = ratio_summer * sqrt((sd_new / moy_new)^2 + (sd_old
/ moy_old)^2)
  ) %>%
  ungroup()


```

#### Statistical analysis LGR (lmerTest) - Spring and Summer

```{r}
## stat test growth rate
# Function to analyze data by species and season
analyze_species2 <- function(data, species, season) {
  cat("##", species, "----/n")
```

```
  # Filter the data by species and season
  data_filtered <- data %>% filter(ESPECE == species)
  # data_filtered$TEMPS_MESURE <- as.factor(data_filtered$TEMPS_MESURE)
  data_filtered$EXPERIENCE <- factor( data_filtered$EXPERIENCE, levels =
c("C", "2","4", "6"))

  # Test the full model and the null model
  test_model <- lmer(Taux_de_croissance_par_pieds ~ EXPERIENCE *
TEMPS_MESURE + (1 | REPLICAT),
                     data = data_filtered, REML = FALSE)
  test_null <- lmer(Taux_de_croissance_par_pieds ~ 1 + (1 | REPLICAT),
                    data = data_filtered, REML = FALSE)
  summary(test_model)
  anova(test_model, test_null)

  # Pairwise comparisons for untransformed data
 test_model_REML <- lmer(Taux_de_croissance_par_pieds ~ EXPERIENCE *
TEMPS_MESURE + (1 | REPLICAT),
                        data = data_filtered, REML = TRUE)
  summary(test_model_REML)
  anova_results <- anova(test_model_REML)

  # Convert the ANOVA results to a data frame for export
anova_df <- as.data.frame(anova_results)

# Save the results with write.csv2
write.csv2(anova_df, file = paste0("anova_results_", species, "_",
season, ".csv"), row.names = TRUE)

  comp <- emmeans(test_model_REML, ~ EXPERIENCE * TEMPS_MESURE)
  pairwise_SOUSP <- pairs(comp, simple = "TEMPS_MESURE")
  pairwise_EXPERIENCE <- pairs(comp, simple = "EXPERIENCE")

    # Extract the results in a structured table (e.g., estimates,
standard errors, p-values)
pairwise_SOUSP_df <- as.data.frame(summary(pairwise_SOUSP)[,
c("contrast","estimate", "SE", "df", "t.ratio", "p.value")])
pairwise_EXPERIENCE_df <- as.data.frame(summary(pairwise_EXPERIENCE)[,
c("contrast","estimate", "SE", "df", "t.ratio", "p.value")])

# Save the results in a CSV file using write.csv2 (separator ';')
write.csv2(pairwise_SOUSP_df, file = paste0("tx_brut_", species,
"_SOUSP_", season, ".csv"), row.names = FALSE)
write.csv2(pairwise_EXPERIENCE_df, file = paste0("tx_brut_", species,
"_EXPERIENCE_", season, ".csv"), row.names = FALSE)

  residuals <- resid(test_model, type = "pearson")
  png(file.path(getwd(), paste0(species, "_", season, "_tx_brut.png")))
  op <- par(mfrow = c(2, 2))
  plot(x = fitted(test_model), y = residuals)
  abline(0, 0, col = "red")
  hist(residuals)
  qqnorm(residuals)
  boxplot(residuals ~ EXPERIENCE:TEMPS_MESURE, data = data_filtered,
na.action = na.omit, varwidth = TRUE)
  dev.off()
}
```

````
# Analyze for each species and season
analyze_species2(croissance_net, "Cymodocea nodosa", "summer")
analyze_species2(croissance_net_printemps, "Cymodocea nodosa", "spring")
analyze_species2(croissance_net, "Zostera noltei", "summer")
analyze_species2(croissance_net_printemps, "Zostera noltei", "spring")
analyze_species2(croissance_net, "Ruppia cirrhosa", "summer")
analyze_species2(croissance_net_printemps, "Ruppia cirrhosa", "spring")

```

#### Statistical analysis LLR (lmerTest) - Spring and Summer

```{r}
library(lmerTest)
## stat test growth rate
# Function to analyze data by species and season
analyze_species <- function(data, species, season) {
  cat("##", species, "----/n")


  # Filter the data by species and season
  data_filtered <- data %>% filter(ESPECE == species)
  data_filtered$EXPERIENCE <- factor( data_filtered$EXPERIENCE, levels =
c("C", "2","4", "6"))

  # Test the full model and the null model
  test_model <- lmer(indice_PF ~ EXPERIENCE * PERIODE_FINAL + (1 |
REPLICAT),
                     data = data_filtered, REML = FALSE)
  test_null <- lmer(indice_PF ~ 1 + (1 | REPLICAT),
                    data = data_filtered, REML = FALSE)
  summary(test_model)
  anova(test_model, test_null)

  # Pairwise comparisons for untransformed data
 test_model_REML <- lmer(indice_PF ~ EXPERIENCE * PERIODE_FINAL + (1 |
REPLICAT),
                        data = data_filtered, REML = TRUE)
  summary(test_model_REML)
  anova_results <- anova(test_model_REML)

  # Convert the ANOVA results to a data frame for export
anova_df <- as.data.frame(anova_results)

# Save the results with write.csv2
write.csv2(anova_df, file = paste0("anova_results_LOSS_", species, "_",
season, ".csv"), row.names = TRUE)

  comp <- emmeans(test_model_REML, ~ EXPERIENCE * PERIODE_FINAL)
  pairwise_SOUSP <- pairs(comp, simple = "PERIODE_FINAL")
  pairwise_EXPERIENCE <- pairs(comp, simple = "EXPERIENCE")

    # Extract the results in a structured table (e.g., estimates,
standard errors, p-values)
pairwise_SOUSP_df <- as.data.frame(summary(pairwise_SOUSP)[,
c("contrast","estimate", "SE", "df", "t.ratio", "p.value")])
````

```
pairwise_EXPERIENCE_df <- as.data.frame(summary(pairwise_EXPERIENCE)[,
c("contrast","estimate", "SE", "df", "t.ratio", "p.value")])

# Save the results in a CSV file using write.csv2 (separator ';')
write.csv2(pairwise_SOUSP_df, file = paste0("LOSS_", species, "_SOUSP_",
season, ".csv"), row.names = FALSE)
write.csv2(pairwise_EXPERIENCE_df, file = paste0("LOSS_", species,
"_EXPERIENCE_", season, ".csv"), row.names = FALSE)

  residuals <- resid(test_model, type = "pearson")
  png(file.path(getwd(), paste0(species, "_", season, "_LOSS.png")))
  op <- par(mfrow = c(2, 2))
  plot(x = fitted(test_model), y = residuals)
  abline(0, 0, col = "red")
  hist(residuals)
  qqnorm(residuals)
  boxplot(residuals ~ EXPERIENCE:PERIODE_FINAL, data = data_filtered,
na.action = na.omit, varwidth = TRUE)
  dev.off()
}

# Analyze for each species and season
analyze_species(merge_PF_evol_summer_sansT3, "CN", "summer")
analyze_species(merge_PF_evol_spring_sansT3, "CN", "spring")
analyze_species(merge_PF_evol_summer_sansT3, "ZN", "summer")
analyze_species(merge_PF_evol_spring_sansT3, "ZN", "spring")
analyze_species(merge_PF_evol_summer_sansT3, "RC", "summer")
analyze_species(merge_PF_evol_spring_sansT3, "RC", "spring")

```

#### Oxygen data (GCP and CR)

#### SUMMER

```{r}

#########################################################################
########
########################## Leaf area data SUMMER
################################
#########################################################################
########

fichier <- "biometrie_ete.csv"

# Set the working directory to the location of the data
# Donnees_biometrique <- "~" # add your working directory here
setwd(Donnees_biometrique)

# Load the data
biometrie_ete <- read.table(fichier, sep = ",", h = TRUE, dec = ",")


# Convert the measurement date to Date format
biometrie_ete$DATE_MESURE <- dmy(biometrie_ete$DATE_MESURE)

# Create unique identifiers for each leaf and replicate
```

```
biometrie_ete <- biometrie_ete %>% mutate(ind = paste0(ESPECE, REPLICAT,
EXPERIENCE, INDIVIDUS, NUMERO_FEUILLE))  ## combiner les colonnes en une
seule
biometrie_ete <- biometrie_ete %>% mutate(ind_pieds = paste0(ESPECE,
REPLICAT, EXPERIENCE, INDIVIDUS))  ## combiner les colonnes en une seule
biometrie_ete <- biometrie_ete %>% mutate(ID_POTS =
paste0(ESPECE,REPLICAT))

# Remove rows where the "MORT" column is not empty
indice_mort <- which(biometrie_ete[,"MORT"] != "")
biometrie_ete <- biometrie_ete[-indice_mort,]

#### Calculate leaf area in mm² ####
biometrie_ete$LARGEUR_FEUILLE_mm <-
as.numeric(biometrie_ete$LARGEUR_FEUILLE_mm)
biometrie_ete[,"SURFACE_FOLIAIRE_MM2"] <-
(biometrie_ete[,"LONGUEUR_FEUILLE_mm"])*(biometrie_ete[,"LARGEUR_FEUILLE_
mm"])
biometrie_ete[,"SURFACE_FOLIAIRE_CM2"] <-
((biometrie_ete[,"LONGUEUR_FEUILLE_mm"])*(biometrie_ete[,"LARGEUR_FEUILLE
_mm"])) / 100

# Now calculate leaf area by replicate (pot)
df_pots <- biometrie_ete %>%
  group_by(ESPECE, EXPERIENCE, REPLICAT, TEMPS_MESURE) %>%
  summarise(SF_POT_CM2 = sum(SURFACE_FOLIAIRE_MM2),
            n_pieds = n_distinct(INDIVIDUS)) %>%
  ungroup()

df_pots <- df_pots %>% mutate(ID_POTS = paste0(ESPECE,REPLICAT))

df_pots$EXPERIENCE <- as.factor(df_pots$EXPERIENCE)

########################################################################
########
######################## Oxygen data SUMMER
###################################
########################################################################
########
fichier <- "incubation_ete.csv"


# Set the working directory for the incubation data
# Donnees_incub <- "~" # add your working directory here


setwd(Donnees_incub)

# Load the incubation data
incubation_ete <- read.table(fichier, sep = "\t", h = TRUE, dec = ",")

# Convert EXP column to factor and redefine the levels
incubation_ete$EXP <- as.factor(incubation_ete$EXP)
levels(incubation_ete$EXP) <- c("2", "4", "6", "C")

# Combine DATE_MESURE and HEURE_MESURE into a single column for time
  incubation_ete[,"DATE_MESURE"] <- dmy(incubation_ete[,"DATE_MESURE"])
```

```
  incubation_ete[,"Temps"] <- paste(incubation_ete[,"DATE_MESURE"],
incubation_ete[,"HEURE_MESURE"])
  incubation_ete[,"Temps"] <-
as.POSIXct(strptime(incubation_ete[,"Temps"], format="%Y-%m-%d
%H:%M:%S"), tz="UTC")

# Reshape the data: Pivot to get OXY_mg.L and Temps columns for different
conditions
  incubation_ete_cast_oxy <- pivot_wider(incubation_ete,
                                     id_cols = c(ID_POTS,
EXP,SOUS_PERIODE, ESPECE, NB_PIEDS, DATE_MESURE),
                                     names_from = CONDITIONS,
                                     values_from = OXY_mg.L)


  incubation_ete_cast_heure <- pivot_wider(incubation_ete,
                                       id_cols = c(ID_POTS, EXP,
SOUS_PERIODE, ESPECE, NB_PIEDS, DATE_MESURE),
                                       names_from = CONDITIONS,
                                       values_from = Temps)


# Calculate the time differences between "JOUR" and "NUIT" conditions
  incubation_ete_cast_heure[,"diff_temps_jour"] <-
(incubation_ete_cast_heure[,"JOUR 2"] - incubation_ete_cast_heure[,"JOUR
1"]) / 60
  incubation_ete_cast_heure[,"diff_temps_nuit"] <-
(incubation_ete_cast_heure[,"NUIT 2"] - incubation_ete_cast_heure[,"NUIT
1"]) / 60

# Calculate the oxygen differences between "JOUR" and "NUIT" conditions
incubation_ete_cast_oxy[,"diff_oxy_jour"] <-
(incubation_ete_cast_oxy[,"JOUR 2"] - incubation_ete_cast_oxy[,"JOUR 1"])
incubation_ete_cast_oxy[,"diff_oxy_nuit"] <-
(incubation_ete_cast_oxy[,"NUIT 2"] - incubation_ete_cast_oxy[,"NUIT 1"])

# Merge the data based on identifiers
identifiant <- c("ID_POTS", "EXP", "DATE_MESURE", "SOUS_PERIODE",
"NB_PIEDS", "ESPECE")
incubation_ete_final <- merge(incubation_ete_cast_heure[, c(identifiant,
"diff_temps_jour", "diff_temps_nuit")],
                            incubation_ete_cast_oxy[, c( c("ID_POTS",
"EXP", "DATE_MESURE", "SOUS_PERIODE", "NB_PIEDS", "ESPECE"),
"diff_oxy_jour", "diff_oxy_nuit")])

# Convert the time difference and oxygen difference columns to numeric
incubation_ete_final$diff_temps_jour <-
as.numeric(incubation_ete_final$diff_temps_jour)
incubation_ete_final$diff_oxy_jour <-
as.numeric(incubation_ete_final$diff_oxy_jour)
incubation_ete_final$diff_temps_nuit <-
as.numeric(incubation_ete_final$diff_temps_nuit)
incubation_ete_final$diff_oxy_nuit <-
as.numeric(incubation_ete_final$diff_oxy_nuit)
 # Calculate net production and respiration rates in mg O2 / cm2 / hour
incubation_ete_final[,"Production_net"] <-
(incubation_ete_final[,"diff_oxy_jour"] /
```

```
incubation_ete_final[,"diff_temps_jour"]) * 2  # Multiply by pot volume
(2 liters)
incubation_ete_final[,"Respiration"] <-
(incubation_ete_final[,"diff_oxy_nuit"] /
incubation_ete_final[,"diff_temps_nuit"]) * 2

# Calculate gross production by summing absolute values of net production
and respiration
incubation_ete_final[,"Production_brute"] <-
abs(incubation_ete_final[,"Production_net"]) +
abs(incubation_ete_final[,"Respiration"])

# Correct respiration values: Set positive values to zero
incubation_ete_final[,"Respiration_corr"] <-
incubation_ete_final[,"Respiration"]
indice_positive <- which(incubation_ete_final[,"Respiration_corr"] > 0)
incubation_ete_final[indice_positive,"Respiration_corr"] <- 0

# Calculate corrected gross production
incubation_ete_final[,"Production_brute_corr"] <-
abs(incubation_ete_final[,"Production_net"]) +
abs(incubation_ete_final[,"Respiration_corr"])

# Merge the leaf area data with the incubation data
colnames(incubation_ete_final)[which(colnames(incubation_ete_final) ==
"EXP")] <- "EXPERIENCE"
colnames(incubation_ete_final)[which(colnames(incubation_ete_final) ==
"SOUS_PERIODE")] <- "TEMPS_MESURE"
merge_data <- merge(incubation_ete_final, df_pots, by = c("ESPECE",
"TEMPS_MESURE", "EXPERIENCE", "ID_POTS"))

# Select relevant columns for further analysis
work_data <- merge_data[c("ESPECE", "TEMPS_MESURE", "EXPERIENCE",
"ID_POTS", "DATE_MESURE", "NB_PIEDS",
                          "Production_net", "Respiration_corr",
"Production_brute_corr", "SF_POT_CM2", "n_pieds")]

# Calculate mean leaf area per individual
resultats_moyens <- work_data %>%
  group_by(ESPECE, TEMPS_MESURE, EXPERIENCE, ID_POTS) %>%
  summarize(Moyenne_SF_pieds = mean(SF_POT_CM2 / n_pieds, na.rm = TRUE))

# Merge the mean leaf area with the original data
work_data_bis <- merge(work_data, resultats_moyens, by = c("ESPECE",
"TEMPS_MESURE", "EXPERIENCE", "ID_POTS"))

# Calculate the difference between total number of plants and distinct
number of plants
work_data_bis$Difference_Pieds <- work_data_bis$NB_PIEDS -
work_data_bis$n_pieds

# Adjust the leaf area based on the number of plants
work_data_bis <- work_data_bis %>%
  mutate(SF_POT_CM2_Corrigee = SF_POT_CM2 + Difference_Pieds *
Moyenne_SF_pieds)

# Correct the net production, respiration, and gross production based on
corrected leaf area
```

```
work_data_bis[,"Production_net_area"] <- work_data_bis[,"Production_net"]
/ work_data_bis[,"SF_POT_CM2_Corrigee"]
work_data_bis[,"Respiration_area"] <- work_data_bis[,"Respiration_corr"]
/ work_data_bis[,"SF_POT_CM2_Corrigee"]
work_data_bis[,"Production_brute_area"] <-
work_data_bis[,"Production_brute_corr"] /
work_data_bis[,"SF_POT_CM2_Corrigee"]

```

#### SPRING

```{r}
##########################################################################
########
########################## Leaf area data SPRING
################################
##########################################################################
########

  fichier <- "biometrie_printemps.csv"

# setwd to the data directory
# Donnees_biometrique <- "~" # add your working directory here
setwd(Donnees_biometrique)

# Load the data
biometrie <- read.table(fichier, sep = ",", h = TRUE, dec = ",")


# Calculate SF per pot:
biometrie$DATE_MESURE <- dmy(biometrie$DATE_MESURE)

biometrie <- biometrie %>% mutate(ind = paste0(ESPECE, REPLICAT,
EXPERIENCE, INDIVIDUS, NUMERO_FEUILLE))  ## combine columns into one
biometrie <- biometrie %>% mutate(ind_pieds = paste0(ESPECE, REPLICAT,
EXPERIENCE, INDIVIDUS))  ## combine columns into one
biometrie <- biometrie %>% mutate(ID_POTS = paste0(ESPECE,REPLICAT))

#### Calculate leaf area in mm² ####
biometrie$LARGEUR_FEUILLE_mm <- as.numeric(biometrie$LARGEUR_FEUILLE_mm)
biometrie[,"SURFACE_FOLIAIRE_MM2"] <-
(biometrie[,"LONGUEUR_FEUILLE_mm"])*(biometrie[,"LARGEUR_FEUILLE_mm"])
biometrie[,"SURFACE_FOLIAIRE_CM2"] <-
((biometrie[,"LONGUEUR_FEUILLE_mm"])*(biometrie[,"LARGEUR_FEUILLE_mm"]))
/ 100

# Now I want the leaf area per REPLICAT (pot)
df_pots <- biometrie %>%
  group_by(ESPECE, EXPERIENCE, REPLICAT, TEMPS_MESURE) %>%
  summarise(SF_POT_CM2 = sum(SURFACE_FOLIAIRE_MM2),
            n_pieds = n_distinct(INDIVIDUS)) %>%
  ungroup()

df_pots <- df_pots %>% mutate(ID_POTS = paste0(ESPECE,REPLICAT))

# Now I need to change the way EXP is written:
df_pots$EXPERIENCE <- as.factor(df_pots$EXPERIENCE)
```

```
# Remove the ID_pots for T0 in the spring since I don't have the initial
measurement
indice_T0 <- which(df_pots[,"TEMPS_MESURE"] == "T0")
df_pots <- df_pots[-indice_T0,]

########################################################################
########
########################## Oxygen data SPRING
###################################
########################################################################
########


# Now I need the real number of feet, so I will use the incubation table,
the ID_POTS column, and nb_pieds
fichier <- "incubation_printemps.csv"

# setwd to the data directory
# Donnees_incub <- "~" # add your working directory here

setwd(Donnees_incub)

# Load the data
incubation <- read.table(fichier, sep = "\t", h = TRUE, dec = ",")

# Now I need to change the way EXP is written:
incubation$EXP <- as.factor(incubation$EXP)
levels(incubation$EXP) <- c("2", "4", "6", "C")

# I will process the incubation data to get production per pot
# Convert the "DATE_MESURE" and "HEURE_MESURE" columns into a single
"Temps" column in POSIXct format (date + time)
incubation[,"Temps"] <- paste(incubation[,"DATE_MESURE"],
incubation[,"HEURE_MESURE"])
incubation[,"Temps"] <- as.POSIXct(strptime(incubation[,"Temps"],
format="%d/%m/%Y %H:%M:%S"), tz="UTC")

# Perform a "cast" to group the data by certain columns
incubation_cast_oxy <- cast(incubation, ID_POTS + EXP + SOUS_PERIODE +
ESPECE + NB_PIEDS + DATE_MESURE ~ CONDITIONS, value = "OXY_mg.L")
incubation_cast_heure <- cast(incubation, ID_POTS + EXP + SOUS_PERIODE +
ESPECE + NB_PIEDS + DATE_MESURE ~ CONDITIONS, value = "Temps")

incubation_cast_heure <- pivot_wider(incubation,
                                     id_cols = c(ID_POTS, EXP,
SOUS_PERIODE, ESPECE, NB_PIEDS, DATE_MESURE),
                                     names_from = CONDITIONS,
                                     values_from = Temps)

# Calculate time differences between the "DAY" and "NIGHT" conditions
incubation_cast_heure[,"diff_temps_jour"] <-
(incubation_cast_heure[,"JOUR 2"] - incubation_cast_heure[,"JOUR 1"]) /
60
incubation_cast_heure[,"diff_temps_nuit"] <-
(incubation_cast_heure[,"NUIT 2"] - incubation_cast_heure[,"NUIT 1"]) /
60
```

```
# Calculate oxygen differences between the "DAY" and "NIGHT" conditions
incubation_cast_oxy[,"diff_oxy_jour"] <- (incubation_cast_oxy[,"JOUR 2"]
- incubation_cast_oxy[,"JOUR 1"])
incubation_cast_oxy[,"diff_oxy_nuit"] <- (incubation_cast_oxy[,"NUIT 2"]
- incubation_cast_oxy[,"NUIT 1"])

# Create a vector containing identifiers for future merging
identifiant <- c("ID_POTS", "EXP", "DATE_MESURE", "SOUS_PERIODE",
"NB_PIEDS", "ESPECE")


# Merge the data based on identifiers
incubation_final <- merge(incubation_cast_heure[, c(identifiant,
"diff_temps_jour", "diff_temps_nuit")],
                          incubation_cast_oxy[, c(identifiant,
"diff_oxy_jour", "diff_oxy_nuit")])

# Convert the difftime columns to numeric
incubation_final$diff_temps_jour <-
as.numeric(incubation_final$diff_temps_jour)
incubation_final$diff_oxy_jour <-
as.numeric(incubation_final$diff_oxy_jour)
incubation_final$diff_temps_nuit <-
as.numeric(incubation_final$diff_temps_nuit)
incubation_final$diff_oxy_nuit <-
as.numeric(incubation_final$diff_oxy_nuit)

# Calculate net production and respiration rates in mg O2 / cm2 / hour
incubation_final[,"Production_net"] <-
(incubation_final[,"diff_oxy_jour"] /
incubation_final[,"diff_temps_jour"])*2 #divide by the 2-liter volume
incubation_final[,"Respiration"] <- (incubation_final[,"diff_oxy_nuit"] /
incubation_final[,"diff_temps_nuit"] )*2

# Calculate gross production by taking the absolute value of net
production and respiration rates
incubation_final[,"Production_brute"] <-
abs(incubation_final[,"Production_net"]) +
abs(incubation_final[,"Respiration"])

# Replace positive values in the "Respiration" column with zero in a new
column "Respiration_corr"
incubation_final[,"Respiration_corr"] <- incubation_final[,"Respiration"]

# Find the indices of positive values in the "Respiration_corr" column
indice_positive <- which(incubation_final[,"Respiration_corr"] > 0)

# Replace positive values with zero in the "Respiration_corr" column
incubation_final[indice_positive,"Respiration_corr"] <- 0

# Calculate corrected gross production using the corrected
"Respiration_corr" values
incubation_final[,"Production_brute_corr"] <-
abs(incubation_final[,"Production_net"]) +
abs(incubation_final[,"Respiration_corr"])

# Now I will merge the two tables to change the unit, so df_pots and
incubation final
```

````
# Rename column EXP
colnames(incubation_final)[which(colnames(incubation_final)== "EXP")] <-
"EXPERIENCE"
colnames(incubation_final)[which(colnames(incubation_final)==
"SOUS_PERIODE")] <- "TEMPS_MESURE"

# Replace "T" with "P" in the "TEMPS_MESURE" column
incubation_final$TEMPS_MESURE <- gsub('P', 'T',
incubation_final$TEMPS_MESURE)



merge_data <- merge(incubation_final, df_pots, by = c("ESPECE",
"TEMPS_MESURE", "EXPERIENCE","ID_POTS"))

work_data <- merge_data[c("ESPECE",
"TEMPS_MESURE","EXPERIENCE","ID_POTS","DATE_MESURE","NB_PIEDS","Productio
n_net","Respiration_corr","Production_brute_corr","SF_POT_CM2","n_pieds")
]

resultats_moyens <- work_data %>%
  group_by(ESPECE, TEMPS_MESURE, EXPERIENCE,ID_POTS) %>%
  summarize(Moyenne_SF_pieds = mean(SF_POT_CM2 / n_pieds, na.rm = TRUE))


# Merge the average results with the original dataframe
work_data_bis_printemps <- merge(work_data, resultats_moyens,
                       by = c("ESPECE", "TEMPS_MESURE", "EXPERIENCE",
"ID_POTS"))


# Calculate the difference between NB_PIEDS and n_pieds
work_data_bis_printemps$Difference_Pieds <-
work_data_bis_printemps$NB_PIEDS - work_data_bis_printemps$n_pieds

# Add the average SF_pieds for pots with a difference in feet
work_data_bis_printemps <- work_data_bis_printemps %>%
  mutate(SF_POT_CM2_Corrigee = SF_POT_CM2 + Difference_Pieds *
Moyenne_SF_pieds)

# Now I want to correct the values of NCP, GCP, and CR based on the
photosynthetic area
work_data_bis_printemps[,"Production_net_area"] <-
work_data_bis_printemps[,"Production_net"] /
work_data_bis_printemps[,"SF_POT_CM2_Corrigee"]
work_data_bis_printemps[,"Respiration_area"] <-
work_data_bis_printemps[,"Respiration_corr"] /
work_data_bis_printemps[,"SF_POT_CM2_Corrigee"]
work_data_bis_printemps[,"Production_brute_area"] <-
work_data_bis_printemps[,"Production_brute_corr"] /
work_data_bis_printemps[,"SF_POT_CM2_Corrigee"]

```

#### Figure 3-4-5 (panel c & f)

```{r}
````

```
Resume_oxy_spring <- work_data_bis_printemps %>%
   filter(TEMPS_MESURE == "T2") %>%
  group_by(ESPECE, EXPERIENCE, TEMPS_MESURE) %>%
  dplyr::summarise(
    mean_prod = mean(Production_brute_area),
    sd_prod = sd(Production_brute_area),
    n_obs_prod = n(),
    se_prod = sd_prod / sqrt(n_obs_prod),
    mean_resp = mean(Respiration_area),
    sd_resp = sd(Respiration_area),
    n_obs_resp = n(),
    se_resp = sd_resp / sqrt(n_obs_resp)
  )

 Resume_oxy_summer <- work_data_bis %>%
   filter(TEMPS_MESURE == "T2") %>%
  group_by(ESPECE, EXPERIENCE, TEMPS_MESURE) %>%
  dplyr::summarise(
    mean_prod = mean(Production_brute_area),
    sd_prod = sd(Production_brute_area),
    n_obs_prod = n(),
    se_prod = sd_prod / sqrt(n_obs_prod),
    mean_resp = mean(Respiration_area),
    sd_resp = sd(Respiration_area),
    n_obs_resp = n(),
    se_resp = sd_resp / sqrt(n_obs_resp)
  )


# Perform a pivot of your table to overlay the graphs
pivot_data_O2_printemps <- work_data_bis_printemps %>%
  pivot_longer(cols = c(Production_brute_area, Respiration_area),
names_to = "Variable", values_to = "O2")

# Perform a pivot of your table to overlay the graphs
pivot_data_O2_ete <- work_data_bis %>%
  pivot_longer(cols = c(Production_brute_area, Respiration_area),
names_to = "Variable", values_to = "O2")


# Add a "season" column to each table
pivot_data_O2_printemps$saison <- "spring"
pivot_data_O2_ete$saison <- "summer"


# Convert DATE_MESURE to character type in both tables
pivot_data_O2_printemps <- pivot_data_O2_printemps %>%
  mutate(DATE_MESURE = as.character(DATE_MESURE))
pivot_data_O2_ete <- pivot_data_O2_ete %>%
  mutate(DATE_MESURE = as.character(DATE_MESURE))

# Remove TO from summer:
pivot_data_O2_ete <- pivot_data_O2_ete %>%
  filter(TEMPS_MESURE != "T0")

# Merge the two tables into one
O2_final <- bind_rows(pivot_data_O2_printemps, pivot_data_O2_ete)
```

```
couleur_valeurs <- c("#9ecae1", "#f77f58", "#a1d99b")
etiquettes <- c("Acclimation", "Heatwave", "Recovery")



 ### GRAPH FINAL ###
 metabo_CN <- O2_final %>%
  filter(ESPECE == "CN",
         TEMPS_MESURE == "T2")%>%
  ggplot(aes(x = factor(EXPERIENCE, levels = c("C", "2", "4", "6")), y =
O2, fill = TEMPS_MESURE)) +
  geom_boxplot(width = 0.25, show.legend = F)+
  geom_vline(xintercept = 1.5, linetype = "dashed", color = "gray", size
= 0.5) +
  geom_vline(xintercept = 2.5, linetype = "dashed", color = "gray", size
= 0.5) +
  geom_vline(xintercept = 3.5, linetype = "dashed", color = "gray", size
= 0.5) +
  stat_summary(fun = mean, geom = "point", color = "red", position =
position_dodge(width = 0.75), show.legend = F) +
  scale_fill_manual(values = "#f77f58", labels = "Heatwave") +
  scale_x_discrete(breaks = c("C", "2", "4", "6"),
  labels = c("Control", "+2", "+4", "+6"))+
     scale_y_continuous(labels = scales::comma)+
  theme_bw() +
   facet_grid(Variable ~ saison, scales = "free_y",
              labeller = labeller(saison = c("spring" = "Spring",
"summer" = "Summer"),
                                  Variable = c("Production_brute_area" =
"Gross oxygen prod", "Respiration_area" = "Respiration"))) +
labs(y = expression(O[2]~(mg~mm^2~h^-1)),
     x = "Experimental conditions",
        fill = " ") +
   theme(strip.text.x = element_blank(),
         strip.text.y = element_text(size = 15),
         strip.background = element_rect(fill = "white"),
        axis.text.x = element_text(size = 18),
        axis.text.y = element_text(size = 18),
        axis.title.y = element_text(size = 20),
        axis.title.x = element_text(size = 20))+
    # Ajouter des annotations spécifiques à chaque facette
  geom_text(data = . %>% filter(Variable == "Production_brute_area"),
            aes(x = 0.45, y = 0.0020,
                label = "Kruskal-Wallis = ns"),
            inherit.aes = FALSE, size = 6, color = "#4D4D4D", hjust = 0)
+
  geom_text(data = . %>% filter(Variable == "Respiration_area"),
            aes(x = 0.45, y = -0.0004,
                label = "Kruskal-Wallis = ns"),
            inherit.aes = FALSE, size = 6,color = "#4D4D4D", hjust = 0)



 metabo_ZN  <- O2_final %>%
  filter(ESPECE == "ZN",
         TEMPS_MESURE == "T2")%>%
```

```
  ggplot(aes(x = factor(EXPERIENCE, levels = c("C", "2", "4", "6")), y =
O2, fill = TEMPS_MESURE)) +
  geom_boxplot(width = 0.25, show.legend = F)+
  geom_vline(xintercept = 1.5, linetype = "dashed", color = "gray", size
= 0.5) +
  geom_vline(xintercept = 2.5, linetype = "dashed", color = "gray", size
= 0.5) +
  geom_vline(xintercept = 3.5, linetype = "dashed", color = "gray", size
= 0.5) +
  stat_summary(fun = mean, geom = "point", color = "red", position =
position_dodge(width = 0.75), show.legend = F) +
  scale_fill_manual(values = "#f77f58", labels = "Heatwave") +
  scale_x_discrete(breaks = c("C", "2", "4", "6"),
  labels = c("Control", "+2", "+4", "+6"))+
   scale_y_continuous(labels = scales::comma)+
  theme_bw() +
   facet_grid(Variable ~ saison, scales = "free_y",
              labeller = labeller(saison = c("spring" = "Spring",
"summer" = "Summer"),
                                  Variable = c("Production_brute_area" =
"Gross oxygen prod", "Respiration_area" = "Respiration"))) +
labs(y = expression(O[2]~(mg~mm^2~h^-1)),
     x = "Experimental conditions",
        fill = " ") +
   theme(strip.text.x = element_blank(),
         strip.text.y = element_text(size = 15),
         strip.background = element_rect(fill = "white"),
        axis.text.x = element_text(size = 18),
        axis.text.y = element_text(size = 18),
        axis.title.y = element_text(size = 20),
        axis.title.x = element_text(size = 20))+
  geom_text(data = . %>% filter(Variable == "Production_brute_area"),
            aes(x = 0.45, y = 0.0025,
                label = "Kruskal-Wallis = ns"),
            inherit.aes = FALSE, size = 6, color = "#4D4D4D", hjust = 0)
+
  geom_text(data = . %>% filter(Variable == "Respiration_area"),
            aes(x = 0.45, y = -0.0004,
                label = "Kruskal-Wallis = ns"),
            inherit.aes = FALSE, size = 6,color = "#4D4D4D", hjust = 0)

  metabo_RC  <-
    O2_final %>%
  filter(ESPECE == "RC",
         TEMPS_MESURE == "T2")%>%
  ggplot(aes(x = factor(EXPERIENCE, levels = c("C", "2", "4", "6")), y =
O2, fill = TEMPS_MESURE)) +
  geom_boxplot(width = 0.25, show.legend = F)+
  geom_vline(xintercept = 1.5, linetype = "dashed", color = "gray", size
= 0.5) +
  geom_vline(xintercept = 2.5, linetype = "dashed", color = "gray", size
= 0.5) +
  geom_vline(xintercept = 3.5, linetype = "dashed", color = "gray", size
= 0.5) +
  stat_summary(fun = mean, geom = "point", color = "red", position =
position_dodge(width = 0.75), show.legend = F) +
  scale_fill_manual(values = "#f77f58", labels = "Heatwave") +
  scale_x_discrete(breaks = c("C", "2", "4", "6"),
```

```
  labels = c("Control", "+2", "+4", "+6"))+
  scale_y_continuous(labels = scales::comma)+
  theme_bw() +
   facet_grid(Variable ~ saison, scales = "free_y",
              labeller = labeller(saison = c("spring" = "Spring",
"summer" = "Summer"),
                                  Variable = c("Production_brute_area" =
"Gross oxygen prod", "Respiration_area" = "Respiration"))) +
labs(y = expression(O[2]~(mg~mm^2~h^-1)),
     x = "Experimental conditions",
        fill = " ") +
   theme(strip.text.x = element_blank(),
         strip.text.y = element_text(size = 15),
         strip.background = element_rect(fill = "white"),
        axis.text.x = element_text(size = 18),
        axis.text.y = element_text(size = 18),
        axis.title.y = element_text(size = 20),
        axis.title.x = element_text(size = 20)) +
  geom_segment(data = . %>% filter(Variable == "Production_brute_area",
saison == "summer"),
               aes(x = 1, xend = 2, y = 0.0035, yend = 0.0035), col =
"black") + # trait horizontal
  geom_segment(data = . %>% filter(Variable == "Production_brute_area",
saison == "summer"),
               aes(x = 1, xend = 1, y = 0.0035, yend = 0.0030), col =
"black") + # petit trait vertical à gauche
  geom_segment(data = . %>% filter(Variable == "Production_brute_area",
saison == "summer"),
               aes(x = 2, xend = 2, y = 0.0035, yend = 0.0030), col =
"black") +  # petit trait vertical à droite
geom_label(data = . %>% filter(Variable == "Production_brute_area",
saison == "summer"),
          aes(x = 1.5, y = 0.0035, label = "**"),
          fill = "white", color = "black", size = 5, label.size = 0)+
    # Ajouter des annotations spécifiques à chaque facette
  geom_text(data = . %>% filter(Variable == "Production_brute_area",
saison == "spring"),
            aes(x = 0.45, y = 0.0045,
                label = "Kruskal-Wallis : ns"),
            inherit.aes = FALSE, size = 6, color = "#4D4D4D", hjust = 0)
+
  geom_text(data = . %>% filter(Variable == "Production_brute_area",
saison == "summer"),
            aes(x = 0.45, y = 0.0045,
                label = "Kruskal-Wallis : p < 0.05"),
            inherit.aes = FALSE, size = 6,color = "#4D4D4D", hjust = 0) +
  geom_text(data = . %>% filter(Variable == "Respiration_area"),
            aes(x = 0.45, y = -0.00085,
                label = "Kruskal-Wallis : ns"),
            inherit.aes = FALSE, size = 6,color = "#4D4D4D", hjust = 0)

```

#### Statistical analysis GCP and CR - Spring and Summer

```{r}
```

```
##########################################################################
#######
##################################      CN
####################################
##########################################################################
#######


### GCP SPRING
spring_gross_CN <- O2_final %>%
  filter(saison == "spring", Variable == "Production_brute_area", ESPECE
== "CN", TEMPS_MESURE == "T2")

# Perform ANOVA for Gross Oxygen Production in Spring
anova_spring_gross_CN <- aov(O2 ~ EXPERIENCE, data = spring_gross_CN)
summary(anova_spring_gross_CN)

# Perform Shapiro-Wilk test on residuals (normality)
shapiro_gross_CN <- shapiro.test(residuals(anova_spring_gross_CN))
print(shapiro_gross_CN) #NON

# Perform Levene's test for homogeneity of variances
levene_gross_CN <- leveneTest(O2 ~ EXPERIENCE, data = spring_gross_CN)
print(levene_gross_CN) # OK


kruskal_gross_CN <- kruskal.test(O2 ~ EXPERIENCE, data = spring_gross_CN)
print(kruskal_gross_CN)

# Extract results into a data frame
results_kruskal_gross_CN <- data.frame(
  Test = "Kruskal-Wallis",
  Statistic = kruskal_gross_CN$statistic,
  P_Value = kruskal_gross_CN$p.value,
  Df = kruskal_gross_CN$parameter
)


### CR SPRING
spring_resp_CN <- O2_final %>%
  filter(saison == "spring", Variable == "Respiration_area", ESPECE ==
"CN",TEMPS_MESURE == "T2")

# Perform ANOVA for Respiration
anova_spring_resp_CN <- aov(O2 ~ EXPERIENCE, data = spring_resp_CN)
anova_summary <- summary(anova_spring_resp_CN)

shapiro_resp_CN <- shapiro.test(residuals(anova_spring_resp_CN))
print(shapiro_resp_CN) # OK

levene_resp_CN <- leveneTest(O2 ~ EXPERIENCE, data = spring_resp_CN)
print(levene_resp_CN) # OK

kruskal_resp_CN <- kruskal.test(O2 ~ EXPERIENCE, data = spring_resp_CN)
print(kruskal_gross_CN)

# Extract results into a data frame
results_kruskal_resp_CN <- data.frame(
```

```
  Test = "Kruskal-Wallis",
  Statistic = kruskal_resp_CN$statistic,
  P_Value = kruskal_resp_CN$p.value,
  Df = kruskal_resp_CN$parameter
)


## GCP SUMMER

summer_gross_CN <- O2_final %>%
  filter(saison == "summer", Variable == "Production_brute_area", ESPECE
== "CN", TEMPS_MESURE == "T2")

# Perform ANOVA for Gross Oxygen Production in summer
anova_summer_gross_CN <- aov(O2 ~ EXPERIENCE, data = summer_gross_CN)
anova_summary_summer <- summary(anova_summer_gross_CN)

shapiro_gross_CN_summer <- shapiro.test(residuals(anova_summer_gross_CN))
print(shapiro_gross_CN_summer)

levene_gross_CN_summer <- leveneTest(O2 ~ EXPERIENCE, data =
summer_gross_CN)
print(levene_gross_CN_summer)

kruskal_gross_CN_summer <- kruskal.test(O2 ~ EXPERIENCE, data =
summer_gross_CN)
print(kruskal_gross_CN_summer)

# Extract results into a data frame
results_kruskal_gross_CN_summer <- data.frame(
  Test = "Kruskal-Wallis",
  Statistic = kruskal_gross_CN_summer$statistic,
  P_Value = kruskal_gross_CN_summer$p.value,
  Df = kruskal_gross_CN_summer$parameter
)



## CR SUMMER
summer_resp_CN <- O2_final %>%
  filter(saison == "summer", Variable == "Respiration_area", ESPECE ==
"CN",TEMPS_MESURE == "T2")

# Perform ANOVA for Gross Oxygen Production in summer
anova_summer_resp_CN <- aov(O2 ~ EXPERIENCE, data = summer_resp_CN)
anova_summary_summer <- summary(anova_summer_resp_CN)

shapiro_resp_CN_summer <- shapiro.test(residuals(anova_summer_resp_CN))
print(shapiro_resp_CN_summer)

levene_resp_CN_summer <- leveneTest(O2 ~ EXPERIENCE, data =
summer_resp_CN)
print(levene_resp_CN_summer)

kruskal_resp_CN_summer <- kruskal.test(O2 ~ EXPERIENCE, data =
summer_resp_CN)
print(kruskal_resp_CN_summer)
```

```
# Extract results into a data frame
results_kruskal_resp_CN_summer <- data.frame(
  Test = "Kruskal-Wallis",
  Statistic = kruskal_resp_CN_summer$statistic,
  P_Value = kruskal_resp_CN_summer$p.value,
  Df = kruskal_resp_CN_summer$parameter
)



##########################################################################
#######
##################################     ZN
###################################
##########################################################################
#######

## GCP SPRING
spring_gross_ZN <- O2_final %>%
  filter(saison == "spring", Variable == "Production_brute_area", ESPECE
== "ZN", TEMPS_MESURE == "T2")

# Perform ANOVA for Gross Oxygen Production in Spring
anova_spring_gross_ZN <- aov(O2 ~ EXPERIENCE, data = spring_gross_ZN)
summary(anova_spring_gross_ZN)

shapiro_gross_ZN <- shapiro.test(residuals(anova_spring_gross_ZN))
print(shapiro_gross_ZN) # NON

kruskal_gross_ZN <- kruskal.test(O2 ~ EXPERIENCE, data = spring_gross_ZN)
print(kruskal_gross_ZN)

# Extract results into a data frame
results_kruskal_gross_ZN <- data.frame(
  Test = "Kruskal-Wallis",
  Statistic = kruskal_gross_ZN$statistic,
  P_Value = kruskal_gross_ZN$p.value,
  Df = kruskal_gross_ZN$parameter
)


## CR SPRING
spring_resp_ZN <- O2_final %>%
  filter(saison == "spring", Variable == "Respiration_area", ESPECE ==
"ZN",TEMPS_MESURE == "T2")

# Perform ANOVA for Respiration
anova_spring_resp_ZN <- aov(O2 ~ EXPERIENCE, data = spring_resp_ZN)
summary(anova_spring_resp_ZN)

shapiro_resp_ZN_spring <- shapiro.test(residuals(anova_spring_resp_ZN))
print(shapiro_resp_ZN_spring) # NON

levene_resp_ZN_spring <- leveneTest(O2 ~ EXPERIENCE, data =
spring_resp_ZN)
print(levene_resp_ZN_spring) # OK
```

```
kruskal_resp_ZN <- kruskal.test(O2 ~ EXPERIENCE, data = spring_resp_ZN)
print(kruskal_resp_ZN)

# Extract results into a data frame
results_kruskal_resp_ZN <- data.frame(
  Test = "Kruskal-Wallis",
  Statistic = kruskal_resp_ZN$statistic,
  P_Value = kruskal_resp_ZN$p.value,
  Df = kruskal_resp_ZN$parameter
)


## GCP SUMMER
summer_gross_ZN <- O2_final %>%
  filter(saison == "summer", Variable == "Production_brute_area", ESPECE
== "ZN", TEMPS_MESURE == "T2")

# Perform ANOVA for Gross Oxygen Production in summer
anova_summer_gross_ZN <- aov(O2 ~ EXPERIENCE, data = summer_gross_ZN)
anova_summary_summer<- summary(anova_summer_gross_ZN)

shapiro_gross_ZN_summer <- shapiro.test(residuals(anova_summer_gross_ZN))
print(shapiro_gross_ZN_summer) # OK

levene_gross_ZN_summer <- leveneTest(O2 ~ EXPERIENCE, data =
summer_gross_ZN)
print(levene_gross_ZN_summer) # OK


kruskal_gross_ZN_summer <- kruskal.test(O2 ~ EXPERIENCE, data =
summer_gross_ZN)
print(kruskal_gross_ZN_summer)

# Extract results into a data frame
results_kruskal_gross_ZN_summer <- data.frame(
  Test = "Kruskal-Wallis",
  Statistic = kruskal_gross_ZN_summer$statistic,
  P_Value = kruskal_gross_ZN_summer$p.value,
  Df = kruskal_gross_ZN_summer$parameter)

## CR SUMMER
summer_resp_ZN <- O2_final %>%
  filter(saison == "summer", Variable == "Respiration_area", ESPECE ==
"ZN",TEMPS_MESURE == "T2")

# Perform ANOVA for Gross Oxygen Production in summer
anova_summer_resp_ZN <- aov(O2 ~ EXPERIENCE, data = summer_resp_ZN)
anova_summary_summer_resp <- summary(anova_summer_resp_ZN)

shapiro_resp_ZN_summer <- shapiro.test(residuals(anova_summer_resp_ZN))
print(shapiro_resp_ZN_summer) # OK

levene_resp_ZN_summer <- leveneTest(O2 ~ EXPERIENCE, data =
summer_resp_ZN)
print(levene_resp_ZN_summer) # OK

kruskal_resp_ZN_summer <- kruskal.test(O2 ~ EXPERIENCE, data =
summer_resp_ZN)
```

```
print(kruskal_resp_ZN_summer)

# Extract results into a data frame
results_kruskal_resp_ZN_summer <- data.frame(
  Test = "Kruskal-Wallis",
  Statistic = kruskal_resp_ZN_summer$statistic,
  P_Value = kruskal_resp_ZN_summer$p.value,
  Df = kruskal_resp_ZN_summer$parameter
)



#########################################################################
#######
################################     RC
##################################
#########################################################################
#######

## GCP SPRING

spring_gross_RC <- O2_final %>%
  filter(saison == "spring", Variable == "Production_brute_area", ESPECE
== "RC", TEMPS_MESURE == "T2")

# Perform ANOVA for Gross Oxygen Production in Spring
anova_spring_gross_RC <- aov(O2 ~ EXPERIENCE, data = spring_gross_RC)
anova_summary_RC_gross <- summary(anova_spring_gross_RC)

shapiro_gross_RC_spring <- shapiro.test(residuals(anova_spring_gross_RC))
print(shapiro_gross_RC_spring)

levene_gross_RC_spring <- leveneTest(O2 ~ EXPERIENCE, data =
spring_gross_RC)
print(levene_gross_RC_spring)

kruskal_gross_RC_spring <- kruskal.test(O2 ~ EXPERIENCE, data =
spring_gross_RC)
print(kruskal_gross_RC_spring)

# Extract results into a data frame
results_kruskal_gross_RC_spring <- data.frame(
  Test = "Kruskal-Wallis",
  Statistic = kruskal_gross_RC_spring$statistic,
  P_Value = kruskal_gross_RC_spring$p.value,
  Df = kruskal_gross_RC_spring$parameter
)


## CR SPRING
spring_resp_RC <- O2_final %>%
  filter(saison == "spring", Variable == "Respiration_area", ESPECE ==
"RC",TEMPS_MESURE == "T2")

# Perform ANOVA for Respiration
anova_spring_resp_RC <- aov(O2 ~ EXPERIENCE, data = spring_resp_RC)
summary(anova_spring_resp_RC)
```

```
shapiro_resp_RC_spring <- shapiro.test(residuals(anova_spring_resp_RC))
print(shapiro_resp_RC_spring) # NON

levene_resp_RC_spring <- leveneTest(O2 ~ EXPERIENCE, data =
spring_resp_RC)
print(levene_resp_RC_spring) # OK

kruskal_resp_RC <- kruskal.test(O2 ~ EXPERIENCE, data = spring_resp_RC)
print(kruskal_resp_RC)

# Extract results into a data frame
results_kruskal_resp_RC <- data.frame(
  Test = "Kruskal-Wallis",
  Statistic = kruskal_resp_RC$statistic,
  P_Value = kruskal_resp_RC$p.value,
  Df = kruskal_resp_RC$parameter
)


## GCP SUMMER
summer_gross_RC <- O2_final %>%
  filter(saison == "summer", Variable == "Production_brute_area", ESPECE
== "RC", TEMPS_MESURE == "T2")

# Perform ANOVA for Gross Oxygen Production in summer
anova_summer_gross_RC <- aov(O2 ~ EXPERIENCE, data = summer_gross_RC)
summary(anova_summer_gross_RC)

shapiro_gross_RC_summer <- shapiro.test(residuals(anova_summer_gross_RC))
print(shapiro_gross_RC_summer) # OK

levene_gross_RC_summer <- leveneTest(O2 ~ EXPERIENCE, data =
summer_gross_RC)
print(levene_gross_RC_summer) # NON

kruskal_gross_RC <- kruskal.test(O2 ~ EXPERIENCE, data = summer_gross_RC)
print(kruskal_gross_RC)

# Extract results into a data frame
results_kruskal_gross_RC <- data.frame(
  Test = "Kruskal-Wallis",
  Statistic = kruskal_gross_RC$statistic,
  P_Value = kruskal_gross_RC$p.value,
  Df = kruskal_gross_RC$parameter
)


library(dunn.test)
dunn_result <- dunn.test(summer_gross_RC$O2, summer_gross_RC$EXPERIENCE,
method = "bonferroni")
print(dunn_result)


## CR SUMMER
summer_resp_RC <- O2_final %>%
  filter(saison == "summer", Variable == "Respiration_area", ESPECE ==
"RC",TEMPS_MESURE == "T2")
```

```
# Perform ANOVA for Gross Oxygen Production in summer
anova_summer_resp_RC <- aov(O2 ~ EXPERIENCE, data = summer_resp_RC)
anova_summary_RC_resp <- summary(anova_summer_resp_RC)

shapiro_resp_RC_summer <- shapiro.test(residuals(anova_summer_resp_RC))
print(shapiro_resp_RC_summer)

levene_resp_RC_summer <- leveneTest(O2 ~ EXPERIENCE, data =
summer_resp_RC)
print(levene_resp_RC_summer)

kruskal_resp_RC_summer <- kruskal.test(O2 ~ EXPERIENCE, data =
summer_resp_RC)
print(kruskal_resp_RC_summer)

# Extract results into a data frame
results_kruskal_resp_RC_summer <- data.frame(
  Test = "Kruskal-Wallis",
  Statistic = kruskal_resp_RC_summer$statistic,
  P_Value = kruskal_resp_RC_summer$p.value,
  Df = kruskal_resp_RC_summer$parameter
)
```

##### EXPERIMENTAL CONDITIONS DATA

#### TEMPERATURE SPRING

```{r}
#LOAD DATA
# Donnees_temp <-  "~" # add your working directory here
setwd(Donnees_temp)

fichier <- "temperature_C2ZO_printemps.csv"
temperature <- read.table(fichier, sep = ";", h = TRUE, dec = ",")

## TREATMENT ##
colnames(temperature) <-
c("N","Date","contrôle","+2","+4","+6","SOUS_PERIODE")

  temperature[,"Temps"] <- as.POSIXct(strptime(temperature[,"Date"],
format="%m/%d/%y %I:%M:%S %p"),tz="UTC")
  temperature[,"DATE"] <- as.Date(temperature[,"Temps"])

  temperature <- temperature %>%
    pivot_longer(cols = c(contrôle, `+2`, `+4`, `+6`),
                 names_to = "Conditions",
                 values_to = "Temperature")

  temperature <- as.data.frame(temperature)

  temperature[,"Conditions"] <- as.factor(temperature[,"Conditions"])
  temperature[,"Temperature"] <- as.numeric(temperature[,"Temperature"])

## STATISTICAL ANALYSIS ##
moyenne_quotidienne <- aggregate(Temperature ~
DATE*Conditions*SOUS_PERIODE, data = temperature, FUN = mean)
```

````
moyenne_quotidienne_P1 <- moyenne_quotidienne %>%
    filter(SOUS_PERIODE == "P1")

#Kruskall
kruskal_result_P1_spring <- kruskal.test(Temperature ~ Conditions, data =
moyenne_quotidienne_P1)
print(kruskal_result_P1_spring)
# Extract results into a data frame
results_kruskal_P1_spring <- data.frame(
  Test = "Kruskal-Wallis",
  Statistic = kruskal_result_P1_spring$statistic,
  P_Value = kruskal_result_P1_spring$p.value,
  Df = kruskal_result_P1_spring$parameter
)


#### P2 ####
moyenne_quotidienne_P2 <- moyenne_quotidienne %>%
    filter(SOUS_PERIODE == "P2")

kruskal_result_P2_spring <- kruskal.test(Temperature ~ Conditions, data =
moyenne_quotidienne_P2)
print(kruskal_result_P2_spring)
# Extract results into a data frame
results_kruskal_P2_spring <- data.frame(
  Test = "Kruskal-Wallis",
  Statistic = kruskal_result_P2_spring$statistic,
  P_Value = kruskal_result_P2_spring$p.value,
  Df = kruskal_result_P2_spring$parameter
)

library(dunn.test)
dunn_result_P2_spring <- dunn.test(moyenne_quotidienne_P2$Temperature,
moyenne_quotidienne_P2$Conditions, method ="BH")
print(dunn_result_P2_spring)

#### P3 ####
moyenne_quotidienne_P3 <- moyenne_quotidienne %>%
    filter(SOUS_PERIODE == "P3")

kruskal_result_P3_spring <- kruskal.test(Temperature ~ Conditions, data =
moyenne_quotidienne_P3)
print(kruskal_result_P3_spring)
# Extract results into a data frame
results_kruskal_P3_spring <- data.frame(
  Test = "Kruskal-Wallis",
  Statistic = kruskal_result_P3_spring$statistic,
  P_Value = kruskal_result_P3_spring$p.value,
  Df = kruskal_result_P3_spring$parameter
)
```

#### Figure 2 - left (spring)

```{r}

temperature %>%
````

```
    ggplot(aes(x = factor(Conditions, levels =  c("contrôle", "+2", "+4",
"+6")), y = Temperature, fill = SOUS_PERIODE))+
    geom_boxplot()+
    geom_vline(xintercept = 1.5, linetype="dashed",
               color = "gray", size=0.5)+
    geom_vline(xintercept = 2.5, linetype="dashed",
               color = "gray", size=0.5)+
    geom_vline(xintercept = 3.5, linetype="dashed",
               color = "gray", size=0.5)+
    stat_summary(fun=mean,
                 geom="point",
                 color = "red",
                 position=position_dodge(width=0.75))+
 scale_fill_manual(labels = c("Before", "Heatwave", "After"),
  values = c("#9ecae1", "#f77f58", "#a1d99b")) +
  scale_x_discrete(breaks = c("contrôle", "+2", "+4", "+6"),
  labels = c("Control", "+2", "+4", "+6")) +
    theme_bw()+
    labs(y = "Temperature (°C)",
         x = "Experimental conditions",
         fill = " ")+
    theme(strip.text.x = element_text(size = 20),
          axis.text.x = element_text(size = 18),
          axis.text.y = element_text(size = 18),
          axis.title.y = element_text(size = 20),
          axis.title.x = element_text(size = 20),
          legend.text = element_text(size = 18))+
    geom_segment(x=1, xend=2, y=35, yend=35, col="black") + # trait
horizontal
    geom_segment(x=1, xend=1, y=35, yend=34, col="black") + # petit trait
verticla à gauche
    geom_segment(x=2, xend=2, y=35, yend=34, col="black") + # petit trait
vertical à droite
    annotate("label", x = 1.5, y = 36.5, label = "*", color = "black",
label.size = NA, size = 8)+
    geom_segment(x=1, xend=3, y=38, yend=38, col="black") +
    geom_segment(x=1, xend=1, y=38, yend=37, col="black") +
    geom_segment(x=3, xend=3, y=38, yend=37, col="black") +
  annotate("label", x = 2, y = 39.5, label = "***", color = "black",
label.size = NA, size = 8)+
    geom_segment(x=1, xend=4, y=41, yend=41, col="black") +
    geom_segment(x=1, xend=1, y=41, yend=40, col="black") +
    geom_segment(x=4, xend=4, y=41, yend=40, col="black") +
  annotate("label", x = 2.5, y = 42.5, label = "***", color = "black",
label.size = NA, size = 8
          )+
    ylim(13, 50)

```

#### TEMPERATURE SUMMER

```{r}
#Load data
setwd(Donnees_temp)
# Charger les données
fichier <- "temperature_C2ZO_ete.csv"
temperature_ete <- read.table(fichier, sep = ";", h = TRUE, dec = ",")
```

```
# Treatment
colnames(temperature_ete) <-
c("N","Date","contrôle","+2","+4","+6","SOUS_PERIODE")

temperature_ete[,"Temps"] <-
as.POSIXct(strptime(temperature_ete[,"Date"], format="%m/%d/%y %I:%M:%S
%p"),tz="UTC")
temperature_ete[,"DATE"] <- as.Date(temperature_ete[,"Temps"])

temperature_ete <- temperature_ete %>%
    pivot_longer(cols = c(contrôle, `+2`, `+4`, `+6`),
                 names_to = "Conditions",
                 values_to = "Temperature")

temperature_ete <- as.data.frame(temperature_ete)

temperature_ete[,"Conditions"] <-
as.factor(temperature_ete[,"Conditions"])
temperature_ete[,"Temperature"] <-
as.numeric(temperature_ete[,"Temperature"])


## STATISTICAL ANALYSIS ##

moyenne_quotidienne_ete <- aggregate(Temperature ~
DATE*Conditions*SOUS_PERIODE, data = temperature_ete, FUN = mean)
moyenne_quotidienne_ete_P1 <- moyenne_quotidienne_ete %>%
    filter(SOUS_PERIODE == "P1")

kruskal_result_P1_summer <- kruskal.test(Temperature ~ Conditions, data =
moyenne_quotidienne_ete_P1)
print(kruskal_result_P1_summer)
# Extract results into a data frame
results_kruskal_P1_summer <- data.frame(
  Test = "Kruskal-Wallis",
  Statistic = kruskal_result_P1_summer$statistic,
  P_Value = kruskal_result_P1_summer$p.value,
  Df = kruskal_result_P1_summer$parameter
)

#### P2 ####
moyenne_quotidienne_ete_P2 <- moyenne_quotidienne_ete %>%
    filter(SOUS_PERIODE == "P2")

kruskal_result_P2_summer <- kruskal.test(Temperature ~ Conditions, data =
moyenne_quotidienne_ete_P2)
print(kruskal_result_P2_summer)
# Extract results into a data frame
results_kruskal_P2_summer <- data.frame(
  Test = "Kruskal-Wallis",
  Statistic = kruskal_result_P2_summer$statistic,
  P_Value = kruskal_result_P2_summer$p.value,
  Df = kruskal_result_P2_summer$parameter
)

library(dunn.test)
```

````
dunn_result_P2_summer <-
dunn.test(moyenne_quotidienne_ete_P2$Temperature,
moyenne_quotidienne_ete_P2$Conditions, method ="BH")
print(dunn_result_P2_summer)

#### P3 ####
moyenne_quotidienne_ete_P3 <- moyenne_quotidienne_ete %>%
    filter(SOUS_PERIODE == "P3")

kruskal_result_P3_summer <- kruskal.test(Temperature ~ Conditions, data =
moyenne_quotidienne_ete_P3)
print(kruskal_result_P3_summer)

results_kruskal_P3_summer <- data.frame(
  Test = "Kruskal-Wallis",
  Statistic = kruskal_result_P3_summer$statistic,
  P_Value = kruskal_result_P3_summer$p.value,
  Df = kruskal_result_P3_summer$parameter
)
```

#### Figure 2 - right (summer)

```{r}
temperature_ete %>%
    ggplot(aes(x = factor(Conditions, levels =  c("contrôle", "+2", "+4",
"+6")), y = Temperature, fill = SOUS_PERIODE))+
    geom_boxplot()+
    geom_vline(xintercept = 1.5, linetype="dashed",
               color = "gray", size=0.5)+
    geom_vline(xintercept = 2.5, linetype="dashed",
               color = "gray", size=0.5)+
    geom_vline(xintercept = 3.5, linetype="dashed",
               color = "gray", size=0.5)+
    stat_summary(fun=mean,
                 geom="point",
                 color = "red",
                 position=position_dodge(width=0.75))+
 scale_fill_manual(labels = c("Before", "Heatwave", "After"),
  values = c("#9ecae1", "#f77f58", "#a1d99b")) +
  scale_x_discrete(breaks = c("contrôle", "+2", "+4", "+6"),
  labels = c("Control", "+2", "+4", "+6")) +
    theme_bw()+
    labs(y = "Temperature (°C)",
         x = "Experimental conditions",
         fill = " ")+
    theme(strip.text.x = element_text(size = 20),
          axis.text.x = element_text(size = 18),
          axis.text.y = element_text(size = 18),
          axis.title.y = element_text(size = 20),
          axis.title.x = element_text(size = 20),
          legend.text = element_text(size = 18))+
    geom_segment(x=1, xend=2, y=41, yend=41, col="black") + # trait
horizontal
    geom_segment(x=1, xend=1, y=41, yend=40, col="black") + # petit trait
verticla à gauche
    geom_segment(x=2, xend=2, y=41, yend=40, col="black") + # petit trait
vertical à droite
````

```
    annotate("label", x = 1.5, y = 42.5, label = "●", color = "black", 
label.size = NA, size = 5)+
    geom_segment(x=1, xend=3, y=45, yend=45, col="black") +
    geom_segment(x=1, xend=1, y=45, yend=44, col="black") +
    geom_segment(x=3, xend=3, y=45, yend=44, col="black") +
  annotate("label", x = 2, y = 46.5, label = "***", color = "black", 
label.size = NA, size = 8)+
    geom_segment(x=1, xend=4, y=48, yend=48, col="black") +
    geom_segment(x=1, xend=1, y=48, yend=47, col="black") +
    geom_segment(x=4, xend=4, y=48, yend=47, col="black") +
  annotate("label", x = 2.5, y = 49.5, label = "***", color = "black", 
label.size = NA, size = 8
          )+
    ylim(13, 50)
```
